# Scalable, Generalizable, and Uncertainty-Aware Integration of Spatial Multi-Omics Across Diverse Modalities and Platforms with SCIGMA

**DOI:** 10.64898/2026.04.19.718223

**Authors:** Seowon Chang, Alexander Fleischmann, Ying Ma

## Abstract

Recent advances in spatial omics technologies have enabled simultaneous profiling of transcriptomic, proteomic, epigenomic, metabolomic, and imaging data at high spatial resolution, offering unprecedented opportunities to dissect tissue complexity. However, integrating these diverse and large-scale spatial multi-modal datasets remains a major computational challenge. We present SCIGMA, a scalable and generalizable deep learning framework for spatial multiomics integration. SCIGMA introduces a novel uncertainty-aware contrastive learning objective and multi-view graph neural networks to preserve modality-specific signals while learning biologically meaningful joint representations. Unlike existing methods, SCIGMA provides spatially resolved uncertainty estimates, improving interpretability and identifying regions of biological or technical heterogeneity. SCIGMA is the first spatial multi-omics method to support integration of up to five modalities—as demonstrated on Spatial-Mux-Seq data—and its modular framework is extensible to future technologies with even more modalities. It also scales to over one million spatial locations, enabling analysis of ultra-high-resolution datasets such as VisiumHD and Xenium Prime. We evaluated SCIGMA across 19 datasets spanning 8 modalities, 10 tissues, and 9 platforms. On benchmarkable datasets, SCIGMA outperformed existing methods in spatial domain detection, modality preservation, feature reconstruction, and reproducibility. Across applications, it uncovered biologically meaningful structures, refined spatial domains, and modality-specific regulatory programs, while its uncertainty estimates revealed tissue regions with potential biological or technical variation. Together, SCIGMA provides a robust, flexible, and future-ready solution for scalable spatial multi-modal integration.

## Introduction

Spatial transcriptomics (ST) has revolutionized gene expression analysis by preserving the spatial context of the transcriptome within tissues, offering critical insights into tissue architecture, intercellular communication, and disease mechanisms^1-7^. However, transcriptomic data alone provide an incomplete view of cellular function^8-13^, as they do not capture other molecular and structural features such as post-transcriptional modifications, chromatin accessibility, protein abundance and localization, or morphological context. To achieve a more comprehensive understanding of tissue biology, it is essential to integrate additional molecular layers, including proteomics, epigenomics, metabolic activity, and imaging-based morphology. For example, proteomic data reveal post-transcriptional modifications that influence protein function, while epigenomic data provide insights into the regulatory landscape governing gene expression^14,15^. Recent technological advances have expanded ST into spatial multi-modal platforms, enabling the simultaneous measurement of transcriptomic, proteomic, epigenomic, metabolomic, and imaging data from the same tissue. Emerging platforms now support the spatial co-profiling of multiple molecular modalities, including spatial-ATAC-RNA-seq (transcriptomics and chromatin accessibility)^16^, SPOTS^17^ and spatial-CITE-seq^18^ (transcriptomics and proteomics), 10x Xenium Prime and Visium HD^19^ (transcriptomics and imaging morphology), and spatial-Mux-seq^20^ (transcriptomics, proteomics, and epigenomics). Meanwhile, high-resolution ST technologies, such as 10x Xenium Prime and Visium HD, enable transcriptomic measurements at hundreds of thousands to millions of spatial locations, coupled with high-resolution morphological imaging. The recent development of Spatial-MUX-seq^20^ further extends the capabilities of spatial multi-modal omics by co-profiling up to five molecular modalities, offering a more comprehensive view of tissue complexity and cellular heterogeneity.

Despite the increasing availability of spatial multi-modal datasets, current computational approaches remain inadequate for fully leveraging their complexity. Existing multi-modal integration methods fall into two broad categories. The first category includes non-spatial multi-omics integration methods, such as Seurat^21^, TotalVI^22^, and MultiVI^23^, which integrate multimodal data at the single-cell level but fail to account for spatial organization. These methods primarily focus on aligning different molecular layers, such as gene expression and chromatin accessibility, for each cell without considering spatial dependencies across tissue architecture. As a result, their learned representations do not accurately capture spatially organized biological structures, an issue highlighted by multiple studies^24-26^. Without spatial context, these methods often misrepresent tissue domains, fail to detect spatially distinct cell types, and overlook critical interactions between neighboring cells. For example, Seurat^21^ uses canonical correlation analysis (CCA) and weighted nearest neighbor (WNN) matching to align multimodal data but lacks any spatial constraints, leading to poor preservation of tissue architecture. TotalVI and MultiVI are variational autoencoder (VAE)-based models that integrate single-cell multi-modal data, but it assumes cells are independent, preventing it from modeling spatially localized molecular programs. The second category consists of limited number of methods that are specifically designed for spatial multi-omics data, such as SpatialGlue^27^. SpatialGlue is a graph neural network (GNN)-based model with a dual-attention mechanism designed to combine molecular profiles with spatial structure. However, SpatialGlue faces several limitations that restrict its generalizability and scalability. It supports integration of at most three molecular modalities and does not incorporate imaging-derived features, limiting its compatibility with comprehensive platforms such as 10x Xenium, VisiumHD, and Spatial-MUX-seq. Its cross-modality correspondence loss, intended to align latent representations across modalities, can excessively enforce similarity between them, thereby reducing the model’s ability to preserve biologically meaningful, modality-specific variation. Additionally, the quadratic scaling of loss terms with the number of modalities introduces computational inefficiencies that hinder extension to higher-order integration. Finally, the absence of graph sampling renders the model unsuitable for large-scale datasets with millions of spatial locations, limiting its applicability to emerging high-resolution spatial transcriptomics technologies. Beyond these issues, many deep learning-based integration methods suffer from poor reproducibility across different runs due to stochastic training procedures and unstable optimization^28^. They often show limited generalization across spatial omics technologies, as they are designed and evaluated on specific datasets without robust adaptation mechanisms^26,29^. Additionally, these methods struggle to handle heterogeneous feature distributions, often arising from diverse molecular modalities such as transcriptomics, epigenomics, and proteomics, yet they assume modality homogeneity during integration by design. Another critical issue is their lack of interpretability, as most deep learning models do not quantify model uncertainty or yield interpretable latent spaces in the integration process, making it difficult to assess confidence in the learned representations and derive reliable biological insights.

To address these limitations, we present SCIGMA (uncertainty-aware **S**patially-informed **C**ontrastive learning-based **I**ntegration with **G**raph neural networks for spatial **M**ulti-modal data **A**nalysis), a scalable and interpretable deep learning framework for spatial multi-modal data integration. SCIGMA employs a novel contrastive learning framework that preserves modality-specific features while learning a cohesive joint representation, ensuring that distinct molecular signals contribute meaningfully to biological interpretation without artificially enforcing similarity. Unlike existing methods that either ignore spatial structure or struggle with heterogeneous multi-modal integration, SCIGMA leverages a multi-view graph that captures both spatial and feature-based similarities, enhancing its ability to dissect complex tissue organization. A unique feature of SCIGMA is its uncertainty-aware objective function, which quantifies uncertainty at each spatial location. This enables the identification of regions with high biological variability, distinguishes true biological heterogeneity from technical noise, and highlights modality-specific discrepancies. Additionally, SCIGMA is explicitly designed to scale efficiently to millions of spatial locations through graph sampling, making it applicable to high-resolution ST datasets such as Visium HD. Its flexible framework enables the seamless integration of up to five molecular modalities, as demonstrated with Spatial-MUX-seq data, and holds the potential to extend beyond five as spatial technologies advance. SCIGMA is accurate, scalable, interpretable, and robust, demonstrating its advantages across 19 datasets from 8 different modalities, covering 10 tissue types sequenced using 9 different technological platforms. Our results show that SCIGMA outperforms existing methods in spatial domain detection, achieving superior accuracy, enhanced reproducibility across multiple runs, recovering known biological structure, and uncovering novel biological insights. These include the delineation of refined tissue structures, the identification of novel marker genes, and the characterization of modality-specific contributions to tissue organization. With its adaptable design, SCIGMA supports the integration of all existing spatial multi-omics modalities, including imaging-based data, making it a versatile and scalable framework for spatial multi-omics analysis. Importantly, SCIGMA’s modeling framework is inherently flexible and robust, enabling seamless generalization to datasets with more than two modalities—including the latest five-modality spatial platform^20^—and positioning it to keep pace with the rapidly evolving landscape of spatial multi-omics technologies.

## Results

### SCIGMA Overview

The SCIGMA method is described in Method and Materials with a schematic in Figure 1 and additional technical details provided in Supplementary Note 1. Briefly, SCIGMA is a multi-view graph attention auto-encoder framework, enhanced by uncertainty-aware contrastive learning, to integrate spatial multi-modal data efficiently. It incorporates both spatial and feature similarity graphs, along with the corresponding feature matrices, as inputs. For each modality, a graph attention network (GAT) serves as the backbone, leveraging the feature matrices and the union of the two graph types to generate low-dimensional, modality-specific embeddings. Unlike existing methods that often focus on either spatial or feature similarities or handle them independently, SCIGMA constructs a unified graph that integrates both spatial and feature similarities, enabling the identification of both local and global tissue relationships. A cross-attention layer and contrastive learning objective integrates these modality-specific embeddings into a joint representation, ensuring that each modality retains its distinct features while preventing artificial homogeneity during training. To improve interpretability and robustness, we introduced a location-level uncertainty parameter within the contrastive learning framework, which not only stabilizes training but also provides precise measures of model uncertainty at each spatial location. For each modality, a GAT decoder reconstructs the original feature matrix, ensuring that the learned representation remains biologically meaningful. The resulting joint representation supports various downstream tasks such as domain detection and trajectory inference. Given the computational demands of graph neural networks, particularly for large datasets, SCIGMA employs an efficient graph sampling strategy to enhance scalability. This allows it to handle large-scale high-resolution datasets, such as 10x VisiumHD platform (e.g., at 2 *μm* resolution), which measures spatial locations over 1 million. Ablation studies evaluate six different settings to assess the necessity of each design component in SCIGMA: (1) removing spatial information, (2) omitting contrastive loss, (3) replacing SCIGMA’s novel contrastive loss with standard contrastive loss, (4) removing feature graphs, (5) replacing GAT with graph convolutional network (GCN), and (6) excluding uncertainty estimation, which affects model robustness and training consistency (Supplementary Figure 1, Supplementary Note 2). These analyses highlight the critical role of each design choice in preserving spatial structure, maintaining modality-specific signals, preventing over-smoothing of latent representations, and optimizing model accuracy, reproducibility, and interpretability.

**Figure 1.**
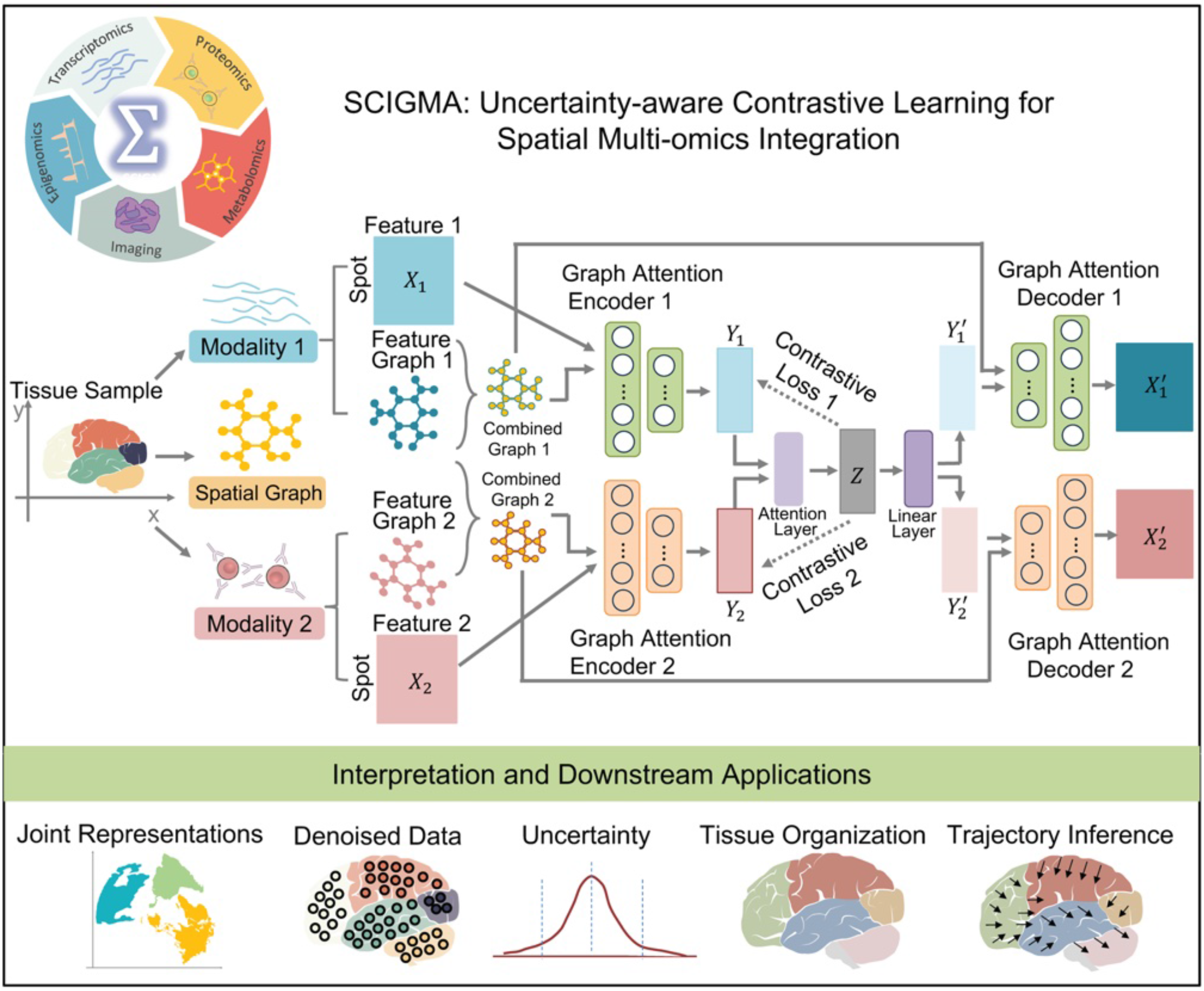
SCIGMA model workflow. SCIGMA is a scalable and interpretable graph neural network-based framework for integrating spatial multi-omics data across multiple modalities. SCIGMA requires spatial multi-omics data from the same tissue, including spatial coordinates and modality-specific features (e.g., transcriptomics, proteomics, imaging, and epigenomics). For each modality, SCIGMA constructs a combined graph that integrates spatial proximity and feature similarity, which is processed through a graph attention encoder to generate modality-specific embeddings. These embeddings are fused through a modality attention layer, guided by an uncertainty-aware contrastive learning objective, which preserves informative modality-specific signals while aligning shared spatial structures. SCIGMA’s uncertainty quantification framework provides location-specific uncertainty estimates, highlighting regions where multi-modal alignment is less reliable. A graph attention decoder reconstructs modality-specific features from the joint representation, ensuring biologically meaningful signals are retained. SCIGMA supports various downstream analyses, including spatial domain detection, trajectory inference (PAGA), denoised feature reconstruction, and uncertainty mapping. This figure shows a two-modality example, but SCIGMA can flexibly integrate more than two modalities. While current spatial technologies profile up to five modalities, SCIGMA’s framework supports the integration of any number of modalities, ensuring adaptability to future multi-omics platforms.

We evaluated SCIGMA on a diverse collection of datasets spanning ten different tissue types, and eight different modalities, including gene expression, chromatin accessibility, protein expression, morphological imaging, histone modifications, and metabolomics (Supplementary Table 1, Supplementary Note 3). Given the current available spatial multi-modal datasets, SCIGMA seamlessly integrates two to five modalities, offering unprecedented flexibility for spatial multi-modal analysis. To benchmark its performance, we compared SCIGMA against four state-of-the-art methods (Supplementary Note 4), demonstrating its advantages through comprehensive downstream analyses (Supplementary Note 5). In datasets that other methods could process (e.g., mouse brain spatial-ATAC-RNA and mouse spleen SPOTS), SCIGMA consistently outperformed existing spatial multi-modal methods in quantitative benchmarks, showcasing superior accuracy and robustness. For datasets with significant computational demands, where other methods failed to generate results even after two days of runtime and over 400 GB of CPU memory (Supplementary Table 2), we focused our analysis exclusively on SCIGMA. Notably, SCIGMA scales to large spatial datasets without encountering out-of-memory (OOM) issues, enabled by a graph sampling strategy that ensures computational efficiency (Method and Materials). It achieves high reproducibility across runs, addressing a key limitation of many deep learning-based integration models. Importantly, SCIGMA is the only method that flexibly supports all available modalities while also providing uncertainty quantification, scaling seamlessly to up to five modalities in a single analysis. This scalability and versatility position SCIGMA as a transformative tool in spatial multi-modal data integration.

### Mouse brain spatial-ATAC-RNA data

To benchmark SCIGMA against other competing methods, we first examined a mouse brain spatial epigenome-transcriptome dataset^16^, which measured the transcriptome and chromatin accessibility of a postnatal day 22 (P22) mouse brain via spatial ATAC-RNA-seq. We obtained the annotations of the brain anatomical regions from Allen brain atlas reference, including cortex layers 2-6, genu of the corpus callosum (CCG), Ventral lateral (VL), caudoputamen (CP), nucleus accumbens (ACB), anterior commissure (ACO), and lateral preoptic area (LPO) (Figure 2A). First, we examined each modality individually and found that the clustering of both modalities was scattered and noisy. While certain regions, such as the cortex layers and CCG, were partially distinguishable, overall spatial structures remained poorly defined (Figure 2B). Second, we used the learned joint representations from each method to cluster the spatial domains. When integrating the two modalities, SCIGMA captured all key anatomical regions. For example, SCIGMA distinctly identified various brain tissue structures, such as cortex layers 2-6 (clusters 1, 2, 4, 8), CCG (cluster 13), CP (cluster 5, 16), ACB (cluster 3), ACO (cluster 13), VL (cluster 14), and LPO (cluster 17) (Figure 2B, Supplementary Figure 2), aligning closely with annotations from the Allen Reference Atlas. In contrast, SpatialGlue generated a noisier identification of cortex regions, with scattered clusters across different layers and failed to accurately capture the LPO structure. Similarly, Seurat, MOFA+, and MultiVI failed to differentiate different cortex layers, as well as CP and ACB regions (Figure 2B), producing noisy structures. Third, we thoroughly evaluated the quality of clusters detected by each method using a series of quantitative metrics (Materials and Methods). Specifically, we calculated Moran’s I to assess the spatial autocorrelation of detected clusters, reflecting how well spatial structure is preserved, and Jaccard similarity coefficients to assess the consistency between the original modality-specific features and the model’s learned joint representation. Our rationale is that a well-integrated representation should 1) retain spatial organization inherent to the tissue, and 2) preserve modality-specific biological signals. Moran’s I quantifies the spatial coherence of the inferred clusters, while the Jaccard index evaluates the extent to which modality-specific information is retained in the integrated representation. SCIGMA consistently outperformed existing methods across these metrics - it achieved the highest Moran’s I (median = 0.862), indicating strong spatial coherence, as well as the highest Jaccard similarity (mean RNA Jaccard index = 0.219 and mean ATAC Jaccard index = 0.227), demonstrating superior preservation of modality-specific features (Figure 2C-2D, Supplementary Table 3A). Furthermore, reproducibility remains a critical challenge in deep learning, particularly in biomedical data science^28,30^. To assess model stability and consistency, we measured pairwise adjusted Rand index (ARI) and Normalized Mutual Information (NMI) across 20 random seeds on the clustering results and compared SCIGMA with other deep learning models, including MultiVI and SpatialGlue. SCIGMA achieves a significant improvement in consistency, outperforming SpatialGlue and MultiVI by 7% and 113% in ARI and 5% and 54% in NMI, respectively (Figure 2E), likely due to the stabilizing effect of its uncertainty-aware objective function, as confirmed in ablation studies (Figure S1E).

**Figure 2.**
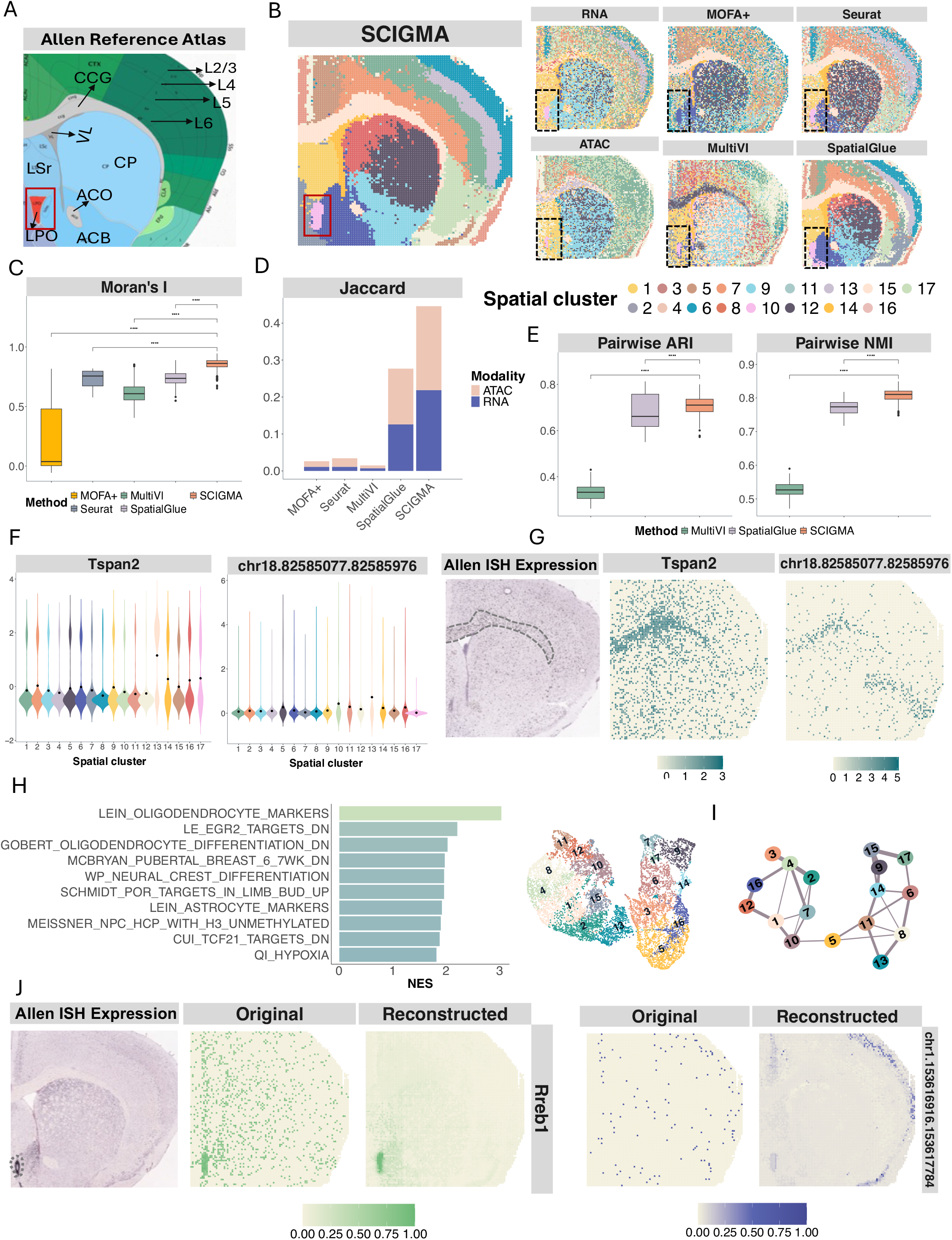
Analysis of the Spatial-epigenome-transcriptome mouse brain dataset. (**A**) Annotated mouse brain sections from the Allen Institute Mouse Brain Atlas, serving as a reference for detected spatial clusters. (**B**) Spatial clusters identified by single-modality and multi-modal integration methods, including Seurat, MOFA+, MultiVI, SpatialGlue, and SCIGMA. (**C**) Boxplots displaying Moran’s I value across methods, quantifying spatial autocorrelation. Each box spans the first to third quartiles, with the median represented by a horizontal line; whiskers extend to 1.5 times the interquartile range. P-values are calculated using a one-sided Wilcoxon rank sum test. ***: p-value < 0.001 (**D**) Jaccard similarity scores comparing preservation of modality-specific relationships in the joint representation across different methods. (**E**) Boxplots displaying pairwise ARI and NMI, which assess the consistency of clustering results across 20 random seeds for each method. (**F**) Violin plots showing the expression of key marker genes and chromatin accessibility peaks in the corpus callosum (CCG) region. (**G**) Spatial visualization of marker gene and peak expression in the CCG region, highlighting spatial patterns detected by SCIGMA. The corresponding Allen Mouse Brain ISH images are shown as a reference to support the anatomical relevance. (**H**) Bar plot of the normalized enrichment scores (NES) for the top 10 enriched gene sets in the CCG region, as identified by GSEA. (**I**) PAGA graph and UMAP embeddings derived from SCIGMA’s joint representation, showing inferred relationships between spatial domains and capturing their overall organization within the tissue. (**J**) Spatial distribution of key marker features in SCIGMA’s reconstructed data. The corresponding Allen ISH images are included for reference to validate anatomical localization.

To validate SCIGMA-detected spatial clusters, we performed downstream analyses to characterize the transcriptomic landscape of the tissue domains (Methods and Materials). First, we conducted differential expression (DE) analysis to identify genes and peaks that were specifically expressed in different spatial domains (Supplementary Figure 3). For example, *Tspan2*, a known marker of the CCG region, was identified as a DE gene and highly expressed in the CCG region (cluster 13, Figure 2F-2G). Similarly, *Mbp*^*31*^, and *Mobp*^32^, key oligodendrocyte marker genes, showed strong expression in the same region (Supplementary Figure 4). In the striatum, *Pde10a*, a well-known striatum marker gene^33^, was differentially expressed and exhibited high expression in the region (cluster 5 and 16) (Supplementary Figure 4). Beyond known makers, we also identified novel marker genes. For example, *Top2a*, a validated marker gene of neuronal intermediate progenitor cells^34^, was highly expressed in the lateral ventricle (VL region, cluster 14, Supplementary Figure 4). While *Snap25*, a marker of excitatory neurons in the cortex^35^, was enriched in cortex layer L5 (cluster 1, Supplementary Figure 4). Notably, the spatial expression patterns of these marker genes were consistent with reference ISH data from the Allen Mouse Brain Atlas, supporting the anatomical accuracy of SCIGMA-derived clusters (Figure 2G, Supplementary Figure 4). In the ATAC-seq data, we identified a peak (chr18.82585077.82585976) that was highly expressed in the CCG region (cluster 13) (Figure 2F-2G, Supplementary Figure 5). Second, we assessed the integration of RNA and ATAC signals by generating a peak-to-gene links heatmap following the original study^16^ (Supplementary Figure 6). Modality-specific clustering showed that cortex layers (clusters 2, 4, 8) were more clearly defined in RNA-based clustering (Figure 2B), whereas the corpus callosum (cluster 13) was more prominent in ATAC-based clustering, and the striatum (clusters 5, 16) appeared in both. The heatmap revealed that SCIGMA successfully captured these spatial domains in a unified representation, even when they were initially modality specific. These results again illustrate SCIGMA’s effectiveness in aligning and integrating spatial transcriptomic and epigenomic signals, enabling the detection of biologically relevant spatial domains that would be missed by unimodal analyses alone. Third, we performed gene set enrichment analysis (GSEA) on the detected genes (Figure 2H, full results in Supplementary Figure 7). The results showed strong enrichment in pathways related to oligodendrocyte and neural crest development, particularly in the CCG region (Figure 2H). To explore the relationships among detected spatial clusters, we conducted PAGA trajectory analysis on SCIGMA-generated embeddings (Materials and Methods, Figure 2I). The results captured domain transitions, such as those between the LPO (cluster 17) and ACB (cluster 6), cortex layers L4 (cluster 4) and L5 (cluster 2), and LSr (cluster 9) and VL (cluster 14). The UMAP further highlighted a clear progression from cortex layers (clusters 1, 2, 4, 8) to the striatum (clusters 3, 5, 16) and relationships between LPO (cluster 17) and ACB (cluster 6). We also examined the modality-specific weights learned by SCIGMA to assess the relative contribution of different modalities to each cluster (Supplementary Figure 8). SCIGMA assigned higher attention to RNA in cortex layers L2/3/4 (clusters 4 & 8) and higher attention to ATAC in the CCG (cluster 13) and LPO (cluster 17) regions, reflecting biologically meaningful modality contributions. This suggests that SCIGMA effectively captures region-specific molecular signals, leveraging RNA for transcriptionally active cortical layers^36^ and ATAC for chromatin accessibility patterns in white matter regions^37^. These results highlight the strength of SCIGMA’s attention mechanism in adaptively integrating spatial multimodal information.

Beyond identifying and characterizing spatial domains, SCIGMA demonstrated strong robustness in reconstructing spatial multi-modal data (Materials and Methods). The reconstructed profiles effectively denoised transcriptomic signals, which are often sparse due to technological limitations such as dropouts and insufficient detection of lowly expressed genes. For example, the reconstructed expression of *Rreb1*^*38*^, a marker gene for neuronal proteostasis, exhibited a stronger, more localized pattern compared to the scattered raw data (Figure 2J), showing high concordance with the expression pattern observed in the Allen Mouse ISH reference image. Similarly, the reconstructed ATAC profile for the peak chr1.153616916.153617784 displayed a more distinct enrichment in cortex layer 1 than the raw data. Additional examples are presented in Supplementary Figure 9. These results highlight SCIGMA’s ability to mitigate technical noise while preserving biologically relevant signals. Finally, to interpret SCIGMA’s multi-modal integration, we examined the model’s uncertainty estimates (Methods and Materials). Spatial mapping of uncertainty estimates revealed particularly high uncertainty in the VL, LPO, and certain cortex regions (Supplementary Figure 10A). We found that regions with higher uncertainty also exhibited greater discrepancies between RNA and ATAC signals (Supplementary Figure 10B), indicating that SCIGMA assigns higher uncertainty in regions where the modalities diverge. Notably, regions with higher uncertainty exhibited greater modality-specific reconstruction errors (RMSE) and lower Jaccard similarity between modality-specific and joint embeddings (Supplementary Figures 10C–10D). These findings suggest that SCIGMA’s uncertainty estimates reflect regions where the alignment of multi-modal signals is less certain, potentially due to biological heterogeneity or data quality differences.

### Mouse spleen SPOTS data

Next, we demonstrated SCIGMA’s ability to integrate RNA and protein expression using mouse spleen spatial profiling data from the SPOTS platform^17^, which captures whole transcriptomes and extracellular proteins through polyadenylated antibody-derived tag (ADT)-conjugated antibodies at a spot level. The detection panel was designed to identify surface markers of B cells, T cells, and macrophages—cell types abundantly present in the spleen. The spleen, an essential organ in the lymphatic system, is responsible for filtering blood and supporting immune functions^39,40^. It consists of two major compartments: the red and white pulp. The red pulp, which contains red pulp macrophages (RpMΦ) responsible for filtering and recycling red blood cells, plays a key role in maintaining blood homeostasis. The white pulp contains periarteriolar lymphoid sheaths with T cells and B cell follicles with germinal centers that facilitate immune responses. Adjacent to the white pulp is the marginal zone (MZ), a specialized interface situated between the white pulp and the red pulp. The MZ is densely populated with two distinct macrophage populations: marginal zone macrophages (MZMΦ), which are primarily located at the outer border of the marginal zone and specialized in capturing blood-borne antigens, and metallophilic macrophages (MMMΦ), which are found lining the inner border of the marginal zone, forming a ring-like structure around the white pulp. These macrophages play critical roles in antigen uptake and initiating immune responses^41,42^ (Figure 3A). We first performed clustering of each modality to examine their correspondence between RNA and protein data. The results clearly showed that the RNA and protein modalities capture different biological structures. For example, both modalities roughly identified the white pulp region, where T cells (cluster 5) and B cells (cluster 4) are located. However, the RNA modality mixed the red pulp region (cluster 2, where RpMΦ are located) with the MZ region (cluster 3, where MZMΦ are located), while the protein modality failed to identify the inner border of the MZ (cluster 1, where MMMΦ are located) (Figure 3B). Second, we used the learned joint representations from each method to cluster the spatial domains. SCIGMA effectively captured all key regions (Figure 3B, Supplementary Figures 11). For example, SCIGMA identified clearer structures, including the red pulp (cluster 2), MZ (cluster 3), inner border of the MZ (cluster 1), subclusters within the white pulp (cluster 4, 5). In contrast, multiVI failed to distinguish the red pulp and the MZ, and could not identify the inner border of the MZ as well as subclusters within the white pulp where T and B cells reside. Seurat and MOFA+ performed similarly, both failing to detect the MZ accurately (Figure 3B). SpatialGlue was able to detect the MZ structure, but misclassified parts of the MZ as belonging to the red pulp and failed to capture the inner MZ border, where MMMΦ are located (cluster 1) (Figure 3B). Quantitative metrics further demonstrated SCIGMA’s superior performance, achieving the highest Moran’s I score (median = 0.685), and Jaccard similarity coefficient (mean Jaccard index for RNA = 0.123 and mean Jaccard index for protein = 0.144, Figure 3C - 3D, Supplementary Table 3B), These results confirm SCIGMA’s effectiveness in identifying spatially autocorrelated clusters while preserving modality-specific information in the joint representation. Furthermore, we evaluated result consistency across 20 seed initializations, SCIGMA once again demonstrated significant improvements over other deep learning methods, outperforming SpatialGlue and MultiVI by 23% and 4319% in ARI and 16% and 2340% in NMI respectively (Figure 3E).

**Figure 3.**
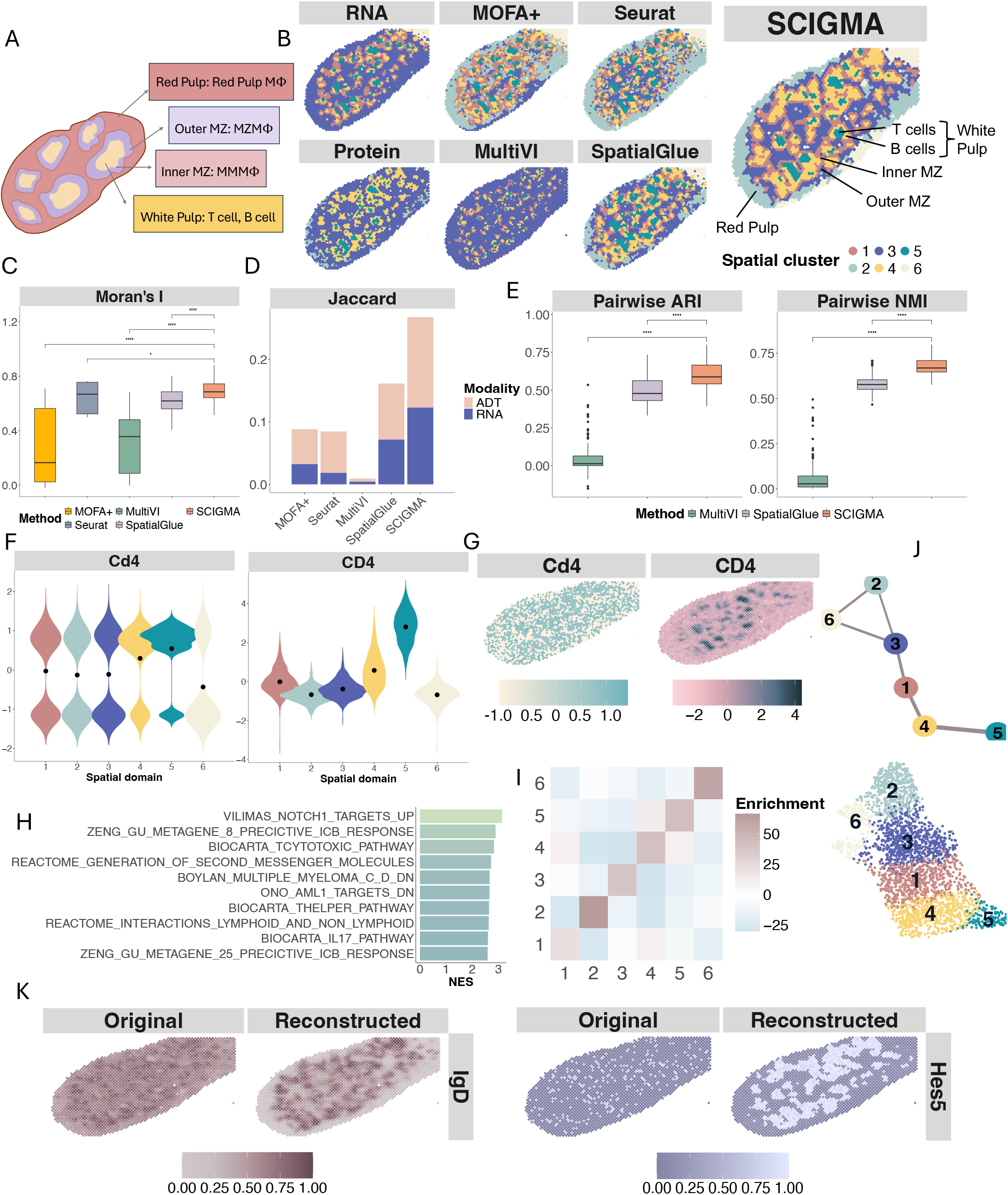
Analysis of the SPOTS transcriptome-proteome mouse spleen dataset. (**A**) Schematic illustration of mouse spleen structure, highlighting key anatomical compartments including the red pulp, marginal zone (MZ), and white pulp, where T cells, B cells, and macrophages reside. The marginal zone contains two specialized macrophage populations: marginal zone macrophages (MZM Φ) at the outer border and metallophilic macrophages (MMMΦ) along the inner border. (**B**) Spatial clusters identified by single-modality and multi-modal integration methods, including Seurat, MOFA+, MultiVI, SpatialGlue, and SCIGMA. (**C**) Boxplots displaying Moran’s I values across methods, quantifying spatial autocorrelation. Each box spans the first to third quartiles, with the median represented by a horizontal line; whiskers extend to 1.5 times the interquartile range. (**D**) Jaccard similarity scores comparing preservation of modality-specific relationships in the joint representation across different methods. (**E**) Boxplots displaying pairwise ARI and NMI, assessing the consistency of clustering results across 20 random seeds for each method. (**F**) Violin plots showing the expression of key marker genes and proteins in the T cell region. (**G**) Spatial visualization of marker gene and protein expression in the T cell region, highlighting spatial patterns detected by SCIGMA. (**H**) Bar plot of the normalized enrichment scores (NES) for the top 10 enriched gene sets in the T cell region, identified via GSEA. (**I**) Neighborhood enrichment analysis quantifying spatial associations between identified clusters. (**J**) PAGA graph and UMAP embeddings derived from SCIGMA’s joint representation, showing inferred relationships between spatial domains and capturing their spatial organization within the tissue. (**K**) Spatial distribution of key marker features in SCIGMA’s reconstructed data.

We carefully examined the molecular and cellular signatures of the spatial clusters identified by SCIGMA. First, domain-specific DE analysis identified known markers associated with different clusters (Supplementary Figure 12). For example, cluster 5, corresponding to the regions where T cells are located, exhibited high enrichment of the gene *Cd4*, a well-known T cell marker^43^ (Figure 3F-G). In addition, DE analysis revealed that this region was enriched for known immune markers *Tcf7* and *Serpina10*^*44,45*^ (Supplementary Figure 13). SCIGMA also identified novel marker genes. In cluster 2, which corresponds to the red pulp, *Nfe2* and *Ctse* were identified as differentially expressed genes (Supplementary Figure 13). Notably, *Nfe2* plays a key role in regulating oxidative stress^46^, while *Ctse* is involved in protein degradation^47^. These findings are consistent with the known functions of the red pulp, highlighting the potential of these genes as markers for this domain. Additionally, we identified important protein markers, including *CD3, CD4*, and *CD8* for T cells^48,49^, *CD19* and *IgD* for B cells^50,51^, and *EpCAM* for the epithelial region^52^ (Figure 3F – 3G, Supplementary Figures 13 – 14). Second, GSEA analysis (full results in Supplementary Figure 15) confirmed that domain-specific DE genes for T cells were highly enriched in pathways related to Notch1 signaling and cytotoxic pathways^53,54^, both of which are key immune-related pathways (Figure 3H). Since T cells and B cells reside in the white pulp, while MMMΦ reside along the inner border of the MZ, we conducted a neighborhood enrichment analysis to quantify their spatial relationships (Materials and Methods). The analysis revealed low enrichment between red pulp, MZ, and B cell clusters (cluster 2, 3, 4) and strong enrichment between the white pulp regions, B (cluster 4) and T cell clusters (cluster 5). This confirms SCIGMA’s ability to capture their close yet distinct spatial organizations (Figure 3I). We further examined the relationships between SCIGMA’s detected spatial clusters through PAGA analysis (Figure 3J). The results showed strong associations between white pulp clusters (clusters 4 and 5), while the transition from the white pulp to the red pulp was captured by a trajectory spanning from the inner border of the MZ (cluster 1) through the MZ (cluster 3) to the red pulp (cluster 2). This trajectory aligns well with the known biological organization of the spleen, highlighting SCIGMA’s ability to detect spatial domains that are both accurate and biologically meaningful. The modality-specific weights learned by SCIGMA indicated that protein contributions were weighted higher than RNA across clusters, which aligns with protein markers showing clearer spatial patterns than RNA markers (Supplementary Figure 16).

Importantly, SCIMA effectively denoised and reconstructed RNA and protein expression patterns (Figure 3K, Supplementary Figure 13A-13B). The reconstructed data revealed more well-defined spatial patterns for protein and RNA markers, providing deeper insights into cellular function and tissue architecture. For example, SCIGMA successfully reconstructed and denoised the spatial patterns of B cell protein marker *IgD*^*50*^ and T cell RNA marker *Hes5*^*55*^ (Figure 3K). Additional examples of these improvements are provided in Supplementary Figure 17, further demonstrating SCIGMA’s ability to reconstruct key feature patterns despite technical variations. Finally, to assess the reliability of SCIGMA’s multi-modal integration and identify regions where modality alignment may be less certain, we analyzed the model’s uncertainty estimates (Methods and Materials). Spatial mapping revealed that uncertainty was highest in the red pulp and epithelial regions (Supplementary Figure 18A). Higher uncertainty values were observed in areas with greater discrepancies between RNA and protein signals (Supplementary Figure 18B), indicating that SCIGMA assigns increased uncertainty where the modalities show less agreement. In these high-uncertainty areas, reconstruction RMSE of both modalities was also higher (Supplementary Figure 18C), suggesting that SCIGMA had greater difficulty reconstructing modality-specific features. However, Jaccard similarity between each modality-specific embedding and the joint embedding was higher in uncertain areas, suggesting that in uncertain regions, SCIGMA was able to rely more on shared structural features across modalities to maintain reasonable alignment of the joint representation (Supplementary Figure 18D). This indicates that uncertainty in these regions reflects weak or ambiguous modality-specific signals rather than complete misalignment, with the model emphasizing common spatial structures. Thus, the weaker feature reconstruction but preservation of spatial topology in the embedding space suggests that uncertainty values capture a mixture of weak biological variation or technical noise without significantly disrupting the joint representation.

### 10x Xenium Prime data

We further demonstrated SCIGMA’s ability to integrate imaging and transcriptomic modalities using two datasets from the 10x Xenium Prime 5K platform. Compared to the 10x Xenium v1, Xenium Prime 5K detects a higher median number of transcripts per cell and achieve lower false discovery rates in both human and mouse tissues^56^. This platform simultaneously captures transcriptomic and morphological features, measuring up to 5,000 genes per assay. For this analysis, we focused on two datasets derived from fresh-frozen human ovarian adenocarcinoma and the FFPE human cervical cancer^19^. Since the compared methods do not support imaging modalities, this analysis focuses exclusively on SCIGMA.

We first analyzed the human ovarian adenocarcinoma dataset, which represents a malignant tumor originating from ovarian epithelial cells. This cancer type is characterized by high intratumoral heterogeneity, with well-defined glandular structures and dense stromal regions influencing tumor progression, immune infiltration, and therapy resistance^57^ (Figure 4A). Integrating spatial transcriptomics and imaging is crucial for understanding tumor microenvironment (TME). First, we examined clustering results from each modality separately. RNA expression captured more heterogeneous biological structures than imaging features, identifying distinct substructures that were not apparent from imaging-based clustering alone (Methods and Materials, Figure 4B). However, both modalities failed to resolve finer structures within the tissue, suggesting limitations in their individual capacities to fully characterize the spatial complexity of ovarian cancer. In contrast, SCIGMA successfully identified key tumor- and immune-related clusters by integrating both modalities. As this dataset lacked predefined annotations, we validated the SCIGMA-identified clusters through comprehensive downstream analyses (Figure 4B, Supplementary Figure 19). First, domain-specific DE analysis identified both cancerous and non-cancerous marker genes associated with SCIGMA-detected clusters (Supplementary Figure 20). Specifically, SCIGMA distinguished normal ovarian tissue from cancerous regions. For example, *SCGB1D2*, a secretory protein gene associated with hormone-responsive epithelial tissues^58,59^, and *HOXB5*, a transcription factor linked to differentiation and tumor progression^60^, were highly expressed in cluster 2 (Figure 4C, Supplementary Figure 21). This suggests that cluster 2 may represent more differentiated tumor regions with glandular or epithelial characteristics. In contrast, *H19*, a non-coding RNA known to promote angiogenesis and tumor invasiveness^61^, and *RSPO3*, a gene that activates the Wnt/*β* -catenin signaling pathway associated with cancer stemness and vascular remodeling^62^, were highly expressed in cluster 17 (Figure 4C, Supplementary Figure 21). These findings suggest that cluster 17 represents more aggressive tumor substructures, such as invasive fronts or hypoxic niches. Additionally, ovarian cancer markers *CCL8 and LAMA3* were highly expressed in clusters 10 and 11, further supporting their classification as tumor regions^63,64^. These findings highlight SCIGMA’s ability to identify heterogeneous tumor subregions, each characterized by distinct biological processes such as differentiation, and invasiveness. SCIGMA also identified several non-cancerous subregions with distinct biological functions. For example, cluster 9 corresponded to an endothelial region, characterized by the high expression of endothelial cell marker genes *KDR* and *CDH5*^65,66^ (Supplementary Figure 21). Cluster 7 was associated with immune regions, marked by the high expression of immune cell related marker genes *CD8A* and *TNFRSF18*^67,68^ (Supplementary Figure 21). Additionally, Clusters 8, 15, and 20 corresponded to granulosa cells, exhibiting strong expression of the granulosa-specific marker genes *FOXL2* and *ESR2*^69,70^(Supplementary Figure 21). GSEA analysis further validated SCIGMA’s spatial clusters by showing that domain-specific DE genes were highly enriched in ovarian cancer-related pathways (Supplementary Figure 22). For example, cluster 2 was significantly enriched in ovarian cancer, *Myc* oncogene target genes^71^, and hypoxia pathways (Figure 4D), reinforcing its classification as a cancerous region. We further examined cluster relationships using PAGA and UMAP (Figure 4E). PAGA trajectories revealed strong inter-cluster relationships among cancer marker-enriched clusters (clusters 2, 10, 11, and 17). Notably, granulosa clusters (clusters 8, 15, and 20) exhibited strong connectivity with other domains, aligning with their central role in hormone production. SCIGMA’s learned modality-specific weights indicated that both modalities contributed equally to the tumor subtypes such as clusters 2 and 17, but RNA modality played a greater role in identifying granulosa cell clusters (cluster 15, Supplementary Figure 23).

**Figure 4.**
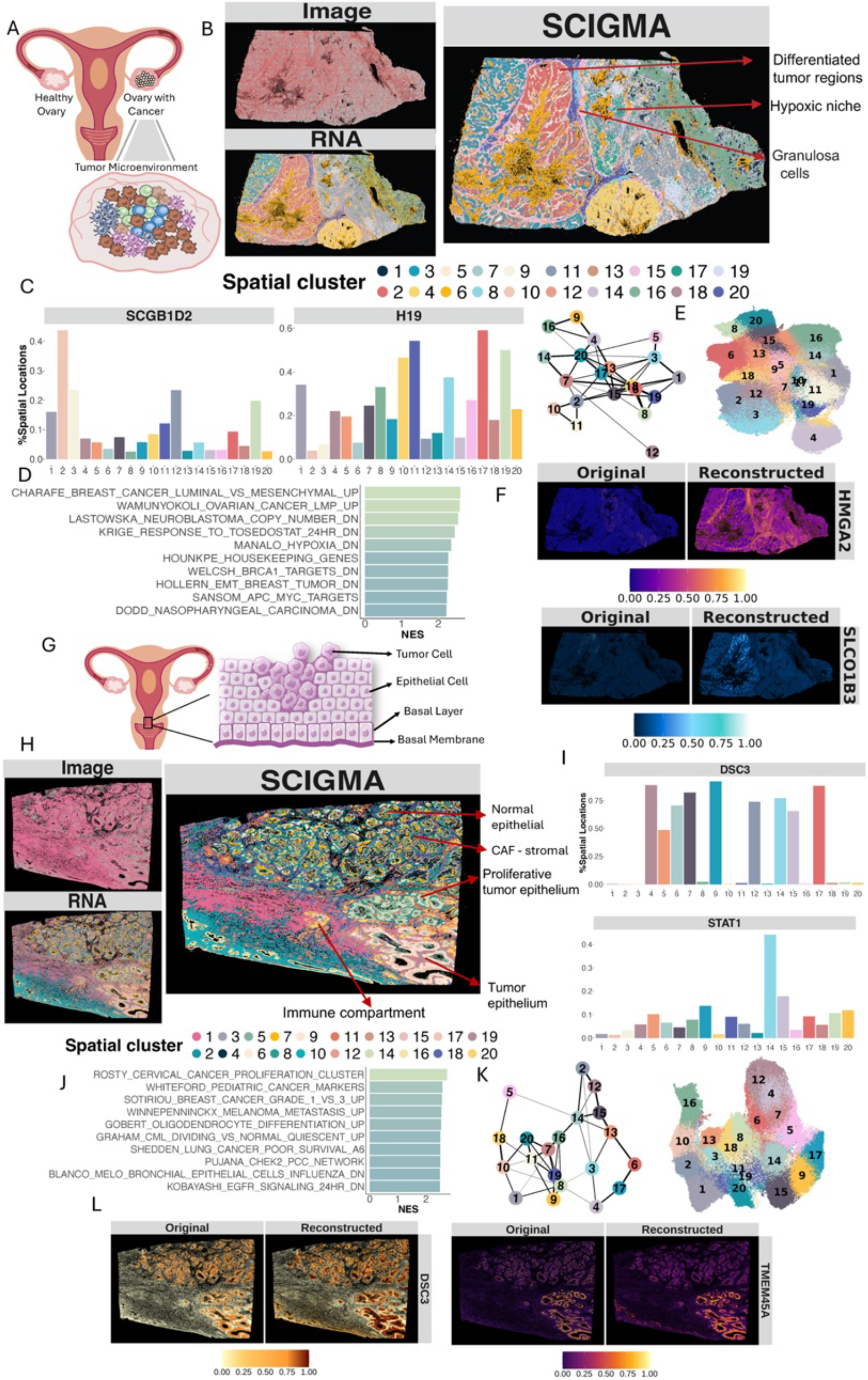
Uncovering tumor heterogeneity with 10x Xenium Prime cancer datasets. (**A**) A diagram displays the tumor microenvironment of the human ovarian adenocarcinoma tissue. (**B**) Spatial clusters identified by single-modality, and SCIGMA. (**C**) Barplots showing the percentage of spatial locations with expression levels above the median for key cancer marker genes across SCIGMA-identified clusters in the ovarian adenocarcinoma dataset. (**D**) Bar plot of the normalized enrichment scores (NES) for the top 10 enriched gene sets in ovarian cancer cluster 2, identified via GSEA, highlighting pathways associated with glandular and epithelial tumor characteristics. (**E**) PAGA graph and UMAP embeddings derived from SCIGMA’s joint representation, showing inferred relationships between spatial domains and capturing their spatial organization within the ovarian tumor tissue. (**F**) Spatial distribution of key cancer markers in SCIGMA’s reconstructed gene expression data. (**G**) A diagram displays the basic anatomical layers of the human cervical cancer tissue (**H**) Spatial clusters identified by single-modality, and SCIGMA. (**I**) Barplots showing the percentage of spatial locations with expression levels above the median for key cancer marker genes across SCIGMA-identified clusters in the cervical cancer dataset. (**J**) Bar plot of the normalized enrichment scores (NES) for the top 10 enriched gene sets in cervical cancer cluster 14, identified via GSEA, highlighting pathways associated with cervical cancer progression. (**K**) PAGA graph and UMAP embeddings derived from SCIGMA’s joint representation, showing inferred relationships between spatial domains and capturing their spatial organization within the cervical tumor tissue. (**L**) Spatial distribution of key cancer markers in SCIGMA’s reconstructed gene expression data.

In the ovarian adenocarcinoma dataset, SCIGMA successfully denoised and enhanced the expression signals of key cancer markers, such as *SLCO1B3* and *HMGA2*, revealing clearer spatial localization patterns that align with known tumor biology (Figure 4F). For example, in the reconstructed gene expression data, *SLCO1B3* was highly expressed in cluster 2, which corresponds to more differentiated tumor regions with glandular or epithelial characteristics. This aligns with *SLCO1B3*’s role as a solute carrier involved in metabolic processes and its frequent upregulation in hypoxia-driven tumor cells^72^. Its association with differentiated regions suggests a role in maintaining metabolic activity and function in glandular or epithelial-like tumor cells, highlighting SCIGMA’s biological relevance in identifying spatially distinct tumor substructures. Similarly, *HMGA2* reconstruction showed highest expression in clusters 17 and 11. Cluster 17 corresponds to aggressive tumor substructures, while clusters 11 are associated with ovarian cancer markers. Because of *HMGA2’s* relation to granulosa proliferation, the reconstructed features also showed elevated feature expression in granulosa cell regions (clusters 8, 15, 20), showing that SCIGMA was able to pick up relationships between normal and cancerous regions. This pattern aligns with *HMGA2*’s known role as a chromatin remodeling protein involved in promoting tumor proliferation and invasion, as well as its potential function in supporting cellular activity and proliferation within granulosa cell populations in ovarian tissue^73,74^. More examples of these improvements are presented in Supplementary Figure 24, illustrating the benefits of SCIGMA on mitigating the effects of the technical variations in feature reconstruction. Furthermore, we examined SCIGMA’s uncertainty estimates. Spatial mapping of uncertainty values revealed that model uncertainty was higher in the granulosa and endothelial regions, both of which are structurally normal tissue areas in the ovary (Supplementary Figure 25A). Higher uncertainty values were observed in regions with greater discrepancies between RNA and imaging signals (Supplementary Figure 25B), indicating that uncertainty captures areas where the modalities provide less consistent information density. Additionally, uncertain regions exhibit lower reconstruction RMSE and higher Jaccard similarity, indicating that SCIGMA effectively preserves biological structure despite uncertainty (Supplementary Figure 25C - 25D). This suggests that the observed uncertainty arises from modality-specific technical differences in feature alignment, rather than a loss of biological information, and does not compromise the quality of the learned embeddings. Consistent with this, the highest uncertainty values were observed in structurally normal tissue regions, rather than in the more heterogeneous tumor areas, where biological variation would be expected to pose greater alignment challenges. Based on these observations, we hypothesize that SCIGMA’s uncertainty estimates primarily capture technical variability arising from the integration of RNA and imaging features within the joint representation.

Following the analysis of the ovarian adenocarcinoma dataset, we applied SCIGMA to the human cervical cancer dataset. Cervical cancer predominantly consists of squamous cell carcinoma or adenocarcinoma, forming well-demarcated spatial regions including epithelial, stromal, and immune components^75^ (Figure 4G). The RNA modality captured structures potentially corresponding to cancerous regions that were not apparent in the imaging modality (Methods and Materials). After integrating the two modalities, SCIGMA successfully delineated heterogeneous spatial domains (Figure 4H, Supplementary Figure 26), distinguishing both normal and cancerous regions. For example, we found that cervical cancer markers *DSC3* and were highly expressed across multiple spatially organized clusters (clusters 4, 6, 7, 9, 12, 14, 15, and 17), while marker genes such as *TMEM45A* and *STAT*^76,77^ were uniquely highly expressed in cluster 14 (Figure 4I, Supplementary Figures 27 - 28). Additionally, DE analysis identified marker genes for cancer-associated fibroblasts (CAFs) such as *SERPINB2, COL5A2*, and *POSTN*^78-80^, which were strongly expressed in cluster 8, distinguishing this region from other cancerous regions (Supplementary Figure 28). Cluster 13 exhibited high expression of epithelial tumor marker genes *TP63* and *ADAMTS1*^81,82^, further supporting SCIGMA’s ability to capture tumor heterogeneity. Beyond identifying tumor regions, SCIGMA also detected distinct normal tissue compartments. Cluster 20 showed high expression of immune marker genes *CD8A* and *CD3D/E*^48,68^, indicating the presence of immune cells (Supplementary Figure 28). Cluster 18 corresponded to endothelial cells, characterized by the high expression of marker genes such as *PECAM1* and *COL4A1*^83,84^ (Supplementary Figure 28). Additionally, SCIGMA captured epithelial progression, with cluster 5 displaying strong expression of epithelial marker genes such as *CECAM5, KRT78, TMPRSS2*^85-87^. These results highlight SCIGMA’s ability to resolve fine-grained spatial domains within both cancerous and normal regions of cervical tissue. GSEA analysis on cluster 14 further revealed significant enrichment in cancer-related pathways, including those associated with cervical cancer progression (e.g., ROSTY_CERVICAL_CANCER_PROLIFERATION_CLUSTER pathway) (Figure 4J, Supplementary Figure 29). PAGA trajectory analysis showed strong connections between cancer clusters (cluster 4, 6, and 17, as well as clusters 14 and 15) and relationships between endothelial and epithelial domains (cluster 18, and 5), suggesting the interconnected nature of these tissue regions (Figure 4K). The modality-specific weights learned by SCIGMA suggests that the model attended to the RNA modality more, suggesting higher feature richness in RNA compared to imaging (Supplementary Figure 30). This trend was consistent across most tumor subclusters, with the exception of cluster 15, where SCIGMA assigned greater weight to imaging features, while the other tumor subregions were predominantly shaped by RNA features. These results underscore the importance of modality-specific contributions in integrating heterogeneous spatial data. SCIGMA also demonstrated its denoising capabilities in the cervical cancer dataset, enhancing the spatial expression of key cancer-related genes. For instance, *DSC3*, and *TMEM45A* showed clearer and more localized spatial patterns in the reconstruction data (Figure 4L, more examples are in Supplementary Figure 31). Finally, we examined SCIGMA’s uncertainty estimates to assess the reliability of multi-modal integration and identify regions where modality alignment may be less certain. Spatial mapping of uncertainty values revealed that uncertainty was particularly high in heterogeneous tumor regions rather than in structurally normal tissue areas (Supplementary Figure 32A), suggesting that tumor heterogeneity poses greater challenges for multi-modal alignment. Regions with higher uncertainty also showed greater discrepancies between RNA and imaging features (Supplementary Figure 32B), indicating that SCIGMA assigns higher uncertainty where the two modalities provide differing information content. These uncertain regions were further associated with higher reconstruction error (RMSE), suggesting that differences between RNA and imaging features lead to increased difficulty in accurately reconstructing the original data in the joint representation (Supplementary Figure 32C). Despite these challenges, the uncertain regions still exhibited comparable or even higher Jaccard similarity between modality-specific and joint embeddings, indicating that SCIGMA preserves spatial consistency between RNA and imaging features even in regions with greater uncertainty (Supplementary Figure 32D). Together, these results suggest that while shared spatial structures support spatial alignment across modalities, biological heterogeneity stemming from the weakens the alignment of finer molecular features in the joint representation. Given that uncertainty is concentrated in tumor regions, we hypothesize that SCIGMA’s uncertainty estimates primarily reflect the inherent complexity of tumor microenvironment, where increased heterogeneity makes multi-modal alignment more challenging.

### High-resolution 10x VisiumHD data

To demonstrate SCIGMA’s scalability, we applied it to high-resolution datasets generated from the 10x VisiumHD platform^19^. This platform provides transcriptomic profiling at up to 2 *μm* resolution alongside high-quality morphology images, enabling exploration of tissue architecture at cellular and subcellular levels. With over a million spatial locations, these large-scale datasets pose significant computational challenges that most existing spatial transcriptomics methods are not designed to efficiently handle. SCIGMA addressed these challenges using an efficient graph sampling algorithm, enabling it to capture tissue heterogeneity in its learned representations. For these large-scale datasets, which measure gene expression across 500K ∼ 1 million locations with paired imaging data, we only focused on SCIGMA, as all other models lacked support for imaging data and/or were unable to handle datasets of this scale (given 48 hours of 400Gb of CPU RAM and 24Gb of GPU RAM). We evaluated SCIGMA on two VisiumHD datasets: human colorectal cancer (CRC) and the mouse small intestine datasets.

CRC is one of the most common malignancies worldwide, characterized by heterogenous tissue architecture, including cancerous epithelial regions, stromal compartments, and infiltrating immune cells^88^ (Figure 5A). Cancerous regions arise from epithelial cells forming abnormal glandular structures with varying degrees of differentiation, while surrounding stromal compartments consist of fibroblasts, blood vessels, and extracellular matrix, supporting tumor growth. Infiltrating immune cells contribute to either anti-tumor immunity or immune evasion. In the CRC dataset, RNA expression provided a clearer representation of the tissue’s layered substructures compared to the more fragmented patterns captured by imaging features (Methods and Materials, Figure 5B). By integrating both modalities, SCIGMA delineated spatial domains more clearly, enhancing the identification of tissue subregions (Figure 5B, Supplementary Figure 33). Since this dataset lacks pre-defined annotations, we validated SCIGMA’s spatial domains through downstream analyses. Domain-specific DE analysis confirmed that SCIGMA effectively distinguished cancerous from normal regions (Supplementary Figure 34). For example, cluster 2 exhibited high expression of hypoxia-related marker genes *NDRG1* and *ANKRD37*^89,90^, indicating hypoxia-driven tumor growth, likely corresponding to poorly vascularized and aggressive tumor regions (Figure 5C, Supplementary Figures 35). In contrast, cluster 5 showed marker genes linked to cancer stem cells (*ALDH1A1*)^91^ and neuroendocrine differentiation (*MAOA*)^92^, suggesting it represents a tumor region with stem-like properties and neuroendocrine activity, which are often associated with therapy resistance and tumor recurrence (Figure 5C, Supplementary Figure 35). Cluster 6, exhibited high expression of immune evasion marker genes such as *FOSB, ATF3*, and *NR4A1*^93-95^, representing an immune-suppressed tumor microenvironment, likely driven by mechanisms that downregulate anti-tumor immune responses (Supplementary Figure 35). These distinct tumor-related regions highlight SCIGMA’s ability to differentiate tumor subtypes and microenvironmental states that influence tumor progression and immune escape. Furthermore, SCIGMA identified non-cancerous regions, such as cluster 9, with high expression of immune cell marker genes *JCHAIN, IGHA1*, and *IGKC*^*96-98*^, which are indicative of active humoral immune responses and B-cell activity (Supplementary Figure 35). This cluster may correspond to lymphoid aggregates or immune-rich areas adjacent to the tumor, reflecting the interplay between tumor and immune compartments within the colorectal cancer microenvironment. GSEA analysis further revealed that cluster 2 was enriched in pathways related to hypoxia and oncogenic signatures (Figure 5D), consistent with the DE analysis results that identified hypoxia-driven tumor markers. Full GSEA results are provided in Supplementary Figure 36. PAGA analysis further revealed distinct tumor progression patterns, with hypoxia- and stem cell-enriched tumor regions (clusters 2 and 5) closely linked, while normal-to-cancerous transitions were observed between clusters 9 and 6, reflecting immune-tumor interactions (Figure 5E). Modality-specific weights learned by SCIGMA indicated a stronger contribution from the RNA modality across clusters, suggesting that RNA features provided more informative signals for spatial domain delineation (Supplementary Figure 37).

**Figure 5.**
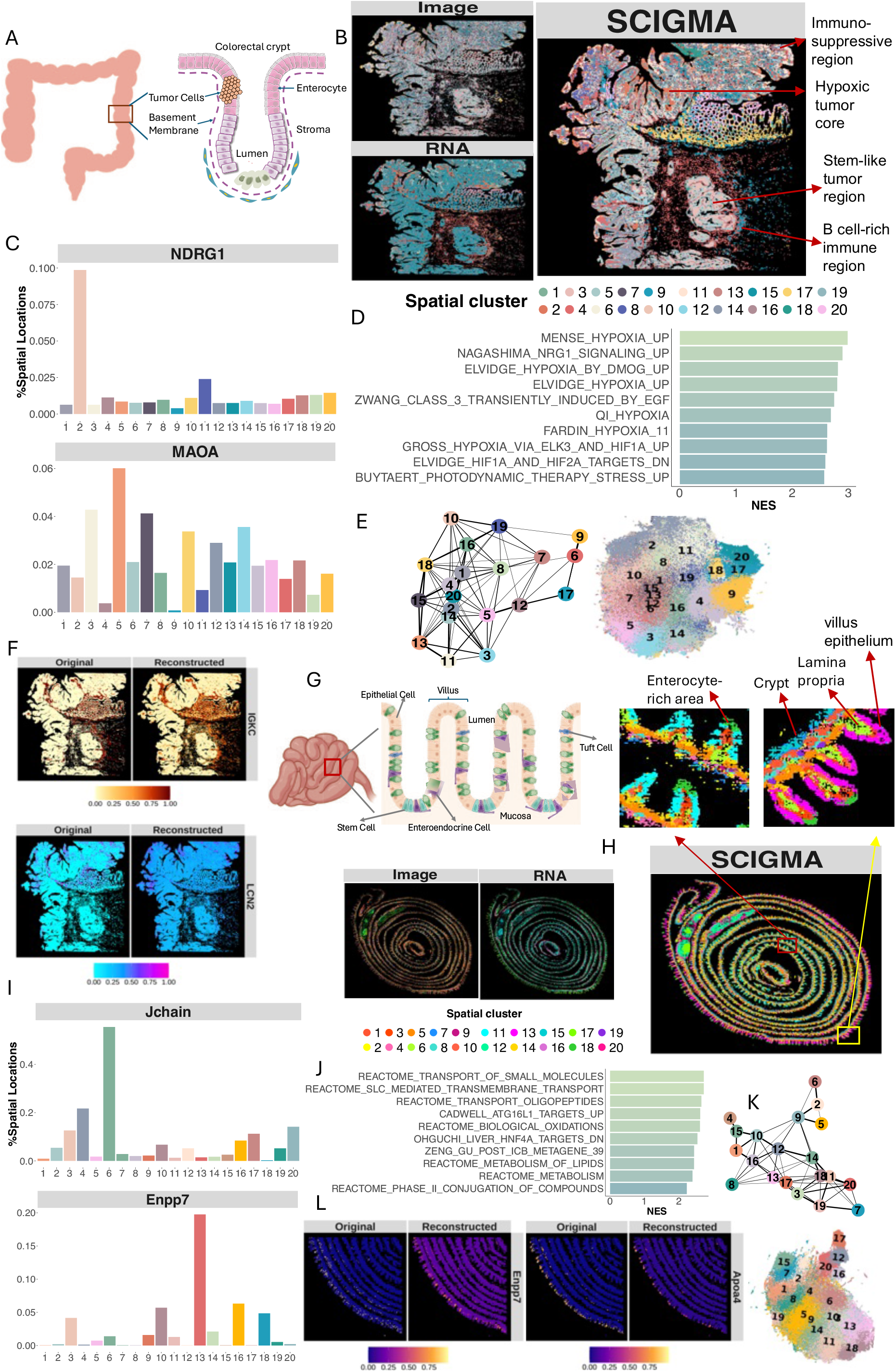
SCIGMA scales to large 10x VisiumHD datasets with over 1 million spatial locations. (**A**) The structure of the colorectal crypt in the human colorectal cancer tissue (**B**) Spatial clusters identified by single-modality, and SCIGMA. (**C**) Barplots showing the percentage of spatial locations with expression levels above the median for key cancer marker genes across SCIGMA-identified clusters in the colorectal cancer dataset. (**D**) Bar plot of the normalized enrichment scores (NES) for the top 10 enriched gene sets in colorectal cancer cluster 2, identified via GSEA, highlighting pathways associated with hypoxia and tumor progression. (**E**) PAGA graph and UMAP embeddings derived from SCIGMA’s joint representation, showing inferred relationships between spatial domains and capturing their spatial organization within the colorectal tumor tissue. (**F**) Spatial distribution of key immune and cancer markers in SCIGMA’s reconstructed gene expression data. (**G**) The structure of the mouse small intestine tissue. (**H**) Spatial clusters identified by single-modality, and SCIGMA. (**I**) Bar plots showing the percentage of spatial locations with expression levels above the median for key metabolism-related genes across SCIGMA-identified clusters in the small intestine dataset. (**J**) Bar plot of the normalized enrichment scores (NES) for the top 10 enriched gene sets in small intestine cluster 13, identified via GSEA, highlighting pathways associated with epithelial metabolism and nutrient transport. (**K**) PAGA graph and UMAP embeddings derived from SCIGMA’s joint representation, showing inferred relationships between spatial domains and capturing their spatial organization within the small intestine tissue. (**L**) Spatial distribution of key metabolism-related markers in SCIGMA’s reconstructed gene expression data.

SCIGMA effectively denoised and enhanced the sparse RNA modality, uncovering transcriptome localization patterns that were previously obscured by technical variations and data sparsity. Consistent with its performance in prior datasets, SCIGMA’s reconstruction produced more defined spatial patterns for key cancer genes. For example, SCIGMA successfully denoised immune markers such as *IGKC* and colorectal cancer markers such as *LCN2*^99^ (Figure 5F). Stronger expression of *LCN2* across tumor regions (clusters 2,5,6) further demonstrates SCIGMA’s ability to discern biological patterns despite technical noises. Additional examples are detailed in Supplementary Figure 38. Furthermore, we examined the uncertainty estimates to assess how biological and/or technical variations affect the model’s confidence in integrating the two modalities (Methods and Materials). Spatial mapping of uncertainty values revealed that tumor and immune regions exhibited the highest uncertainty (Supplementary Figure 39A). Regions with greater RNA-imaging discrepancies showed higher uncertainty (Supplementary Figure 39B). These uncertain regions also displayed higher reconstruction RMSE and lower Jaccard similarity, indicating that biological differences between RNA and imaging modalities contribute to model uncertainty when aligning the latent space (Supplementary Figures 39C - 39D). The increase in RMSE suggests greater difficulty in reconstructing features due to modality-specific discrepancies, while lower Jaccard similarity suggests that RNA and imaging features capture distinct spatial structures in these regions. Thus, we interpret SCIGMA’s uncertainty estimates as a reflection of biological variation, highlighting the challenges of multi-modal representation alignment in heterogeneous tumor microenvironments.

We next analyzed the mouse small intestine dataset, which features well-defined tissue components such as mucosa, epithelium, and immune compartments^100^ (Figure 5G). From the individual modalities, both RNA expression and imaging features captured the major intestinal structures, including the mucosa, epithelium, and lumen (Figure 5H). However, integration with SCIGMA further clarified these spatial domains, enabling improved resolution of distinct substructures and detecting additional layers within the tissue architecture (Figure 5H). Upon closer inspection, SCIGMA identified subregions within the intestinal layers, including fine-grained segmentation of the villi, lamina propria, and crypt regions (Figure 5H, Supplementary Figure 40). Domain-specific DE analysis further validated these spatial domains (Supplementary Figure 41). For example, cluster 13 exhibited high expression of metabolism marker genes *Enpp7, Fabp1*, and *Lct*^101-103^, suggesting that this domain is the villus epithelium (Figure 5I, Supplementary Figure 42). Cluster 6, enriched with immune marker genes such as *Jchain* and *Igha*^96,104^, represented the lamina propria, a region known for hosting immune cells that maintain gut homeostasis and respond to pathogens (Figure 5I, Supplementary Figure 42). Additionally, clusters 7 and 15 corresponded to the intestinal crypt region, characterized by Paneth cell marker genes (*Lyz1* and *Ang4*)^105,106^ and intestinal stem cell markers (*Olfm4*)^107^ (Supplementary Figure 42). GSEA further validated SCIGMA’s identified spatial clusters. For example, cluster 13 was enriched in pathways related to small molecule transports, membrane communication, and metabolism, supporting its identification as the villus epithelium (Figure 5J). Full GSEA results are provided in Supplementary Figure 43. Cluster relationships were further examined using PAGA and UMAP (Figure 5K). In UMAP space, clusters 7 and 15 were positioned closely together, reflecting shared immune-related functions. However, their separation in PAGA highlighted spatial differences that correspond to the distinct layering and density differences within the tissue (Figure 5H, Supplementary Figure 40), suggesting their different respective roles in the intestinal crypt and broader immune response. Modality-specific weights learned by SCIGMA indicated stronger contributions from the RNA modality, consistent with RNA features showing clearer spatial patterns (Supplementary Figure 44). Consistent with previous datasets, SCIGMA effectively reconstructed and enhanced the sparse RNA modality, revealing more defined spatial patterns of RNA markers (Figure 5L, Supplementary Figure 45). For example, SCIGMA successfully denoised key markers *Apoa4*^*108*^ and *Enpp7*, which are highly expressed in the enterocyte (cluster 18) and villus epithelium regions (cluster 13), respectively. Moreover, we examined the uncertainty estimates (Supplementary Figure 46), which were consistent with the CRC dataset, where high-uncertain regions have greater discrepancies between modalities, higher reconstruction RMSE, and lower Jaccard similarity. In the small intestine dataset, uncertainty was primarily concentrated in the epithelial villi. These findings further suggest that biological variation between RNA and imaging modalities contributes to model uncertainty in aligning the latent space.

### Spatial-Mux-Seq mouse brain data with five modalities

Finally, to evaluate SCIGMA’s ability to integrate more than two modalities, we applied it to a dataset generated by the recent Spatial-Mux-Seq platform^20^. This assay enables simultaneous profiling of up to five omics modalities from the same tissue section. Specifically, we analyzed a 5-month-old mouse brain dataset that measured five modalities including RNA, protein, chromatin accessibility and two histone modifications (H3K27ac and H3K27me3). Anatomical annotations from the Allen Brain Atlas (Figure 2A) served as a reference, encompassing key structures such as the cortex layers, genu of the corpus callosum (CCG), caudoputamen (CP), nucleus accumbens (ACB), lateral septal nucleus (LSR), and anterior commissure (ACO). Clustering on individual modality features revealed that the RNA and protein modalities captured distinct spatial patterns corresponding to different mouse brain regions, while ATAC and histone modifications contain high levels of noise (Figure 6A). For example, the RNA modality identified part of the cortex layers and CCG, whereas the protein modality primarily captured the CCG but failed to differentiate the cortex layers. In contrast, the ATAC and histone modifications modalities were noisy and did not distinctly identify any annotated domains, aside from some noisy cortex layers and the CP region. To analyze this dataset, we extended SCIGMA to accommodate *k* modalities, with *k* = 5 in this case (Supplementary Note 1). SCIGMA effectively identified spatial domains that closely aligned with the atlas annotations, capturing structures such as the cortex layers, CCG (cluster 9), CP (cluster 5, 11), ACB (cluster 2), LSR (cluster 1), and ACO (cluster 10) (Figure 6A, Supplementary Figure 47).

**Figure 6.**
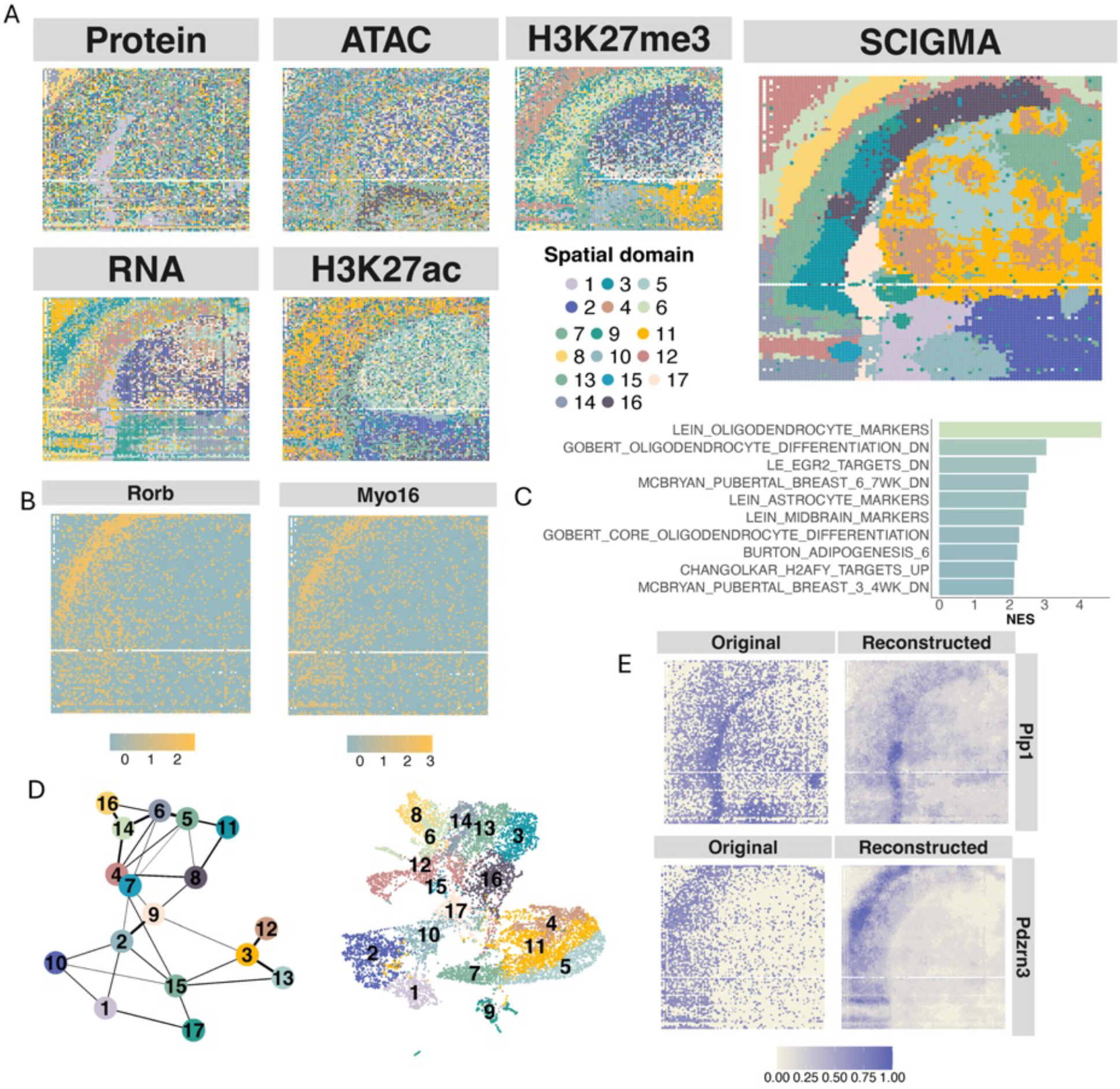
Integration and analysis of Spatial-Mux-Seq mouse brain dataset with five modalities. (**A**) Spatial clusters identified by single-modality, and SCIGMA using integrated multi-modal features, capturing distinct anatomical domains within the mouse brain. (**B**) Spatial visualization of key marker genes associated with cortex layer 4, highlighting spatial patterns detected by SCIGMA. (**C**) Bar plot of the normalized enrichment scores (NES) for the top 10 enriched gene sets in the corpus callosum (CCG) region, identified via GSEA, highlighting biological processes associated with myelination and oligodendrocyte function. (**D**) PAGA graph and UMAP embeddings derived from SCIGMA’s joint representation, showing inferred relationships between spatial domains and capturing their spatial organization within the mouse brain. (**E**) Spatial distribution of key marker features for cortex and CCG regions in SCIGMA’s reconstructed gene expression data.

To validate SCIGMA-detected spatial domains, we performed downstream analysis. In cluster 1, which might correspond to one of the cortex layers, domain-specific DE analysis revealed high expression of cortex markers such as *Rorb, Myo16*, and *Cpne9* ^109-111^(Supplementary Figures 48 - 49). Visualization of these genes showed strong concordance with the SCIGMA-identified cluster. In particular, *Rorb* is a well-known marker of cortex layer 4^109^ (cluster 1), suggesting that *Myo16*, and *Cpne9* may also serve as potential markers for this region (Figure 6B, Supplementary Figure 49). Similarly, cluster 9 showed high expression of myelination and oligodendrocyte marker genes, including *Mobp, Mal*, and *Mog* ^32,112,113^ (Supplementary Figure 49), supporting its correspondence to the CCG region—a region densely populated with oligodendrocytes responsible for producing myelin^114^. GSEA analysis (full results in Supplementary Figure 50) further confirmed that cluster 9 was enriched in pathways related to oligodendrocytes, consistent with the known biology of the CCG. This functional enrichment provides additional evidence supporting the identification of cluster 9 as the CCG domain (Figure 6C). Furthermore, PAGA and UMAP analyses of SCIGMA’s learned representations demonstrated strong agreement with biological organization. For example, we observed a clear progression of cortex layers (cluster 7 corresponding to L2/3, cluster 1 to L4, cluster 2 and 4 to L5, and cluster 6 and 12 to L6) and relationships among the CCG (cluster 9), CP (cluster 5,11), and LSR regions (cluster 1) (Figure 6D). Modality specific weights show that SCIGMA placed greater emphasis on the RNA modality across most domains, except for the CP region, where contributions from all modalities were more balanced (Supplementary Figure 51). This aligns with the individual modality clustering results, where RNA shows clearest spatial patterns in the cortex, CCG, and ACB regions, while all modalities capture the CP region fairly well.

As the number of modalities increased, the reconstruction loss shifted toward learning modality-specific embeddings rather than direct reconstructions, reflecting the challenge of aligning highly diverse data. Despite this, SCIGMA maintained strong reconstruction and denoising capabilities for key markers. For example, SCIGMA effectively reconstructed and denoised marker genes associated with cortex layers and the CCG region (Figure 6E). More examples of the reconstruction of additional genes and other modalities are provided in Supplementary Figure 52. Overall, the reconstructed profiles displayed stronger spatial concordance with their respective domains, demonstrating SCIGMA’s robustness in handling multiple modalities while mitigating technical variation and preserving biological insights. We also examined SCIGMA’s uncertainty estimates to assess how biological and/or technical variations influence model confidence when integrating multiple modalities (Supplementary Figure 53). Spatial mapping of uncertainty values revealed the highest uncertainty in the cortex regions (Supplementary Figures 53A). To investigate potential sources of uncertainty, we focused on discrepancies between RNA and protein as a representative cross-modality comparison, as SCIGMA’s modality weights indicated that RNA was the dominant contributor to spatial domain identification, while protein contributed notably in a subset of domains (Figure 6A, Supplementary Figure 51). We found that RNA-protein discrepancies were not significantly different between certain and uncertain regions (Supplementary Figure 53B), suggesting that direct conflict in feature signal between modalities are not the primary driver of uncertainty. However, RNA reconstruction RMSE was higher in uncertain regions, while protein reconstruction RMSE was lower, indicating that the model has greater difficulty reconstructing RNA features in uncertain areas, while protein features are more easily reconstructed regardless of uncertainty level. This pattern may reflect the relative sparsity and complexity of transcriptomic signals compared to more stable and abundant protein signals at the same spatial locations. Meanwhile, Jaccard similarity values were close to zero for both RNA and protein across both certain and uncertain regions, highlighting that SCIGMA does not explicitly enforce spatial coherence between the modalities during integration due to the biological and technical differences between all five of the modalities resulting in fundamentally distinct spatial patterns (Supplementary Figures 53C-53D). Together, these findings suggest that SCIGMA’s uncertainty estimates reflect a combination of factors, including the greater sparsity and biological complexity of RNA expression, technical noise in spatial transcriptomics, and the intrinsic challenge of integrating diverse modalities with differing spatial resolutions and signal characteristics.

In addition to the above datasets, we demonstrated SCIGMA’s broad applicability across various technology platforms and multi-omics assays. These included three datasets from CUT&Tag-seq^16^ (RNA and histone expression) (Supplementary Figures 54-56), one dataset from Stereo-CITE-seq^115^ (RNA and protein expression) (Supplementary Figure 57), two datasets from 10x Xenium Breast cancer^19^ (RNA and imaging) (Supplementary Figure 58), and six datasets from spatial metabolomics platforms^116^ (RNA and mass spectrometry imaging) (Supplementary Figure 59). SCIGMA’s scalable alignment framework consistently enables the integration and analysis of complex multi-omics data, overcoming challenges posed by modality-specific noise, sparsity, and high dimensionality.

## Discussion

We present SCIGMA, a graph neural network-based framework for spatial multi-omics integration that leverages uncertainty-aware contrastive learning. By incorporating graph attention networks (GATs), SCIGMA captures both local and global spatial structures in tissue organization. A cross-attention fusion layer integrates information across modalities, ensuring alignment while preserving modality-specific signals. To improve interpretability, SCIGMA explicitly models uncertainty, identifying regions where integrating different modalities is more challenging due to biological heterogeneity or technical variation. SCIGMA is modular and scalable, capable of integrating more than two modalities and efficiently processing datasets exceeding a million spatial locations on a single GPU. Through evaluations across seven datasets covering diverse technologies, tissue types, and multi-omics platforms, we show that SCIGMA enables spatial domain detection, trajectory analysis, and denoising, while providing uncertainty estimates to assess integration confidence.

SCIGMA introduces several key advantages over existing methods. First, its modality-specific weighting mechanism ensures that the most informative molecular layers contribute to the integration process, enhancing biological relevance in spatial domain detection. Second, SCIGMA’s scalability allows it to process high-dimensional, high-resolution datasets, such as 10x VisiumHD (at 2 *μm*) and Xenium Prime, which exceed the computational limits of many existing spatial multi-omics tools. Third, SCIGMA supports flexible integration of up to five molecular modalities, as demonstrated with Spatial-Mux-Seq mouse brain data, providing a comprehensive view of tissue organization across transcriptomics, proteomics, chromatin accessibility, and histone modifications. Fourth, SCIGMA’s uncertainty quantification identifies regions where multi-modal alignment is less reliable, capturing both biological heterogeneity and technical variation. Across diverse datasets, uncertain regions consistently corresponded to areas of high tissue complexity, such as heterogeneous tumors, immune compartments, and layered brain structures, providing valuable insights into the confidence of spatial domain assignments. Finally, SCIGMA outperforms existing methods in spatial domain detection, reconstruction accuracy, and reproducibility, while being the only method capable of handling all current spatial multi-omics platforms. Together, these features make SCIGMA a robust and generalizable framework for spatial multi-omics data integration and analysis.

There are several future directions for SCIGMA. First, while SCIGMA currently combines the feature and spatial graph through a simple union, incorporating a weighted combination of the two graphs would further benefit the latent representation learning. Second, SCIGMA’s flexible framework makes it well-suited for 3D spatial multi-omics, where data from adjacent tissue slices or biological replicates could be jointly analyzed to identify shared and differential molecular features. This extension would allow for more generalizable latent representations, further improving cross-sample comparisons. Third, SCIGMA’s uncertainty estimates could be expanded to provide modality-specific confidence scores for each spatial location, offering fine-grained uncertainty assessments that distinguish biological heterogeneity from technical inconsistencies. Finally, integrating uncertainty estimates into downstream analyses remains an open question in spatial multi-omics integration. Future work could explore leveraging uncertainty values in spatial clustering, differential expression analysis, or predictive modeling to enhance biological interpretability and decision-making in tissue profiling. However, we emphasize caution in over-reliance on uncertainty estimates as ground truth indicators of uncertainty in the data or model predictions, as our analyses reveal that SCIGMA’s uncertainty values reflect a complex mixture of biological and technical variation (Supplementary Note 5).

## Methods and Materials

### SCIGMA Overview

We provide a brief overview of SCIGMA (Figure 1) with more technical details in Supplementary Note 1. Briefly, SCIGMA is a multi-view graph-based deep learning model optimized via an unsupervised contrastive learning framework. Given a multi-omics dataset with *M* different modalities, each with distinct feature set 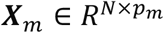, where *N* is the number of spatial locations in the tissue slice, and *p*_*m*_ is the number of features in modality *m*, where 1 ≤ *m* ≤ *M*. For each spatial location 2, the modality-specific feature data is denoted as *X*_*im*_. We denote the 2-dimensional spatial coordinate matrix as ***S***, where ***S*** ∈ *R*^*m*×*k*^. To our knowledge, existing spatial multi-modal technologies can measure up to five different modalities for the same spatial location^20^, with, *M* ≤ 5. For example, in spatial-ATAC-RNA-seq^16^, *p*_1_ and *p*_2_ correspond to genes and chromatin regions, respectively, while in Spatial-CITE-seq^18^ and 10x VisiumHD^19^, they refer to genes co-profiled with proteins and H&E image pixels, respectively. Details of the multimodal measurements from the state-of-the-art spatial multi-modal technologies are listed in Supplementary Table 1. Without loss of generality, we use *M* = 2 as an example to illustrate our model’s ability to jointly integrate two modalities, reflecting the common configuration in most spatial multimodal platforms. SCIGMA effectively projects modality-specific biological information into a meaningful shared latent representation, preserving the unique characteristics of each modality through the following framework:

1. Graph construction: SCIGMA constructs a ‘combined’ graph for each modality. This graph integrates a shared spatial graph, representing the physical locations shared across modalities, with a feature graph that encapsulates the distinct features (*p*_*m*_) of each modality *m*. This integrated graph structure allows for a nuanced representation of both spatial and modality-specific feature data.
2. Modality-specific encoding: Using graph attention layers, a modality-specific encoder is created. These layers are adept at focusing on relevant features while filtering out noise, thereby ensuring that the encoder captures the essential characteristics of each modality. This step translates the high-dimensional biological information into compact modality-specific embedding spaces, enhancing the model’s ability to manage complex data structures efficiently.
3. Integration layer: A cross-attention layer is employed to fuse the modality-specific latent representations into a cohesive joint latent space. This integration is crucial as it harmonizes the embeddings from different modalities, ensuring that inter-modality relationships are maintained and leveraged for subsequent analyses.
4. Decoding and reconstruction: The integrated latent representation is then decoded back into the individual modality feature spaces using a linear layer combined with further graph attention layers. This decoder structure not only reconstructs the original modality features from the joint latent space but also refines the feature representations, making them more robust and informative.

### Feature Matrix Construction

For each modality, we preprocessed the associated feature matrix (Supplementary Note 3) and applied principal component analysis (PCA) to generate lower-dimensional representations. This preprocessing step serves two main purposes: (1) PCA embeddings capture key biological signals while reducing noise that could otherwise propagate through the model, and (2) dimensionality reduction improves model scalability as the number of modalities increases. Combined with our batching strategy during training (Supplementary Note 1), this approach allows SCIGMA to efficiently scale to datasets with millions of spatial locations (Supplementary Table 1). For non-imaging modalities, we applied modality-specific preprocessing before performing PCA. Detailed preprocessing steps for each dataset are provided in Supplementary Note 3. For morphology images, we followed previous studies^117,118^ to extract meaningful imaging features by segmenting the image based on the coordinates of each spatial location and using a pretrained ResNet^119^ model to generate deep features. Specific details on imaging feature extraction for each dataset are provided in Supplementary Note 3.

### Combined Graph Construction

For each modality, we construct a ‘combined’ graph using a shared spatial graph and a feature graph based on the modality’s features. Previous studies^24^, including our work on CARD^24^ have demonstrated the importance of integrating spatial information for studying cellular compositions in spatial transcriptomics. Given that physically adjacent spatial locations typically contain similar cell types or states, we first construct an undirected spatial graph using the shared spatial location information denoted as *G*^*spa*^ = (***V, E***), where ***V*** represents the set of *N* vertices corresponding to spatial locations and ***E*** represents the set of edges connecting neighboring spatial locations. Let ***A***^(*spa*)^ denotes the spatial adjacency matrix of 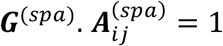 if and only if the location *i* and *j* are physically neighbors and 0 otherwise. For each spatial location, we select the top *k* nearest neighbors, where *k* is a hyperparameter adjusted based on the technology. In our analyses, we set *k* = 6 for smaller datasets (under 8k locations) and *k* = 18 for larger datasets (8k locations or more), optimized via grid search hyperparameter tuning (Supplementary Note 1, Supplementary Table 4). Since this graph is built using shared spatial coordinates, the adjacency matrix is the same across modalities.

To accommodate scenarios where locations might share similar biological information but are spatially distant, we construct a feature graph for each modality, denoted as 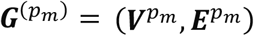, where *m* ∈ {1,2,,, *M*} represents the *m*-th modality. Here, 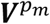 represents the set of *N* vertices for modality *m*, and 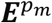 represents the edges connecting spatial locations in modality *m*. This graph is constructed using a k-nearest neighbors (KNN) algorithm applied to the PCA embeddings of the features for each modality. Let 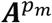 denote the adjacency matrix of 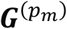. Where 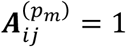 if and only if locations 2 and = are nearest neighbors; else, 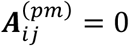. For each spatial location, we select the top 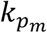 nearest neighbors, where 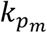 is a hyperparameter that can be adjusted depending on the dataset. For our analyses, we chose 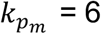 for all datasets, optimized via grid search hyperparameter tuning (Supplementary Note 1, Supplementary Table 4). Since this graph is built using modality-specific data, each modality has its own distinct feature graph.

Given that we aim to incorporate both spatial and feature-based relationships, we construct a combined graph, denoted as 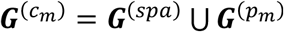, where *m* ∈ {1,2, …, *M*} represents the *m*-th modality. By taking the union of the spatial neighbors and the feature neighbors, we integrate both spatial and feature-based relationships while collapsing any redundant information. For example, if a given spatial location 2 has a neighboring location = in both the spatial and feature graphs, the combined graph retains only one edge between them. This approach preserves biologically relevant connections while reducing redundant information, which aids in efficient model training. By this construction, each modality has its own combined graph 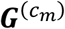.

### Graph Attention Layers

To capture the unique feature distribution of each modality, we use graph attention networks (GATs)^120^, a type of graph-based deep learning networks, as the basis of the modality-specific encoders. Unlike graph convolutional networks (GCNs), which aggregate information via a normalized sum and can lead to “over-smoothing” by averaging neighboring node features^121^, GATs assign weights to neighboring nodes based on attention scores. This approach allows for a more nuanced capture of local patterns while still accommodating global trends, ensuring that key local variations are preserved in each modality’s representation.

#### Graph attention network

We define the graph attention layer, which takes the feature matrix *X*^(*m*)^ for a given modality *m* (either as PCA input or as low-dimensional embedding from previous layers). Each spatial location/node undergoes a linear transformation parameterized by a trainable weight matrix ***W***. This transformation utilizes neighborhood information from the combined graph 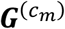 to compute attention scores, including self-attention, via a scoring function *a*(·) as follows:

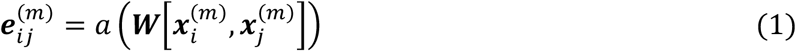

where 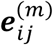 represents the importance of spatial location *j* to *i* for modality *m*. We inject the graph structure into this standard attention by restricting 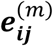 calculations to spatial location/node *j* within the neighborhood 𝒩_𝒾_ of location/node *i*. Taking the softmax of these scores yields attention coefficients:

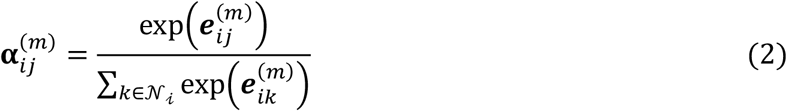

The final node representation is obtained by using the attention coefficients as weights in a linear combination of the features from neighboring nodes *N*_*i*_, which produce the output features for spatial location *i*:

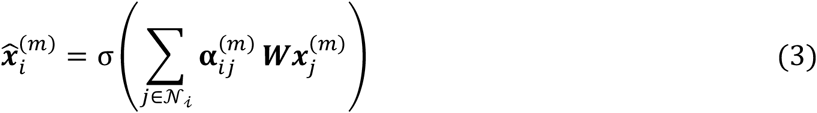

where σ (·) is a nonlinear activation function.

To enhance the model’s learning stability and capture diverse perspectives, we employ a multi-head attention framework^120^, where *H* independent heads perform the attention transformation and their features are concatenated,

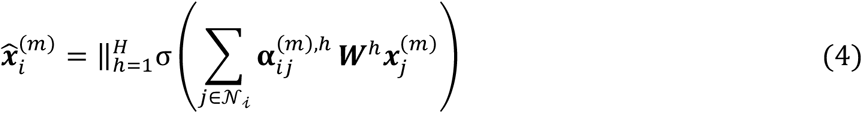

Each modality has an encoder backbone, which consists of a series of GATs. Each encoder takes the preprocessed data input *X*^(*m*)^ (Details of the data preprocessing see Supplementary Note 3) and combined graph 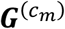 as inputs, and outputs a modality-specific latent representation, denoted ***Y***^(*m*)^.

#### Attention layer

Each modality-specific low-dimensional embedding represents a distinct view of the tissue sample, capturing both shared and unique information across modalities. To integrate these perspectives into a joint latent space, we employ the Cross Attention Mechanism^122^. This mechanism allows SCIGMA to effectively discern and prioritize relevant information from each modality for each location, efficiently filtering out conflicting information or noise. By applying cross-attention to manage the modality-specific embeddings, we enhance the model’s capacity for robust and precise data integration.

Specifically, for a given spatial location *i* in modality *m*, with its corresponding embedding 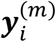, the score function for cross-attention is defined and implemented as

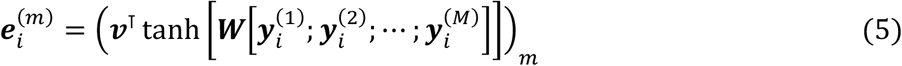

where ***v*** is a trainable weight vector, ***W*** is a trainable weight matrix, ***M*** represents the total number of modalities being integrated, the ‘;’ operator denotes concatenation, and 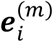 is the attention score value for spatial location *i* in modality *m*. Taking the softmax of the scores transforms them into attention coefficients, scaled as probabilities:

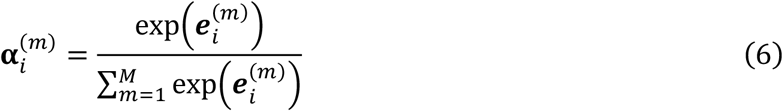

Finally, the joint latent representation ***Z*** is computed as a weighted sum of modality-specific embeddings with the attention coefficients as the weights:

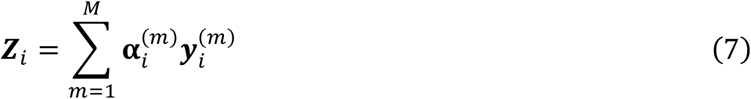

### Training Framework

#### Graph sampling

With the rapid advancements of spatial omics technologies, multimodal datasets have grown significantly, now capturing gene expression at millions of spatial locations per tissue slice (e.g., 10x VisiumHD^19^). This increase in data scale leads an exponential growth in graph size constructed from these datasets, resulting in higher computational demands and memory constraints. To address these challenges, we utilize graph sampling techniques^123,124^ to train on batches of the combined graphs ***G***^(*cm*)^. Specifically, we employ forest fire sampling^124^, which allows efficient sampling while preserving the graph’s structural integrity. The process begins with a randomly selected seed node, from which edges and adjacent nodes are iteratively “burned,” meaning they are included in the sampled subgraph. Each burned node, in turn, has a probability of burning its own edges. This process ensures that the final sampled subgraph retains a representative structure of the original graph. By setting a batch size that determines the number of nodes in each subgraph (see Supplementary Note 1), SCIGMA can scale efficiently even for datasets containing millions of spatial locations, making it feasible to train on a standard 24Gb Nvidia GPU without exceeding memory constraints.

#### Loss function

SCIGMA aims to 1) accurately learn the feature distribution of each modality and 2) integrate them into a biologically meaningful joint representation. Therefore, the loss function is carefully formulated to optimize both objectives.

To capture the feature distribution of each modality *m*, SCIGMA learns to reconstruct the input features *X*^(***m***)^ by transforming the joint representation ***Z*** via a decoder architecture. Each modality-specific decoder mirrors its corresponding encoder’s architecture and applies a linear transformation to ***Z*** to yield reconstructed modality-specific feature matrices 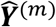. The output of these decoders are the reconstructed feature matrices for each modality *m*, 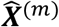. For these reconstructions, we apply *L*_2_ or mean squared error (MSE) loss to ensure the accurate recovery of the model-specific features. The reconstruction loss function is defined as follows:

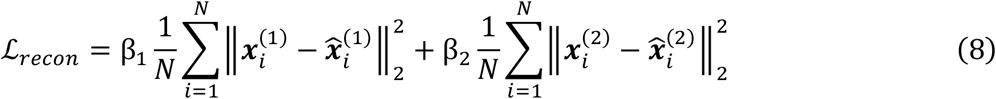

Here, 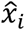 represents the reconstructed features for modality *m* at spatial location *i*. The coefficients β_1_ and β_2_ are hyperparameters representing loss weighting for balancing the contributions of each loss term, which are determined through hyperparameters tuning (Supplementary Note 1, Supplementary Table 5). Minimizing ℒ_*recon*_ ensures that SCIGMA effectively learns the feature distribution for each modality.

To integrate multiple modalities, we implement a variant of contrastive loss known as CrossCLR^125^ (Supplementary Note 1), which aligns joint and modality-specific representations while preserving each modality’s unique feature distribution. Unlike standard contrastive learning, which focuses only on pushing similar samples together and dissimilar ones apart in a shared embedding space, CrossCLR defines negative samples both within and across different latent spaces. This design is particularly well-suited for integrating heterogeneous modalities, such as RNA, protein, ATAC-seq, imaging, and etc. Specifically, between the joint representation ***Z*** and each modality-specific representation ***Y***^(*m*)^, we apply a separate CrossCLR loss. Thus, SCIGMA learns a joint representation that aligns closely with each modality-specific representation while maintaining the distinct feature distributions of individual modalities. Unlike conventional approaches that enforces similarity across modalities, SCIGMA allows each modality to retain its unique biological signals, ensuring that the integrated representation captures both shared and modality-specific variations. Let ***Z***_*i*_ denotes the joint representation at spatial location *i*, and *y*_*i*_ the modality-specific representation for modality *m* ∈ {1, …, 2}. Then the following function measures the similarity of the two embeddings:

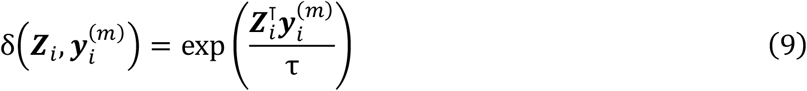

Where 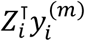 represents the dot product, capturing alignment between the vectors in the joint and modality-specific embedding space, and *τ* is a temperature hyperparameter controlling the sensitivity of the alignment. The CrossCLR loss between the joint representation and the modality specific representation at spatial location 2 is then defined as:

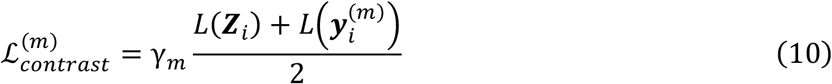

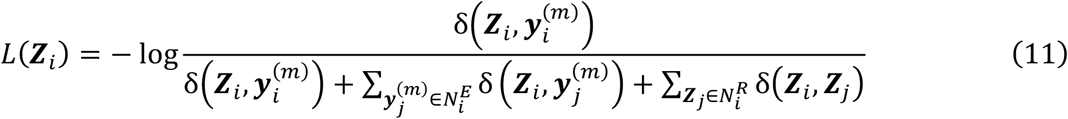

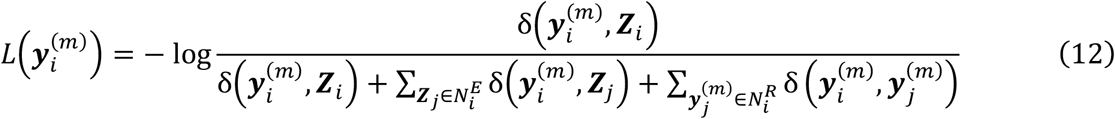

Here, 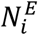 denotes the set of inter-modality negative samples and 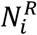 denotes the set of intra-modality negative samples, and γ_*m*_ is a hyperparameter weighting the contribution of contrastive loss (hyperparameter tuning detailed in Supplementary Note 1 and Supplementary Table 5). A positive pair is defined as the joint and modality-specific latent representations corresponding to the same spatial location. Negative samples are defined as representations from all other spatial locations, both within the same modality (intra-modality) and across different modalities (inter-modality). Based on this setup, we define the overall loss function of SCIGMA when *m* ∈ {1,2} as

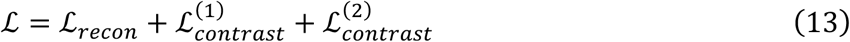

Equation (13) presents a representative loss function for integrating two modalities. However, SCIGMA’s modeling framework is designed to be both robust and flexible, enabling it to generalize seamlessly to datasets with more than two modalities. For instance, SCIGMA can effectively model the five modalities generated by the recent Spatial-MUX-Seq technology^20^, which include modalities such as histone, chromatin accessibility, and protein. Details on the generalization to multiple modalities (more than two) are provided in the Supplementary Note 1. This comprehensive loss function allows SCIGMA to learn a biologically meaningful joint representation that aligns closely with each modality-specific representation while preserving the unique feature distributions of individual modalities.

#### Uncertainty modeling

In genomics, current deep learning models often lack robust mechanisms for quantifying or accounting for uncertainty, which is crucial for interpreting outputs, particularly when integrating diverse modalities into a joint representation^26,126,127^. While Variational Autoencoders (VAEs) attempt to incorporate uncertainty by modeling the distribution of latent variables, their effectiveness hinges on accurately specified prior and posterior distributions. This dependency can lead to significant drawbacks when prior knowledge is inaccurate or unavailable, as VAEs can struggle with noise, artifacts, and inadequate representation of the underlying data distribution, potentially leading to misleading uncertainty estimates and overfitting to poor-quality data^128-130^. Our approach addresses this deficiency by explicitly inferring the parameter *τ* in our contrastive loss formulation, as in equation (9), which modulates the model’s sensitivity to differences between embeddings. Specifically, smaller values of *τ* amplify these differences, resulting in larger gradients, while larger values make the model less sensitive to such discrepancies. Following^131^, we dynamically learn *τ* at each spatial location as a measure of model uncertainty of the joint representation for that sample. At each spatial location, the value of *τ* indicates the model’s ability to learn aligned representations for a positive pair according to the contrastive loss formulation, thus representing the model’s uncertainty or certainty in integrating a particular spatial location’s modality-specific data. Larger values of *τ* suggest greater uncertainty, implying that the optimization of the objective function is less responsive to differences in the embeddings at that location and thus accommodate larger differences in the embedding spaces. Conversely, smaller values of *τ* indicate smaller uncertainty, as the objective is more sensitive to differences, making the model more likely to learn well-aligned embeddings at that spatial location. In this framework, we directly learn a data-driven uncertainty metric *τ* that dynamically adjusts based on the data, rather than relying on fixed assumptions about data quality. This adaptive approach enables robust and context-aware uncertainty estimation across different modalities and spatial locations.

### Compared Methods for spatial multi-modal integration

To benchmark SCIGMA against state-of-the-art methods, we evaluated its performance on the datasets that the other methods can deal with, including mouse brain spatial-RNA-ATAC dataset and the mouse spleen SPOTS dataset. Specifically, we compared SCIGMA to Seurat (version v5), MOFA+ (version 1.13.0), MultiVI (version v1), and SpatialGlue (version v1). For each method, we followed their GitHub tutorials with default settings, training and running them for various downstream analyses. This range of methods was chosen to evaluate SCIGMA across different analytical approaches, highlighting its versatility in handling both single-cell and spatial omics datasets and its specialized capabilities for integrating complex multimodal data. Comparing SCIGMA with these diverse methodologies also helps demonstrate its robustness against traditional and spatial-specific challenges, positioning it within the current landscape of genomic tools. However, for other large-scale datasets, due to computational limitations (e.g., 48 hours of runtime, 400Gb of CPU memory, 24Gb GPU memory) and the inability of some methods to handle complex modalities such as imaging (details see Supplementary Table 2), we focused our analysis exclusively on SCIGMA.

### Real Data Analysis

We applied SCIGMA to nineteen different spatial multi-omics datasets collected via 9 different techniques and resolutions (Supplementary Table 1). Details of preprocessing are in Supplementary Note 3. Benchmarked model details are in Supplementary Note 4. Runtime for training SCIGMA is provided in Supplementary Table 6. Additional details for each downstream analysis are in the Supplementary Note 5.

#### Spatial domain detection

For each dataset, SCIGMA learns a joint representation from the expression data of different modalities. We then applied the *mclust* algorithm^132^ to the learned joint representation to identify spatial clusters. For each dataset, we used the number of clusters determined from the analyses as a starting point (if available) to test different numbers of clusters to select the clustering that best captured biological structures (Supplementary Note 5). To evaluate integration performance, we assessed four quantitative metrics, each capturing a different aspect of performance: (1) Moran’s I score, which evaluates whether the pattern of the joint representations is spatially clustered, dispersed or random. A higher score indicates stronger spatial clustering in the data. (2) Jaccard similarity, which measures the similarity between each modality-specific representation and the learned joint representation. A higher value indicates better preservation of modality-specific spatial and feature relationships in the integrated representation. (3) Pairwise ARI or NMI, which assess the stability of deep learning models by performing pairwise comparisons of clustering results across 20 random seeds, given that deep learning methods often exhibit variability. A higher value indicates more consistent clustering results from training on different seeds. Details about these metrics are provided in Supplementary Note 5.

#### Differential gene expression and differential protein expression analysis

For differential gene expression (DEG) and differential protein expression (DEP), we followed standardized tutorial from the Seurat package^21^. For either gene expression, and protein expression, we first normalized and scaled the data, and then used the ‘FindAllMarkers’ function to identify either DEGs or DEPs. Specifically, we used the following parameter settings: logfc. threshold = 0.1, min.pct = 0.1 and ‘wilcoxon’ test for all datasets except for the SPOTS and VisiumHD datasets. As the VisiumHD datasets were especially sparse and the SPOTS dataset had few samples, we used the following parameter settings for ‘FindAllMarkers’: logfc.threshold = 0.0, min.pct = 0.0 and ‘wilcoxon’ test.

#### Differential peak analysis

In the Spatial-ATAC-RNA-seq data, we used the ArchR package (v1.0.2)^133^ and followed their tutorial to identify the differentially expressed peaks. Specifically, we first created arrow files using the parameters: minFrags = 0, maxFrags = 1e+07, tile_size = 5000, addGeneScoreMat = True, and ‘the mm10’ genome. Dimensionality reduction was performed via IterativeLSI with dims = 1:30. We defined the peak set found in the anndatas if the dataset provided them, or generated them using ‘addReproduciblePeakSet’. The differentially expressed peaks were then identified for SCIGMA’s clusters in the ‘PeakMatrix’ with the ‘getMarkerFeatures’ function. Marker genes with differential gene scores were also computed from the ‘GeneScoreMatrix’ using the same function. Finally, we computed the linkage between genes and peaks using the ‘addPeak2GeneLinks’ function with the ‘Iterative LSI’ reductions and ‘GeneScoreMatrix’ values. We ran the ‘addPeak2GeneLinks’ function with the following settings: corCutOff = 0.45 and resolution = 1,000. To visualize the correlation between peaks and genes, we used the ‘plotPeak2GeneHeatmap’ function to plot the peak-to-gene links heatmap.

#### Gene set enrichment analysis

We performed gene set enrichment analysis (GSEA) on DE genes using the python library ‘gseapy’ with the corresponding GSEA function ‘prerank’^134^. For mouse datasets, we used the MSigDB Mouse curated gene sets, and for the human datasets, we used the MSigDB Human curated gene sets^135^. Specifically, we followed their tutorial and set the following parameters for ‘prerank. For the two modality mouse datasets, we set min_size = 5, max_size = 500, permutation_num=1000. For the two modality human datasets, we set min_size = 5, max_size = 300, permutation_num=1000. For the five-modality mouse dataset, we set min_size = 2, max_size = 1000, permutation_num=1000. The resulting GSEA outputs were filtered with the following to identify upregulated gene sets: for the mouse datasets, NES >=0 and FDR <= 0.25; for the human datasets, NES >= 1 and FDR <= 0.25, following the recommended thresholds.

#### PAGA analysis

Partition-based graph abstraction (PAGA) generates a graph-like manifold of clusters, where vertices represent clusters and edges capture their similarity^136^. Thus, it preserves the underlying topology of the data to provide interpretable trajectory or progression of clusters. Applying PAGA to SCIGMA-generated embeddings allow us to infer relationships between biological structures and assess their alignment with expected biology. Specifically, we performed PAGA analysis using scanpy package in Python^137^. From the PAGA representation, we examined the manifold structures in the integrated representation and interpreted relationships between biological structures captured by SCIGMA.

#### Reconstruction and denoising

To evaluate SCIGMA’s ability to recover biologically meaningful signals from multimodal data while mitigating technical noise, we reconstructed the original modality-specific feature data from the model’s output. Since SCIGMA takes PCA-transformed features as input, its decoder produces a reconstructed PCA matrix, which can be interpreted as a denoised version of the input. To assess how well SCIGMA preserves biological information, we projected this reconstructed PCA matrix back into the original feature space using an XGBoost regressor^138^. Specifically, we first trained an XGBoost model using the dimensionality reduced matrix (e.g., PCA-transformed features) derived from the original modality feature matrix as input and the corresponding original modality-specific features as output. This model learns to map between the dimensionality reduced features (e.g., PCA-transformed features) and the original feature matrix. We then applied the trained XGBoost regressor to SCIGMA’s reconstructed dimensionality reduced matrix, enabling us to recover an approximation of the original feature matrix from the denoised representation. By comparing these reconstructed features with the measured data in the original feature space, we assessed SCIGMA’s ability to enhance biological signal retention while reducing the impact of technical variations. In the XGBoost model training steps, we systematically tuned the hyperparameters of the XGBoost model, optimizing key parameters such as the number of trees, learning rate, maximum depth, and regularization terms. Details of the hyperparameter tuning process and the final selected values are provided in the Supplementary Note 1 and Supplementary Table 7.

#### Neighborhood enrichment analysis

To assess the spatial relationships between clusters, we calculated neighborhood enrichment scores using the Squidpy package^139^. We first obtained clusters from SCIGMA’s joint representations and used them, along with spatial coordinates, as input for Squidpy’s neighborhood enrichment function nhood_enrichment to evaluate spatial autocorrelations of clusters. Specifically, this function calculates builds a spatial connectivity graph using Delaunay triangulation and calculates an estimate of the association between two clusters. The function outputs a z-score for each pair of clusters indicating whether that cluster pair is over-represented or over-depleted for node-node interactions. Details of the calculation are provided in the Supplementary Note 5.

#### Uncertainty analysis

The uncertainty parameter *τ* in the contrastive loss represents the model’s confidence in aligning embeddings from different modalities. In the biological context, the alignment between modalities is influenced by two sources of variation: biological variation and technical variation. Biological variation arises when different modalities such as RNA expression and chromatin accessibility, capture distinct yet meaningful biological signals. This makes it more difficult for the model to align embeddings, resulting in higher uncertainty. On the other hand, technical variation results from experimental noise or measurement artifacts, leading to differences in feature distributions between modalities that do not necessarily reflect underlying biology. Given the presence of both types of variation in multimodal datasets, we hypothesize that the uncertainty parameter learned by SCIGMA captures these discrepancies, with higher uncertainty values indicating greater difficulty in integrating the modalities. To validate this, we performed downstream analyses to interpret the learned uncertainty values, and determine whether they primarily reflect biological or technical variation. To do so, we first stratified the uncertainty values into different levels (top 1%, 5%, 10%, 20%). Locations with lower-uncertainty values were categorized as certain regions while those with higher uncertainty values as uncertain regions. We then conducted three types of quantitative assessments: (1) we compared the distribution of differences in scaled expression values between different modalities for certain versus uncertain regions. If uncertain regions exhibit greater expression differences compared to certain regions, this suggests that uncertainty primarily modality misalignment, as the magnitude of feature information differs between the modalities. If both uncertain and certain regions show similar expression differences, this indicates that there is a difference in the feature distributions between the modalities. Depending on the spatial distribution of the uncertainty the values and the following analyses, these differences could either reflect biological or technical variation. (2) we calculated SCIGMA’s modality-specific reconstruction error at each spatial location for certain versus uncertain regions. A higher RMSE in uncertain regions suggests higher likelihood that biological variation contributes to uncertainty, as modality-specific differences prevent effective alignment. Lower RMSE in uncertainty regions might indicate that the uncertainty is likely driven by technical variation, as the features remain well-aligned despite model uncertainty. (3) we calculated the Jaccard similarity coefficient between the joint representation and each modality-specific representation for uncertain and certain regions. A lower Jaccard similarity coefficient in uncertainty regions suggest that embeddings fail to preserve biological signals due to modality-specific differences (biological variation). Conversely, a higher Jaccard similarity coefficient in both uncertain and certain regions suggests that uncertainty primarily captures technical variations as the biological features are preserved well after alignment. By performing these analyses, we aimed to determine whether SCIGMA’s uncertainty values predominantly capture biological variation, technical variation, or both. This evaluation informs us about the reliability and provides interpretability of the integrated joint representation, as well as the model’s ability to resolve differences across modalities. Specifically, the differences in scaled expression values across modalities indicate the extent of initial variation in the input features. The reconstruction RMSE reflects confidence in feature alignment, and the Jaccard similarity coefficient measures the alignment of embedding topology. Details of the implementation are provided in the Supplementary Note 5.

### Ablation studies

To systematically evaluate the contribution of individual components and design choices within SCIGMA, we conducted a series of ablation studies. These studies aim to quantify the impact of each component on the model’s overall performance and integration capabilities. Specifically, we evaluated SCIGMA’s modeling design from six distinct perspectives: (1) Contrastive loss: To demonstrate the necessity of contrastive loss for learning well-integrated joint representations, we trained SCIGMA without the contrastive loss component. This helped assess how critical this element is for aligning different modalities within the integrated framework. (2) Spatial Information: To highlight the importance of spatial information, we trained SCIGMA using only feature graph. This variation allowed us to understand the extent to which spatial relations contribute to model accuracy and data integration. (3) Feature graph: To highlight the importance of combined graph, we trained SCIGMA using only spatial graph, testing whether modality-specific feature similarities further enhance integration (4) Graph attention layers: To demonstrate the benefits of using graph attention layers, we substituted GAT with GCN. This change helped to evaluate the impact of different graph processing techniques on the model’s ability to learn from complex structures. (5) CrossCLR loss: To quantify the importance of the CrossCLR loss, we trained SCIGMA with a regular contrastive loss objective, assessing its role in aligning modality-specific and joint representations (6) Uncertainty parameter: Finally, to underscore the importance of learning the uncertainty parameter for training stability and effective integration of heterogeneous data, we trained SCIGMA using vanilla CrossCLR contrastive loss (without uncertainty parameter). This experiment demonstrates that incorporating an uncertainty parameter improves model stability and enhances multimodal integration. These ablation experiments were conducted on the Spatial ATAC-seq dataset, where the biological structure of the P22 mouse brain is well-defined. We evaluated each model variant from different perspectives, including clustering performance, pairwise ARI/NMI, Moran’s I score, Jaccard index, and the RMSE of SCIGMA’s modality-specific reconstruction error. Additional details are provided in Supplementary Note 2.

## Data Availability

We analyzed the following datasets: (1) spatial epigenome-transcriptome P22 mouse brain ATAC datasets (https://web.atlasxomics.com/visualization/Fan/, preprocessed anndatas provided by the SpatialGlue paper https://zenodo.org/records/10362607); (2) SPOTS mouse spleen (accession no. GSE198353, preprocessed anndatas provided by the SpatialGlue paper https://zenodo.org/records/10362607); (3) 10x Xenium Prime 5K Human Ovarian Adenocarcinoma (https://www.10xgenomics.com/datasets/xenium-prime-fresh-frozen-human-ovary); (4) 10x Xenium Prime 5K Human Cervical Cancer (https://www.10xgenomics.com/datasets/xenium-prime-ffpe-human-cervical-cancer); (5) 10x Visium HD Mouse small intestine (https://www.10xgenomics.com/datasets/visium-hd-cytassist-gene-expression-libraries-of-mouse-intestine); (6) 10x Visium HD Human colorectal cancer (https://www.10xgenomics.com/datasets/visium-hd-cytassist-gene-expression-libraries-of-human-crc); (7) Spatial Mux Seq 5 month mouse brain (accession no. GSE263333); (8) Stereo-CITE-seq mouse thymus (preprocessed anndatas provided by SpatialGlue https://zenodo.org/records/10362607); (9) spatial epigenome-transcriptome P22 mouse brain histone datasets (https://web.atlasxomics.com/visualization/Fan/, preprocessed anndatas provided by SpatialGlue https://zenodo.org/records/10362607); (10) 10x Xenium Human BRC (https://www.10xgenomics.com/products/xenium-in-situ/human-breast-dataset-explorer); (11) spatial metabolomics mouse brain (https://data.mendeley.com/datasets/w7nw4km7xd/1).

## Code Availability

The SCIGMA software package and source code have been deposited at https://github.com/YMa-lab/SCIGMA. Scripts used to reproduce all the analysis are also available at the same website.

## Author contributions statement

Y.M. conceived the idea and designed the study. S.C. and Y.M. designed the model. S.C. developed and implemented SCIGMA software. S.C. and Y.M conducted data analysis. S.C. and Y.M. prepared the figures. S.C. and Y.M wrote the manuscript. A.F. provided expert input on the biological interpretation of results and offered detailed review and revisions of the manuscript.

