## Supplementary for "Scalable, Generalizable, and Uncertainty-Aware Integration of Spatial Multi-Omics Across Diverse Modalities and Platforms with SCIGMA"

### Supplementary Figures

**Supplementary Figure 1. Ablation studies performed for assessing individual design choices in SCIGMA.** Figure is displayed next page **(A)** Spatial domain detection results for six ablated SCIGMA models and full version of SCIGMA. We note poor domain detection without contrastive loss or spatial information; over smoothing without feature graph information or GATs; and slightly fragmented domains without incorporating uncertainty estimation. **(B)** Moran's, I result for the ablated models. Each box represents an ablated model, the y-axis represents the metric value. **(C)** Jaccard index results for the ablated models. Each bar represents an ablated model, the y-axis represents the mean Jaccard value, and the colors represents the modality for which the Jaccard index was calculated. (D) Dirichlet energy metric that measures the over smoothness in GNNs, each box represents an ablated model, the y-axis represents the metric value. **(E)** Pairwise ARI, and pairwise NMI results for the ablated models. Each box represents an ablated model, the y-axis represents the metric value. **(F)** RMSE reconstructed feature results for each modality at uncertain spatial locations estimated from the full SCIGMA model. For each modality, the x-axis represents the uncertain location threshold (top k% most uncertain locations, where k = 1, 5, 10, and 20), y-axis represents the RMSE of the reconstructed feature matrix. Details interpretations of the results see Supplementary Note 2.

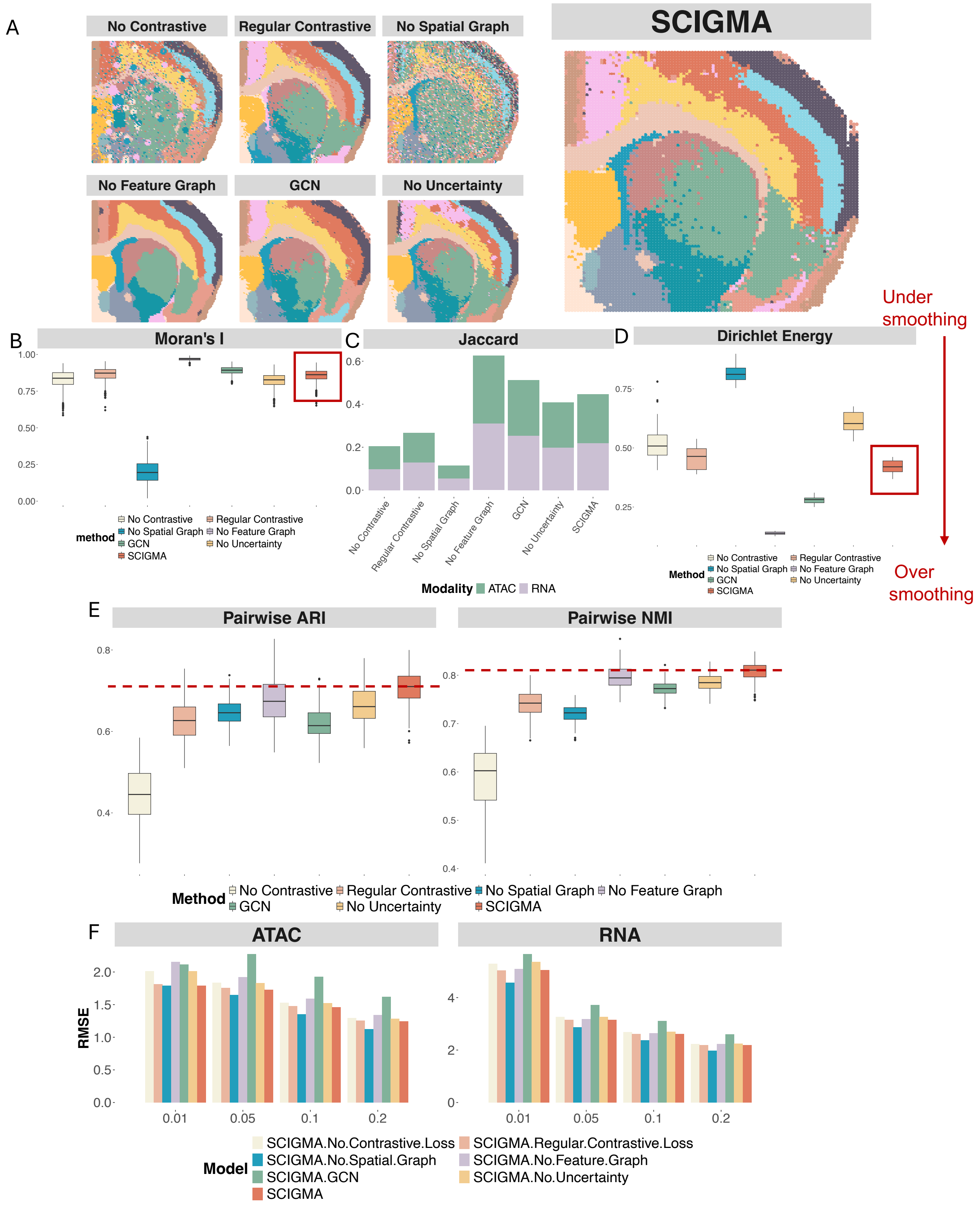
A
No Contrastive
Regular Contrastive
No Spatial Graph
SCIGMA
No Feature Graph
GCN
No Uncertainty
Under smoothing
Over smoothing
B
Moran's I
method
No Contrastive
Regular Contrastive
No Spatial Graph
No Feature Graph
GCN
No Uncertainty
SCIGMA
C
Jaccard
Modality ATAC RNA
D
Dirichlet Energy
Method
E
Pairwise ARI
Pairwise NMI
Method
F
ATAC
RNA
RMSE
Model
SCIGMA.No.Contrastive.Loss
SCIGMA.Regular.Contrastive.Loss
SCIGMA.No.Spatial.Graph
SCIGMA.No.Feature.Graph
SCIGMA.GCN
SCIGMA.No.Uncertainty
SCIGMA

**Supplementary Figure 2. Spatial clusters identified by SCIGMA in the spatial ATAC-seq mouse brain dataset.** This scatterplot displays the individual spatial clusters separately. The identified localization of the 17 spatial clusters by SCIGMA satisfactorily captured the brain anatomic structure, including cortex layers 2-6 (clusters 1, 2, 4, 8), CCG (cluster 13), CP (cluster 5, 16), ACB (cluster 3), ACO (cluster 13), VL (cluster 14), and LPO (cluster 17), aligning closely with annotations from the Allen Reference Atlas.

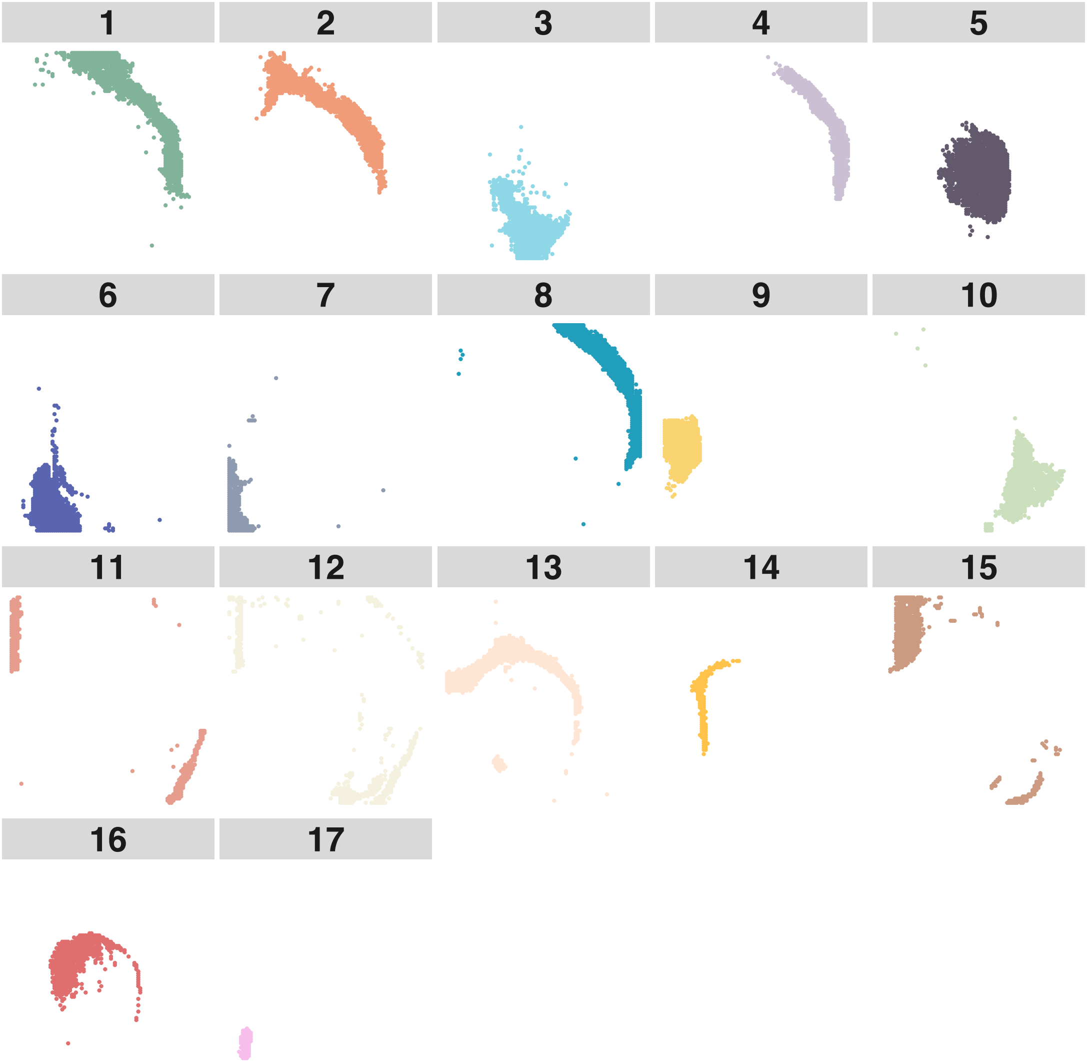

**Supplementary Figure 3. Heatmap of expression pattern of the domain specific DE genes by SCIGMA.** Due to the space issue, we only display the top 10 selected DE genes. The yellow color represents a higher expression while the purple color represents a lower expression.

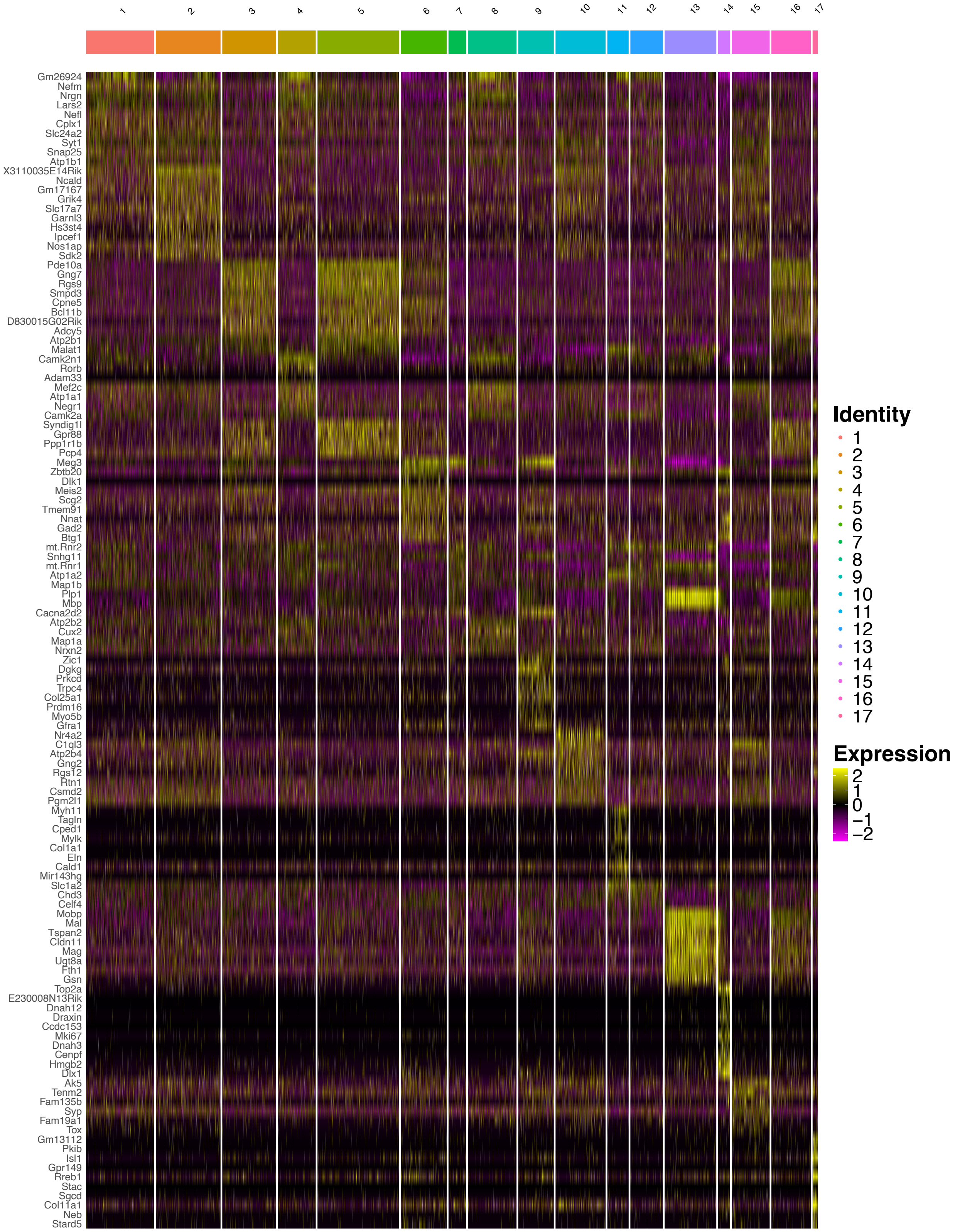

**Supplementary Figure 4. Scatter plots displaying the spatial distribution of important marker genes identified in the DE analysis in the spatial ATAC-seq mouse brain dataset.** For each gene, the left panel shows its spatial distribution in the ATAC-seq data, and the right panel shows the corresponding in situ hybridization (ISH) image from the Allen Mouse Brain Atlas, where available. The observed spatial enrichment of these DE genes is consistent with their known anatomical expression patterns, supporting the biological relevance of the identified markers. ISH reference data for Mobp were not available from Allen Mouse Brain Atlas.

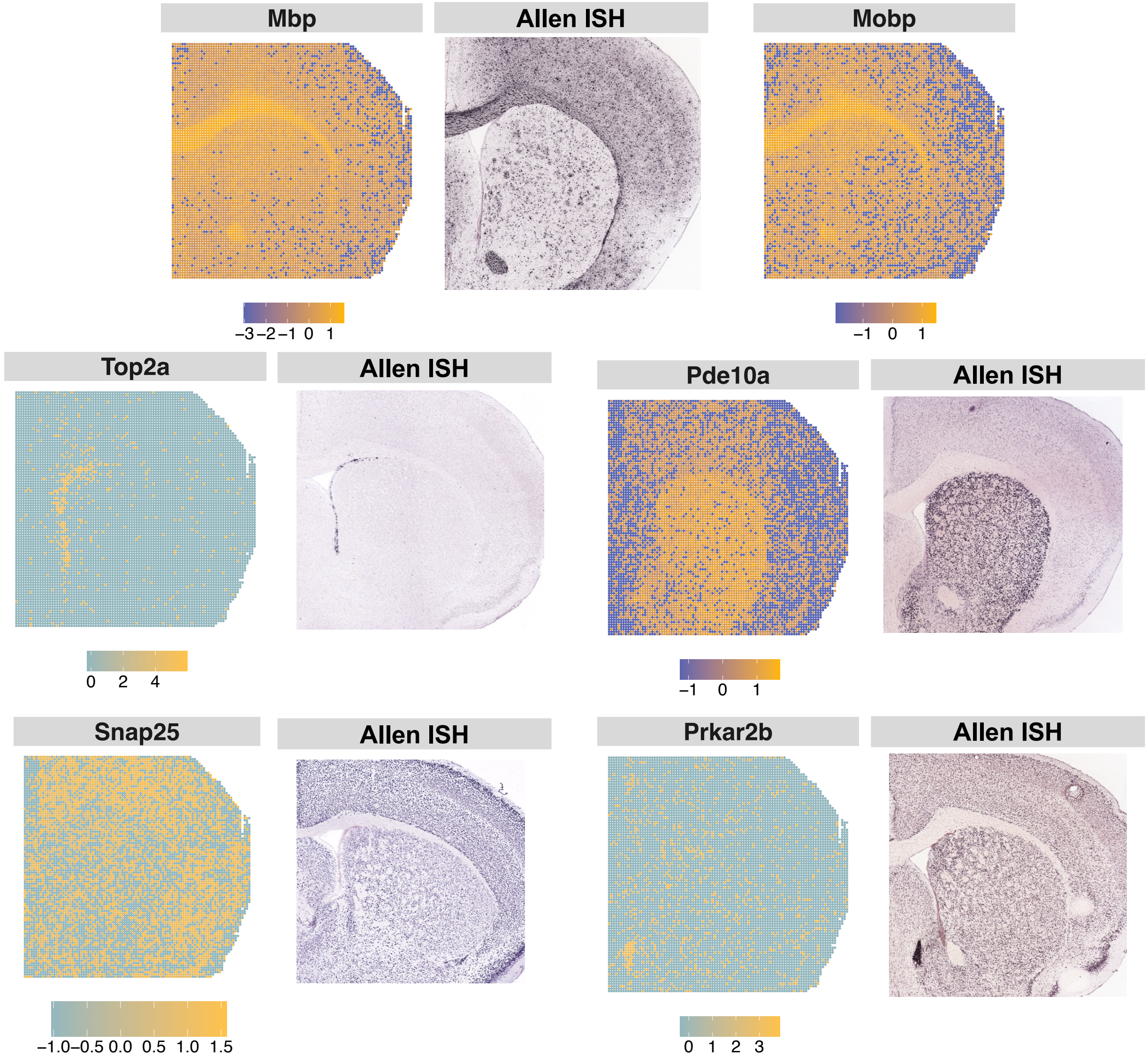

**Supplementary Figure 5. Differentially expressed peak analysis for spatial ATAC-seq mouse brain dataset.** Heatmap of the differentially expressed peaks for each SCIGMA cluster. Red color represents a higher chromatin accessibility while blue color represents a lower value.

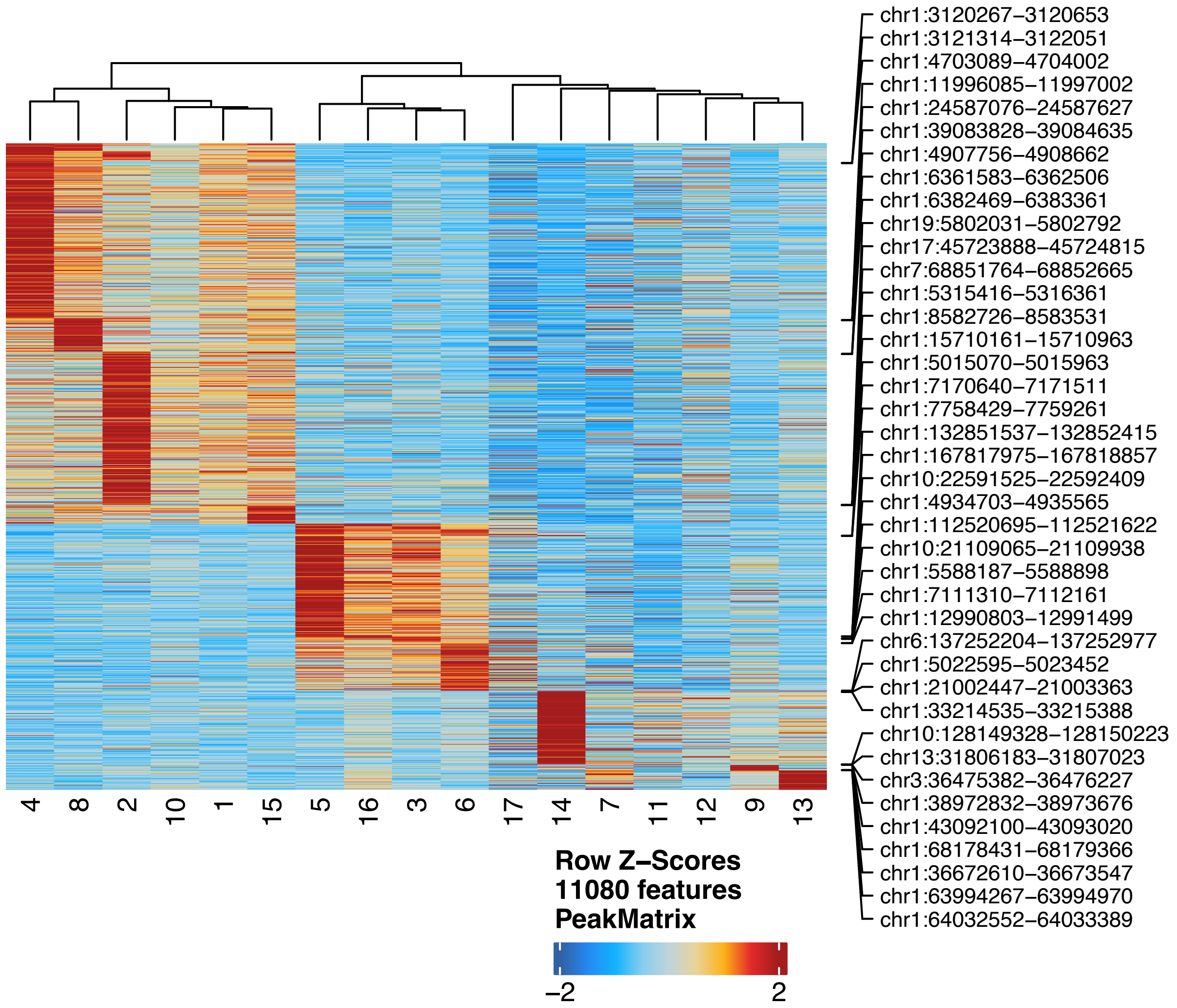

**Supplementary Figure 6. Peak-to-gene link heatmap for spatial ATAC-seq mouse brain dataset.** Peak-to-gene link heatmap of the differentially expressed peaks with the differentially expressed genes for each SCIGMA cluster. This heatmap plots side by side the linked ATAC and Gene regions. The highlighted boxes along the top x-axis represent the regions captured by SCIGMA – the purple boxes represent the cortex layers (cluster 2, 4, 8), the green boxes represent the striatum (clusters 5, 16), and the light blue boxes represent the corpus callosum (cluster 13). The red boxes on the heatmap show the linkage corresponding to the highlighted regions.

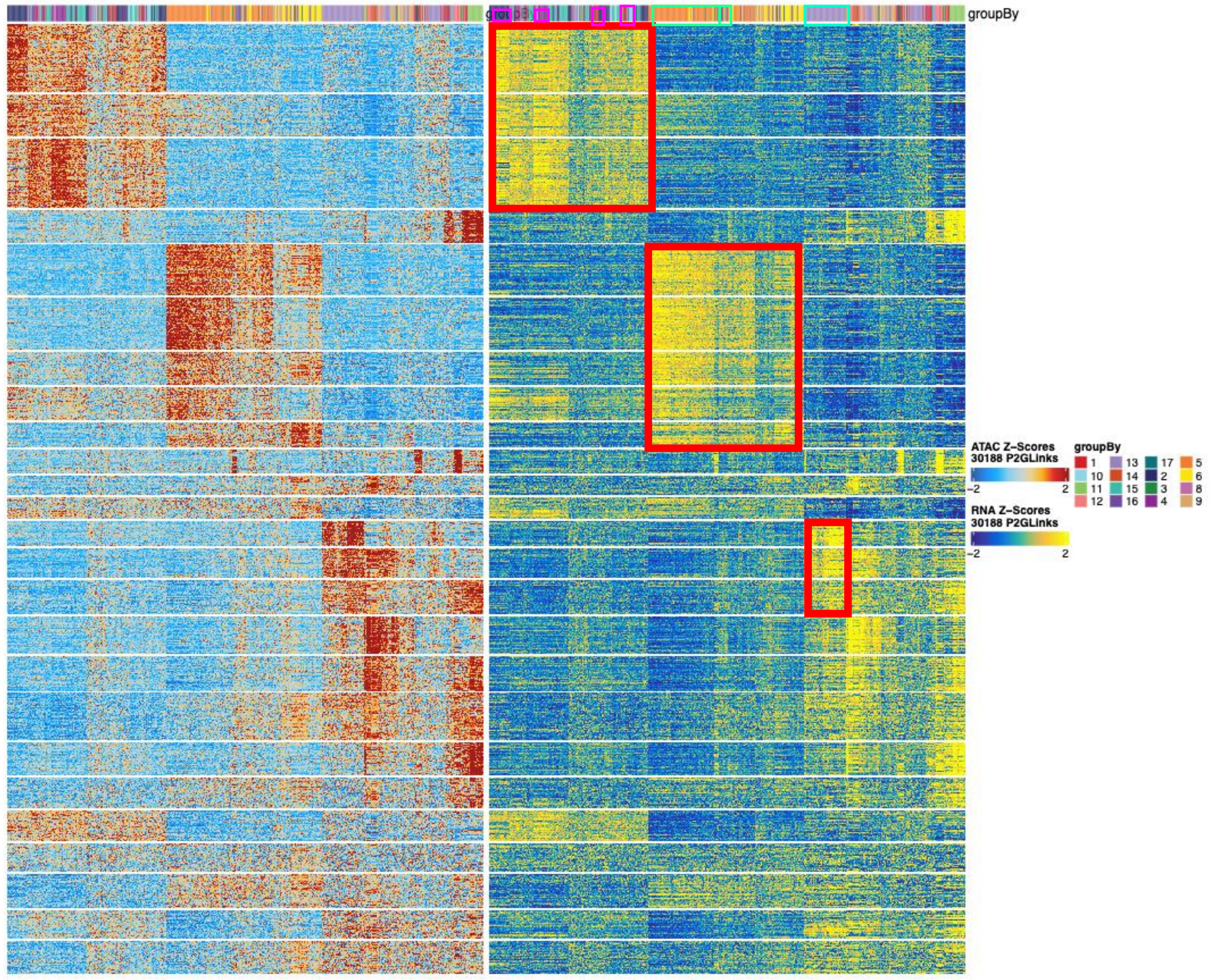

**Supplementary Figure 7. GSEA results on each spatial cluster detected by SCIGMA in the spatial ATAC-seq mouse brain dataset.** Here, top 10 (if available) enriched gene sets were displayed.

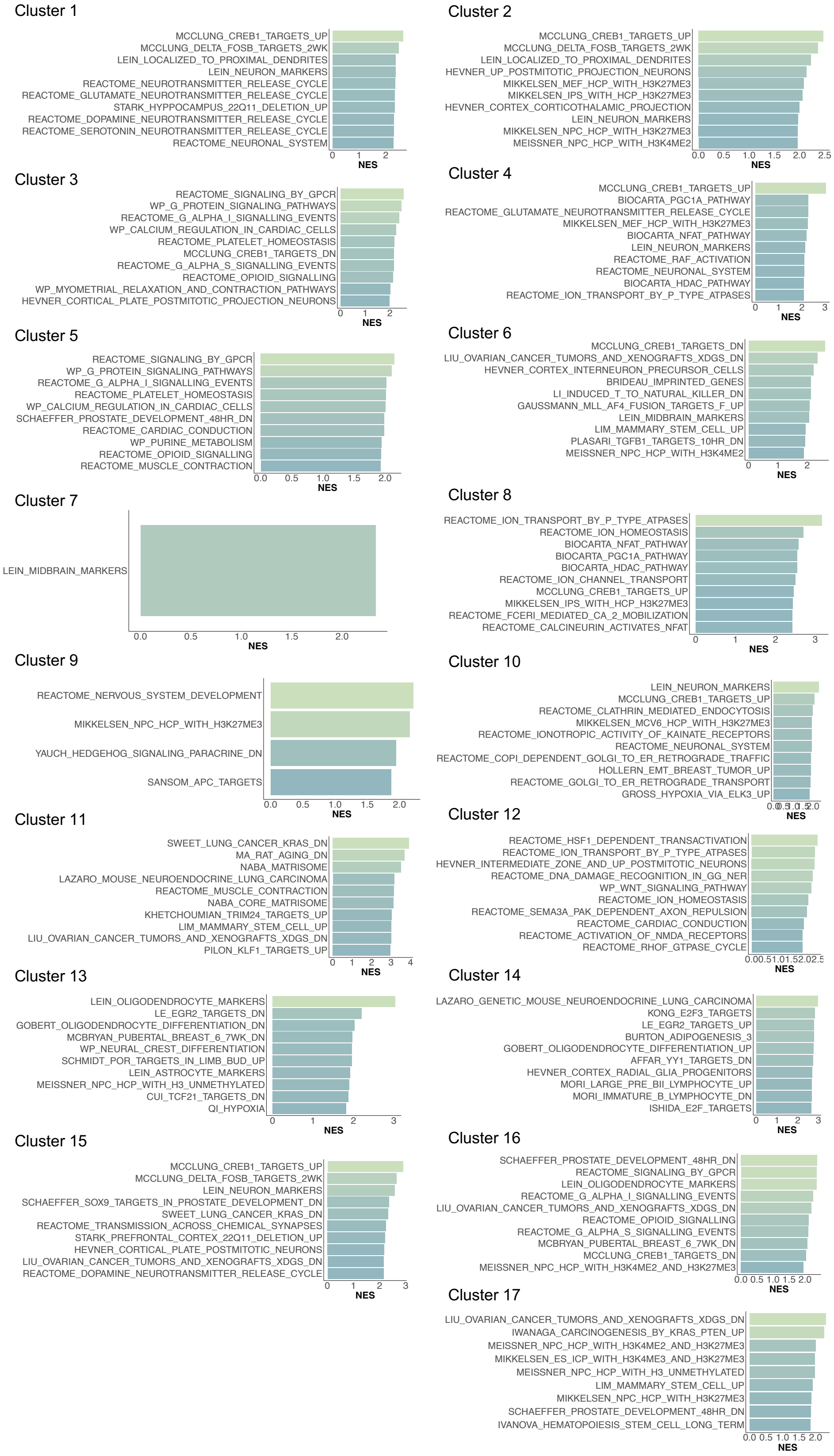

**Supplementary Figure 8. Violin plots display the distribution of modality-specific weights in the spatial ATAC-seq mouse brain dataset.** Here the weights represent the Attention weights from SCIGMA's attention layer, with the higher weight representing that SCIGMA learned to attend to a specific modality more for a given cluster.

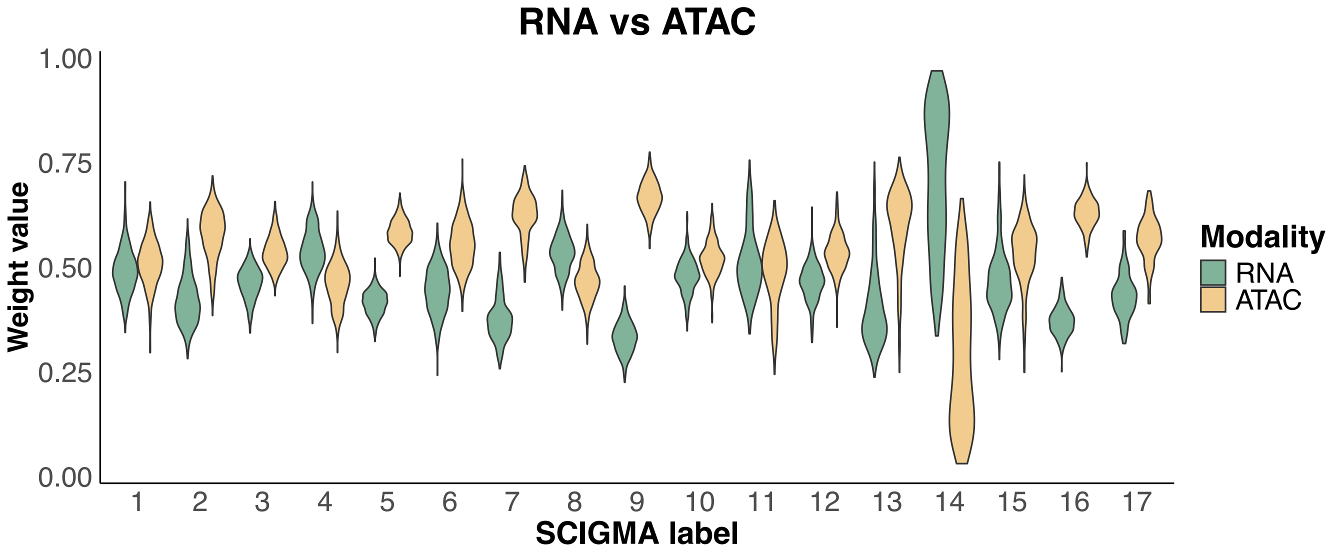

**Supplementary Figure 9. Spatial distribution of key marker features (e.g., peaks, or genes) in SCIGMA's reconstructed data versus original data for the spatial ATAC-seq mouse brain dataset.**

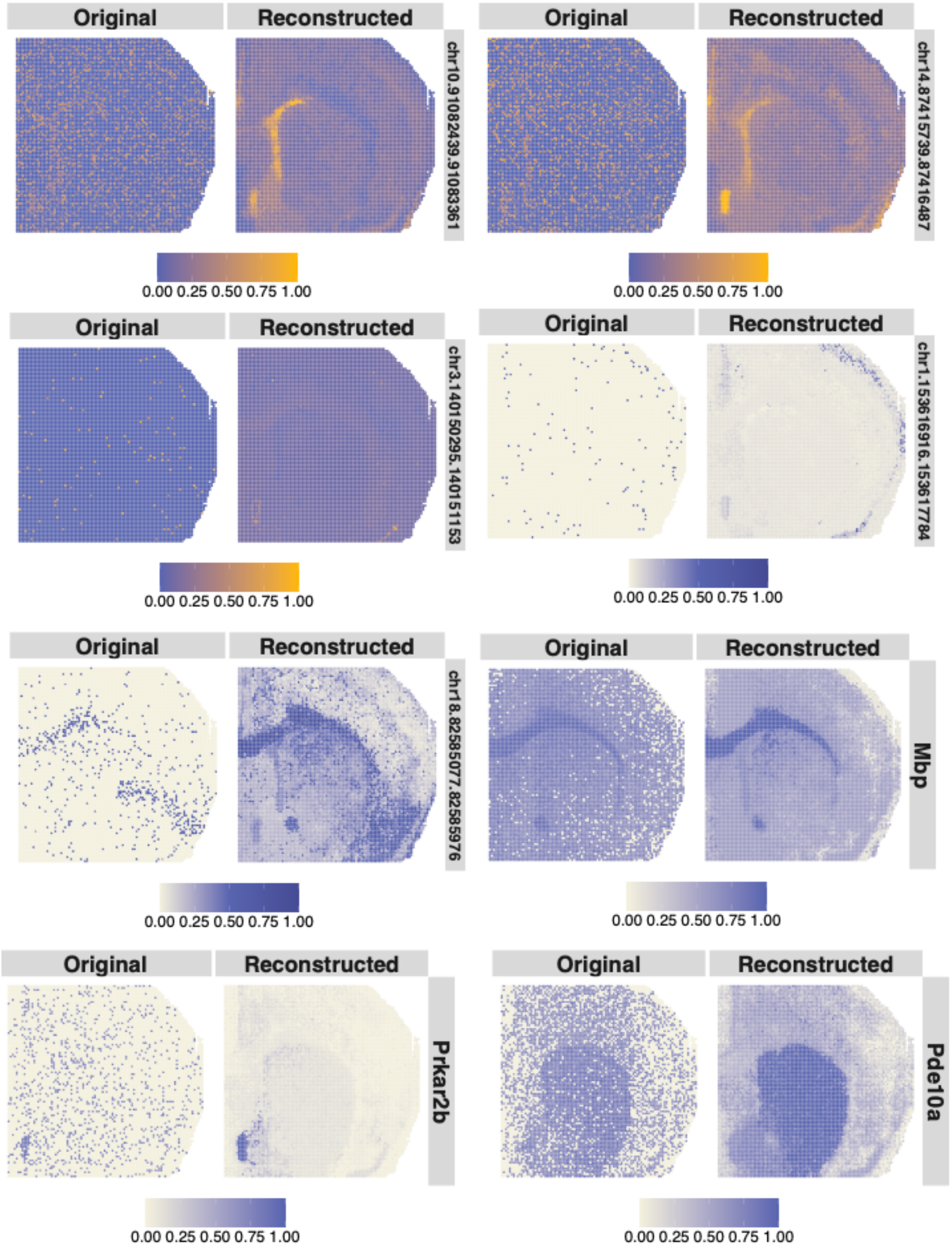

**Supplementary Figure 10. Interpretations on uncertainty estimates for the spatial ATAC-seq mouse brain dataset. (A)** Spatial distribution of uncertainty estimates generated by SCIGMA. **(B)** Boxplot of the total scaled expression absolute difference between RNA and ATAC modalities, comparing uncertain and normal spatial locations. The y-axis represents the absolute difference. **(C)** Boxplot of the RMSE between the reconstructed and original feature matrices for RNA and ATAC modalities, comparing uncertain and normal spatial locations. The y-axis represents the RMSE. **(D)** Boxplot of the Jaccard index, measuring the similarity between RNA-specific and joint embeddings, as well as ATAC-specific and joint embeddings, respectively. Comparisons are made between uncertain and normal spatial locations. The y-axis represents the Jaccard index. Here, the stars ("*") indicate the significance of a one-sided Wilcoxon rank-sum test between certain and uncertain regions, where the uncertain regions are defined as the top 1%, 5%, 10%, and 20% of uncertainty values, ranked from largest to smallest.

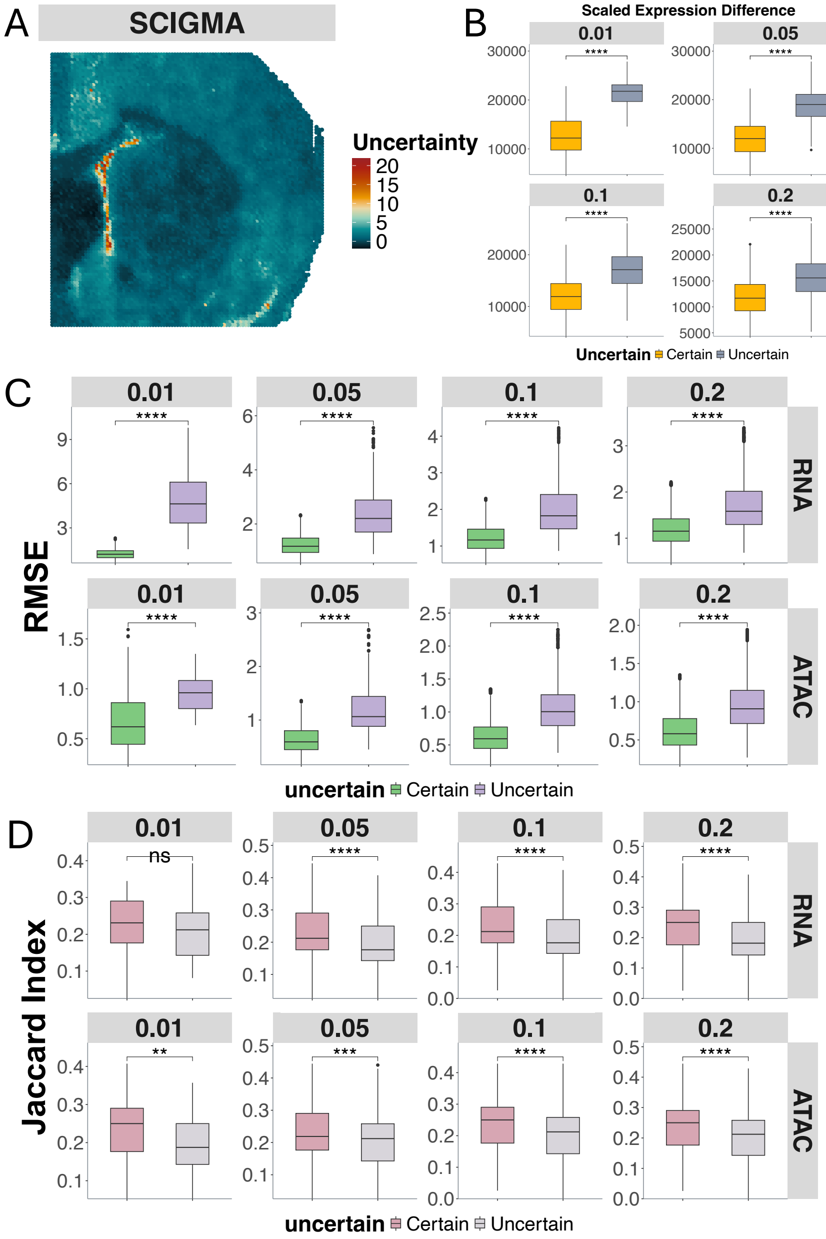

**Supplementary Figure 11. Spatial clusters identified by SCIGMA in SPOTS mouse spleen dataset.** This scatterplot displays the individual spatial clusters separately. The identified localization of the 6 spatial clusters by SCIGMA satisfactorily captured the spleen anatomic structure, including T cell region (cluster 5), B cell (cluster 4), red pulp (cluster 2), MZ region (cluster 3), inner border of MZ (cluster 1), and epithelial cells (cluster 6).

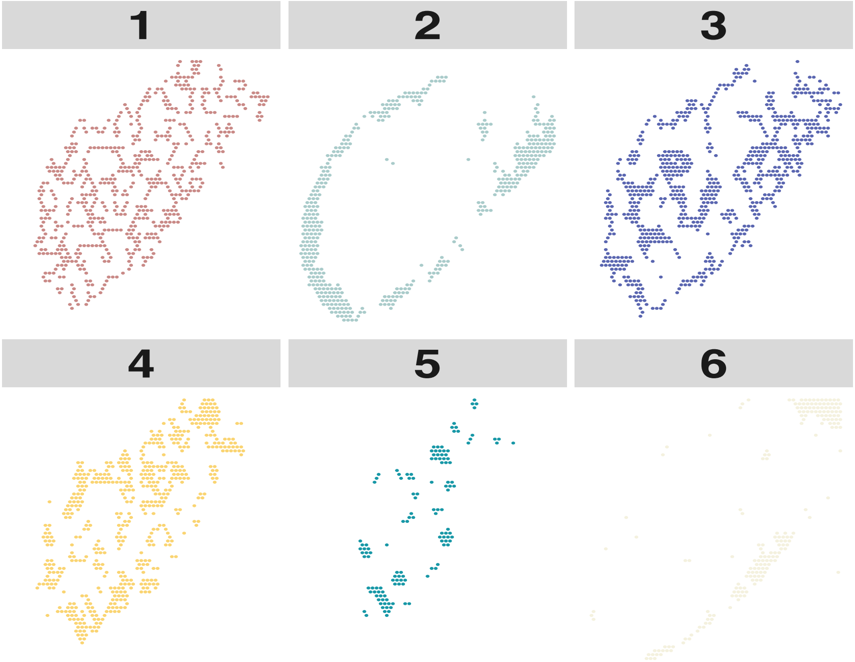

**Supplementary Figure 12. Differentially expressed genes analysis for SPOTS mouse spleen dataset.** Due to the space issue, we only display the top 10 selected DE genes. The red color represents a higher expression while the blue color represents a lower expression.

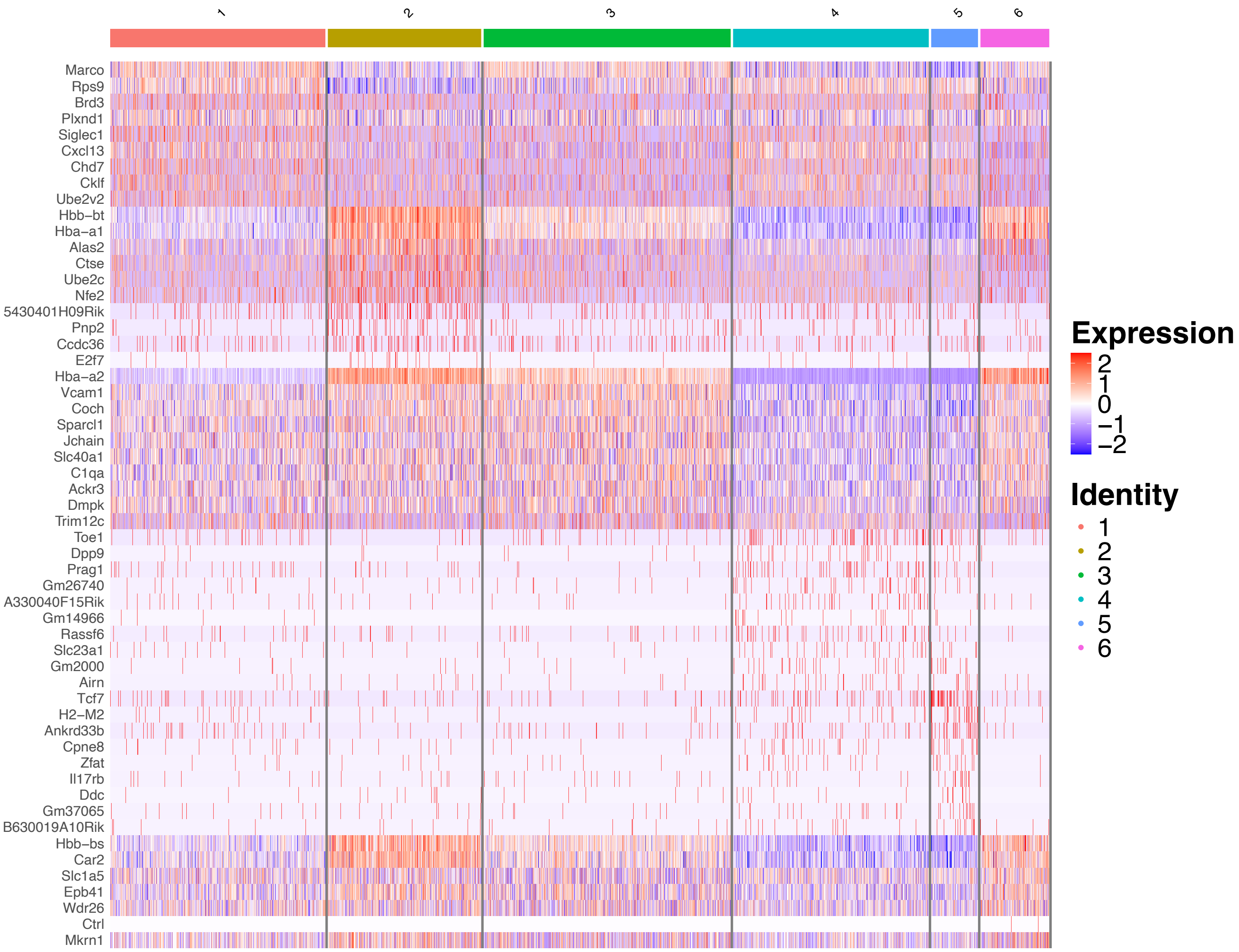

**Supplementary Figure 13. (A) Scatter plots display the spatial distribution of important marker genes identified in the DE analysis in the SPOTS mouse spleen dataset. (B) Scatter plots display the spatial distribution of important marker proteins identified in the Differential protein analysis in the SPOTS mouse spleen dataset.**

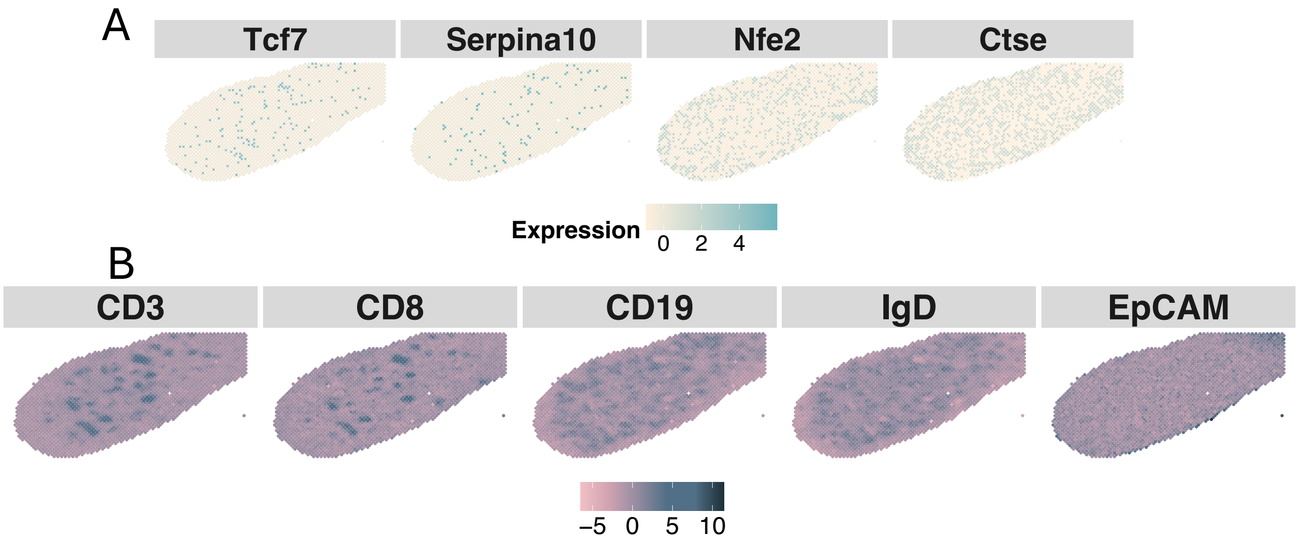

**Supplementary Figure 14. Differentially expressed protein analysis for SPOTS mouse spleen dataset.** Due to the space issue, we only display the top 10 selected DE ADTs. Red color represents a higher expression while blue color represents a lower expression.

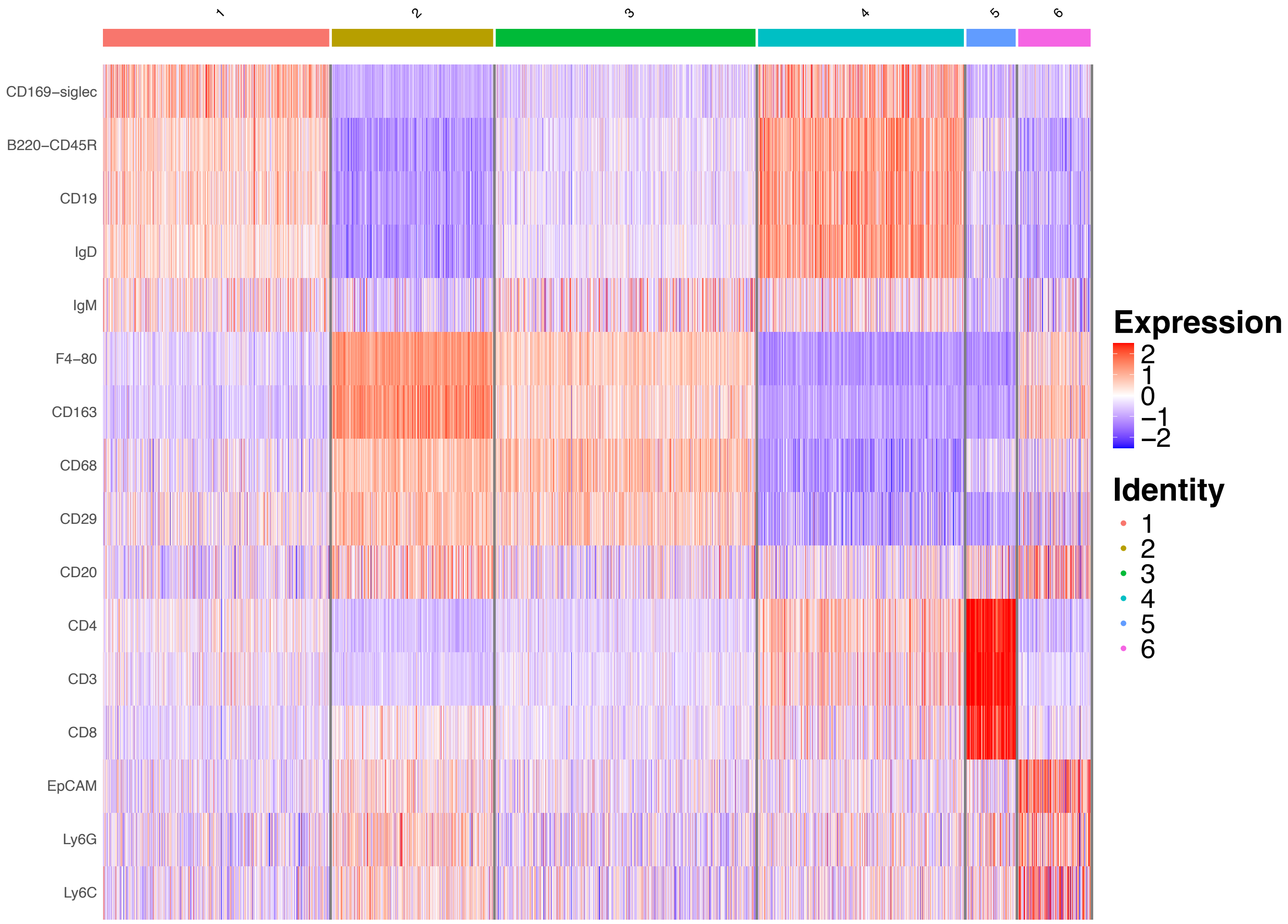

**Supplementary Figure 15. GSEA results on each spatial cluster detected by SCIGMA in the SPOTS mouse spleen dataset.** Here, top 10 (if available) enriched gene sets were displayed.

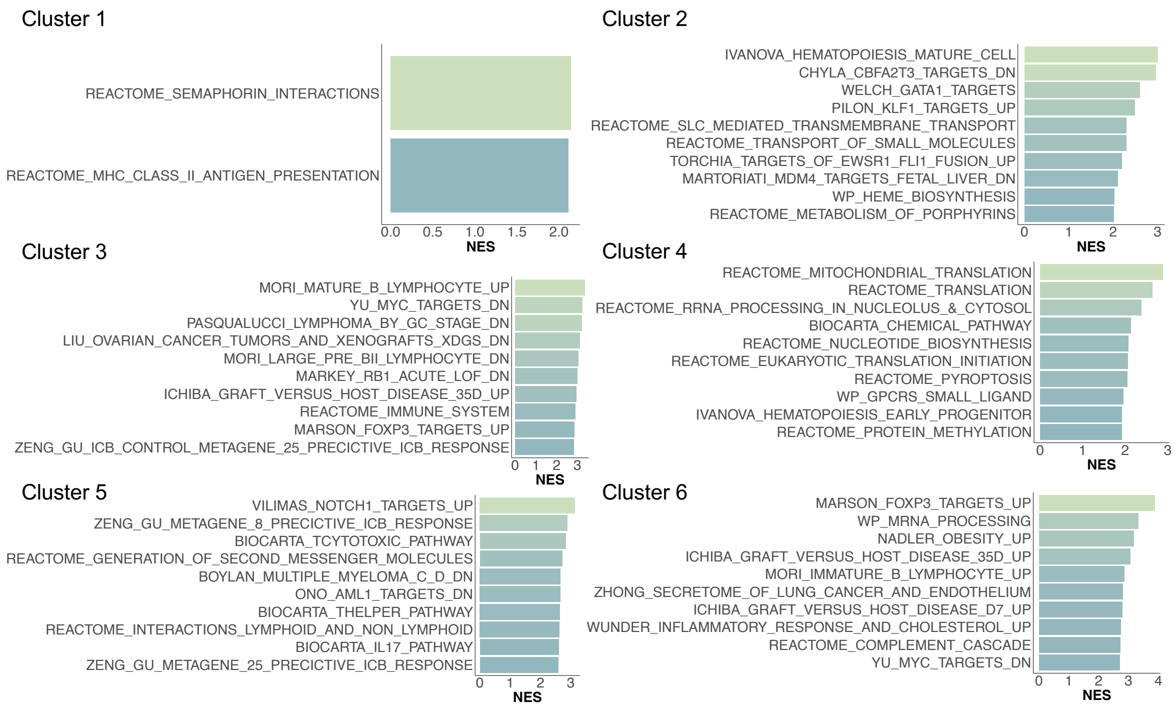

**Supplementary Figure 16. Violin plots display the distribution of modality-specific weights in the SPOTS mouse spleen dataset.** Here the weights represent the Attention weights from SCIGMA's attention layer, with the higher weight representing that SCIGMA learned to attend to a specific modality more for a given cluster.

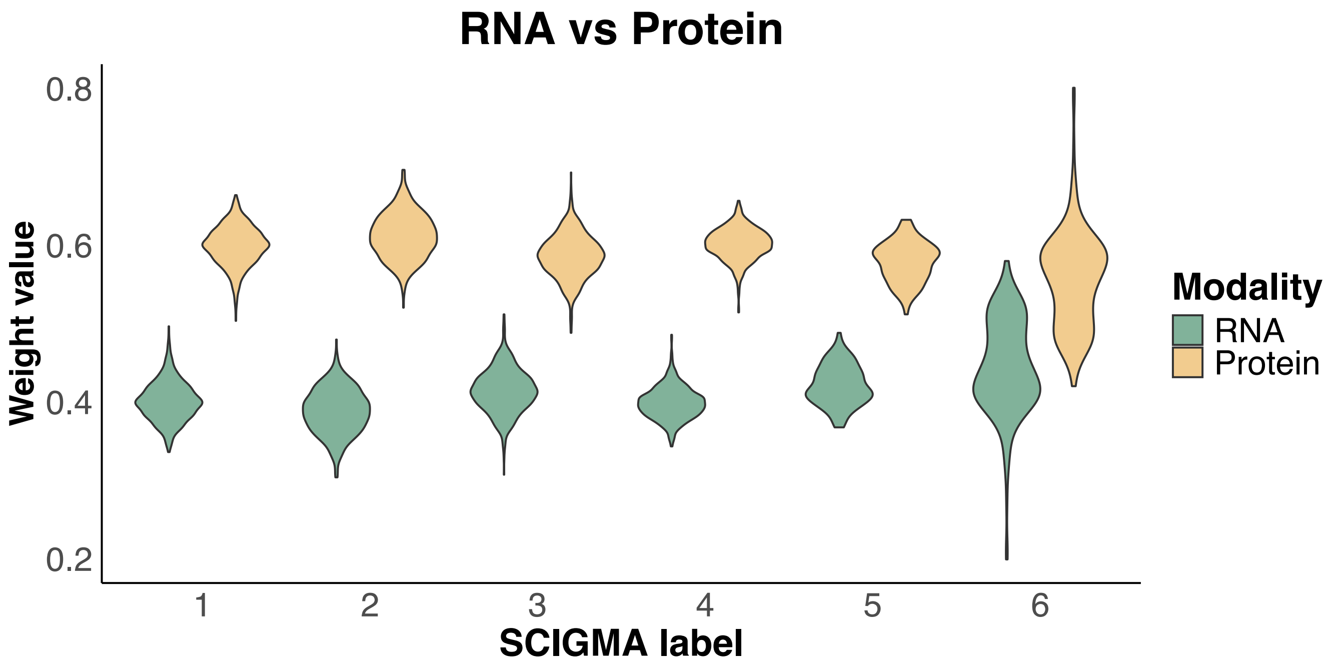

**Supplementary Figure 17. (A) Spatial distribution of key marker proteins in SCIGMA's reconstructed data versus original data for the SPOTS mouse spleen dataset. (B) Spatial distribution of key marker genes in SCIGMA's reconstructed data versus original data for the SPOTS mouse spleen dataset.**

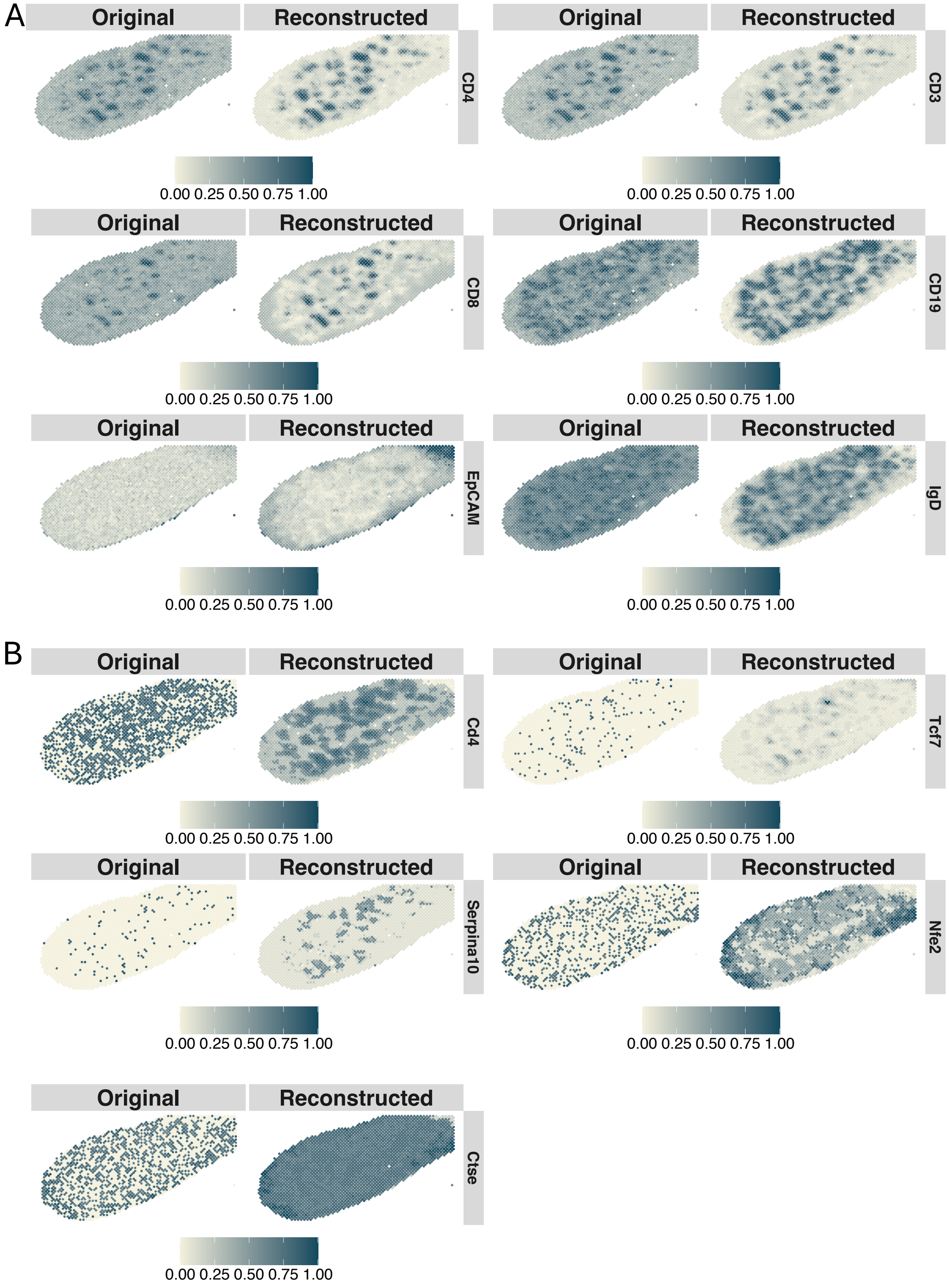

**Supplementary Figure 18. Interpretations of uncertainty estimates for the SPOTS mouse spleen dataset. (A)** Spatial distribution of uncertainty estimates generated by SCIGMA. **(B)** Boxplot of the total scaled expression absolute difference between RNA and protein modalities, comparing uncertain and normal spatial locations. The y-axis represents the absolute difference. **(C)** Boxplot of the RMSE between the reconstructed and original feature matrices for RNA and protein modalities, comparing uncertain and normal spatial locations. The y-axis represents the RMSE. **(D)** Boxplot of the Jaccard index, measuring the similarity between RNA-specific and joint embeddings, as well as protein-specific and joint embeddings, respectively. Comparisons are made between uncertain and normal spatial locations. The y-axis represents the Jaccard index. Here, the stars ("*") indicate the significance of a one-sided Wilcoxon rank-sum test between certain and uncertain regions, where the uncertain regions are defined as the top 1%, 5%, 10%, and 20% of uncertainty values, ranked from largest to smallest.

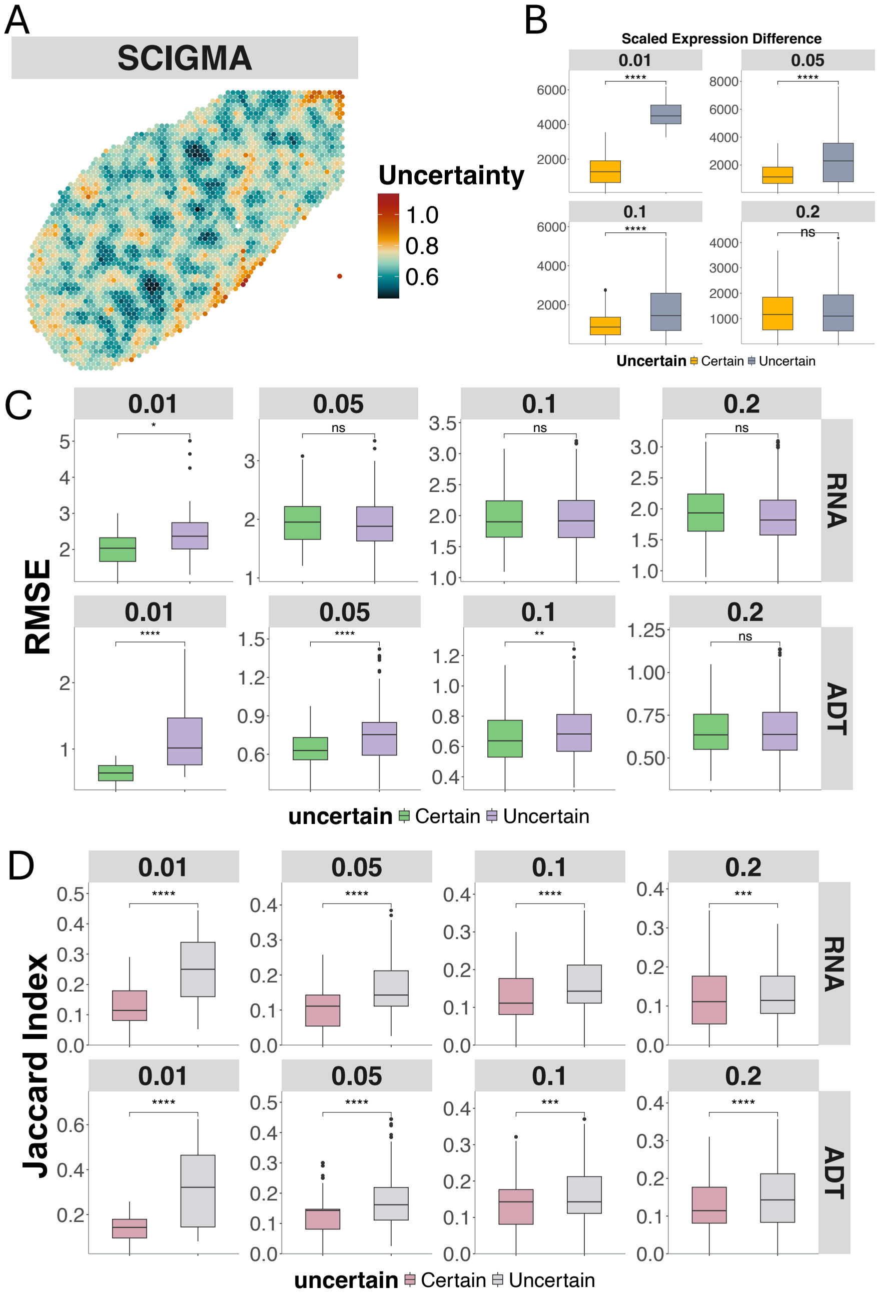

**Supplementary Figure 19. Spatial clusters identified by SCIGMA in 10xXenium human ovarian adenocarcinoma dataset.** This scatterplot displays the individual spatial clusters separately. The identified localization of the 20 spatial clusters by SCIGMA satisfactorily captured the human ovarian adenocarcinoma anatomic structure, including tumor subregions (cluster 2, 10, 11, 17) such as hypoxia-driven tumors (cluster 2) and aggressive invasive fronts (cluster 17); endothelial region (cluster 9); immune regions (cluster 7); and granulosa cells (cluster 8, 15, 20).

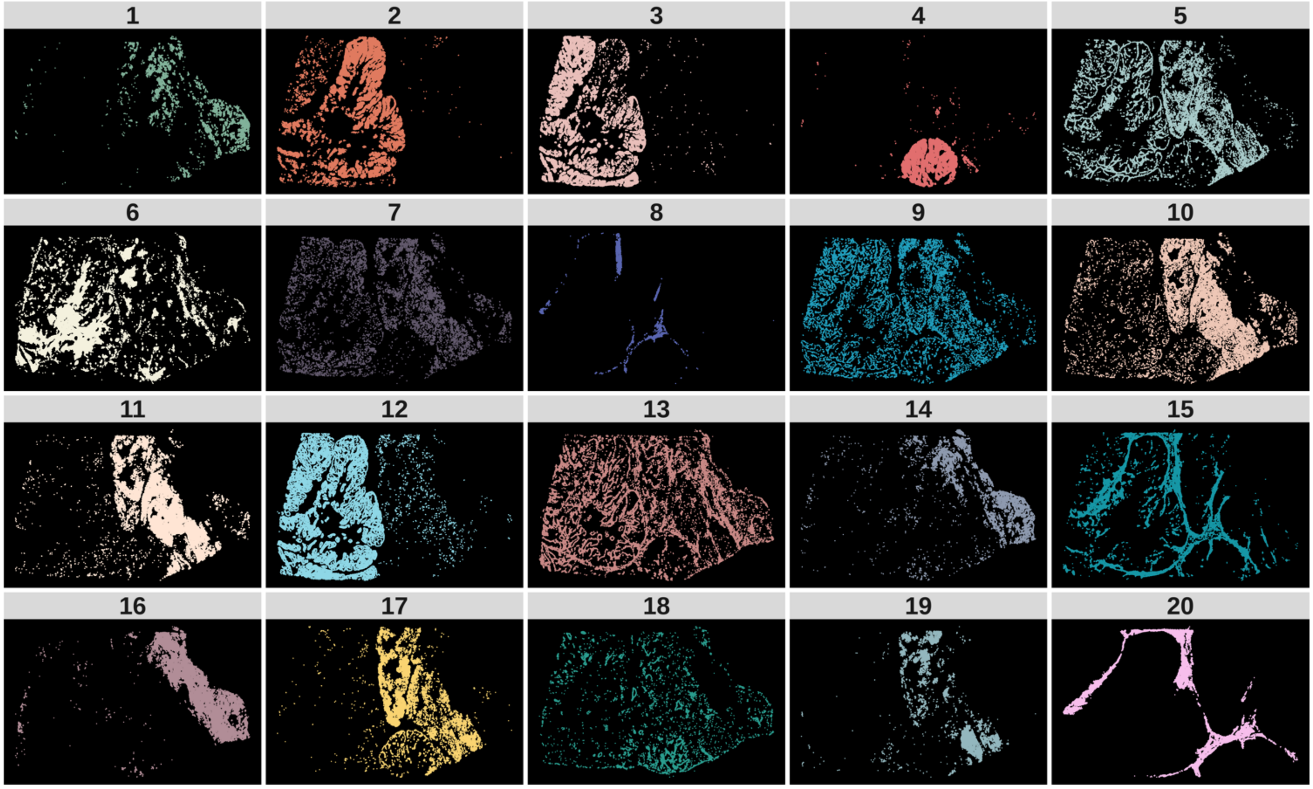

**Supplementary Figure 20. Differentially expressed genes analysis for 10xXeniumPrime human ovarian adenocarcinoma dataset.** Due to the space issue, we only display the top 10 selected DE genes. The red color represents a higher expression while the blue color represents a lower expression.

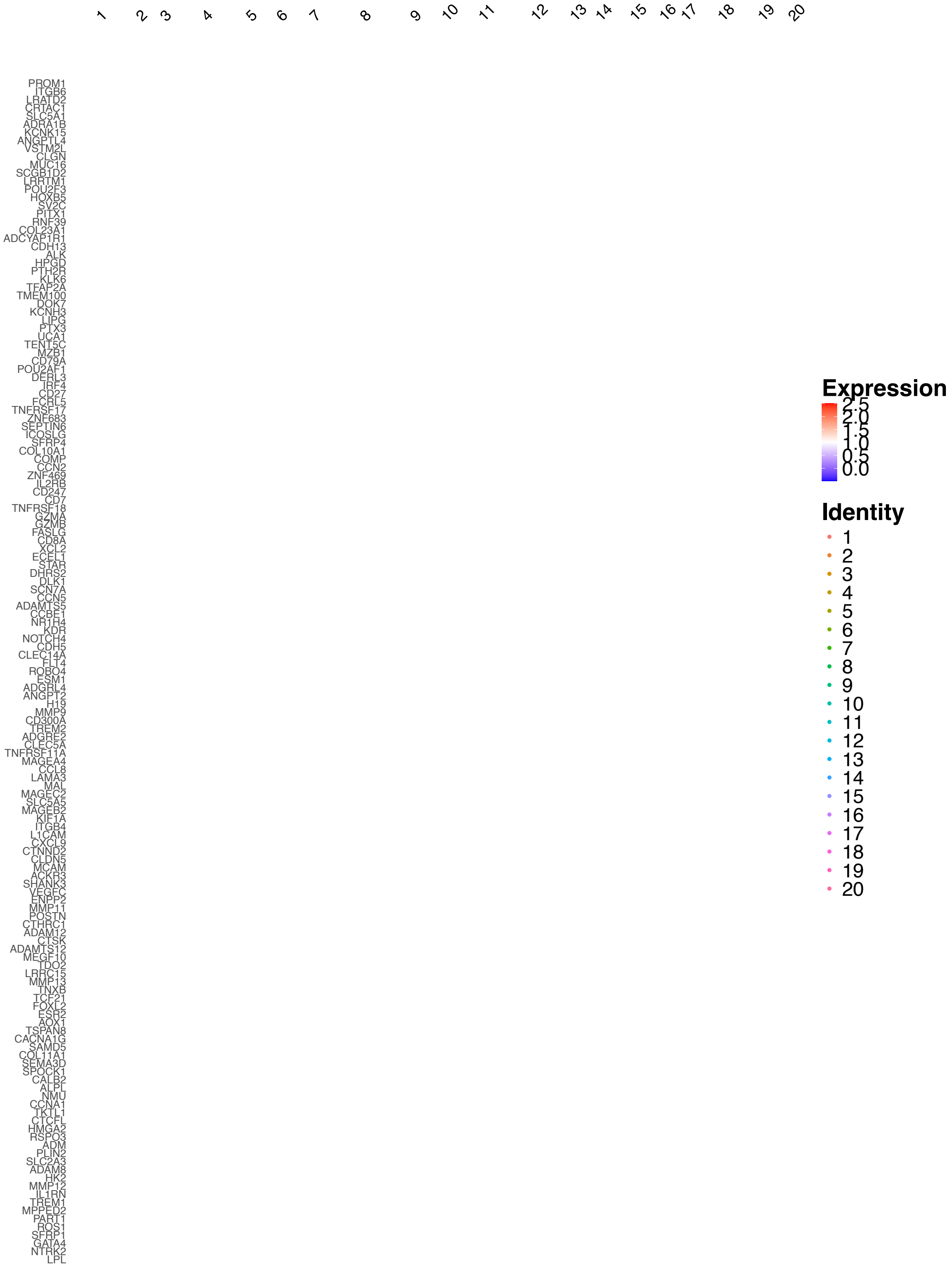

**Supplementary Figure 21. Scatter plots display the spatial distribution of important marker genes identified in the DE analysis in the 10xXeniumPrime human ovarian adenocarcinoma dataset.**

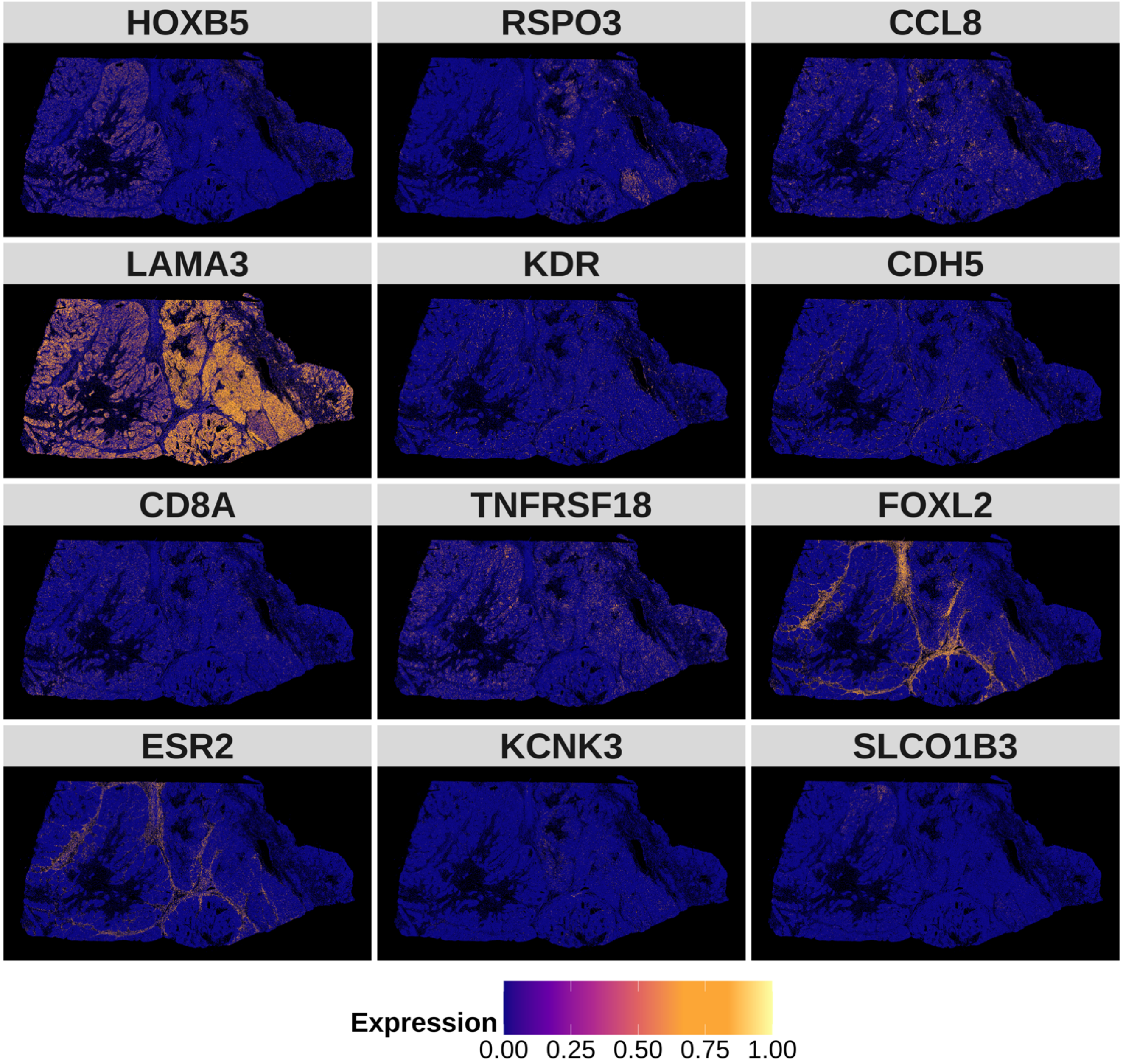

**Supplementary Figure 22. GSEA results for the 10xXeniumPrime human ovarian adenocarcinoma dataset.** Top 10 (if available) gene set enrichment results.

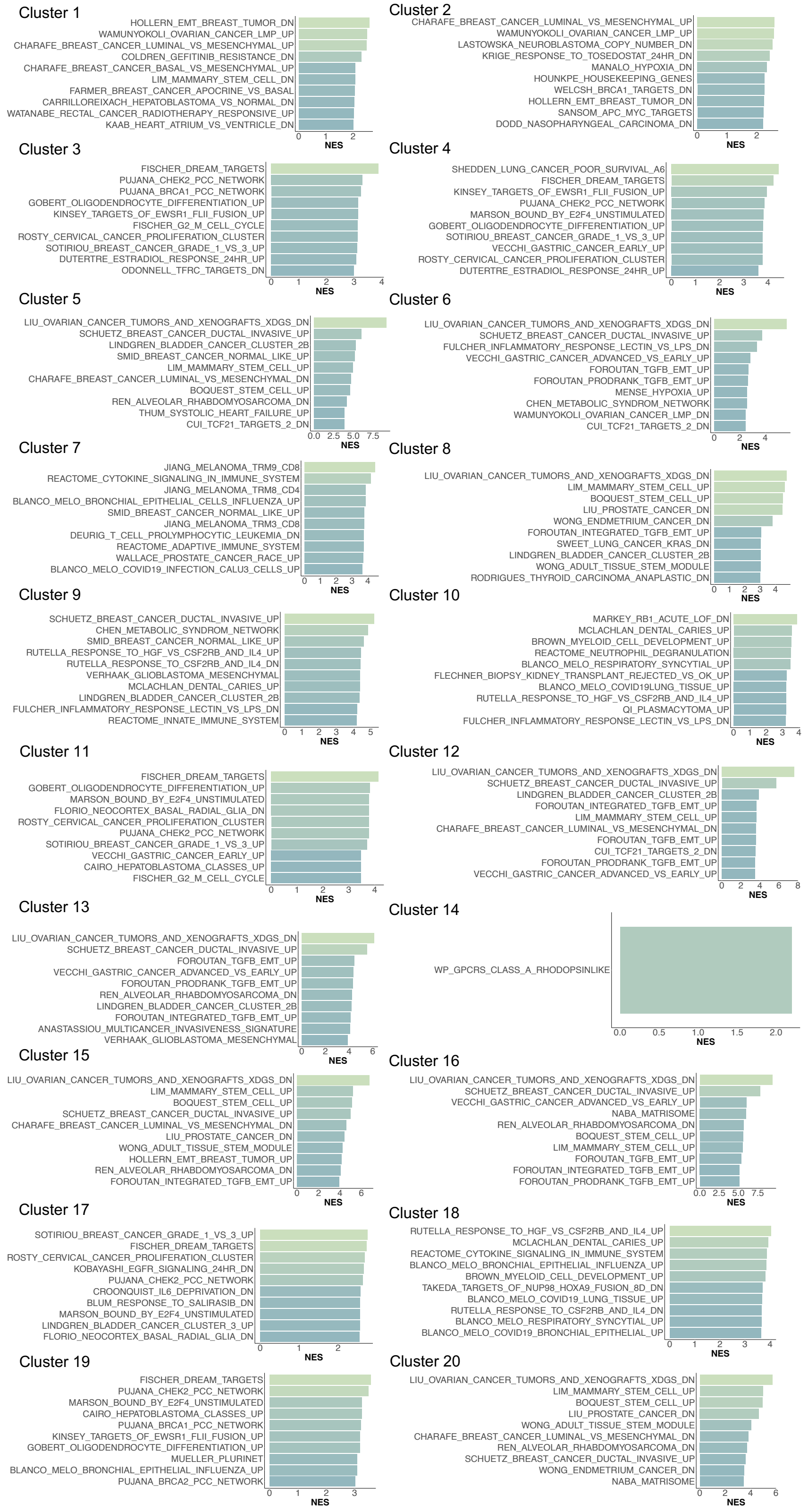

**Supplementary Figure 23. Modality specific weights for 10xXeniumPrime human ovarian adenocarcinoma dataset.** Here the weights represent the Attention weights from SCIGMA's attention layer, with the higher weight representing that SCIGMA learned to attend to a specific modality more for a given cluster.

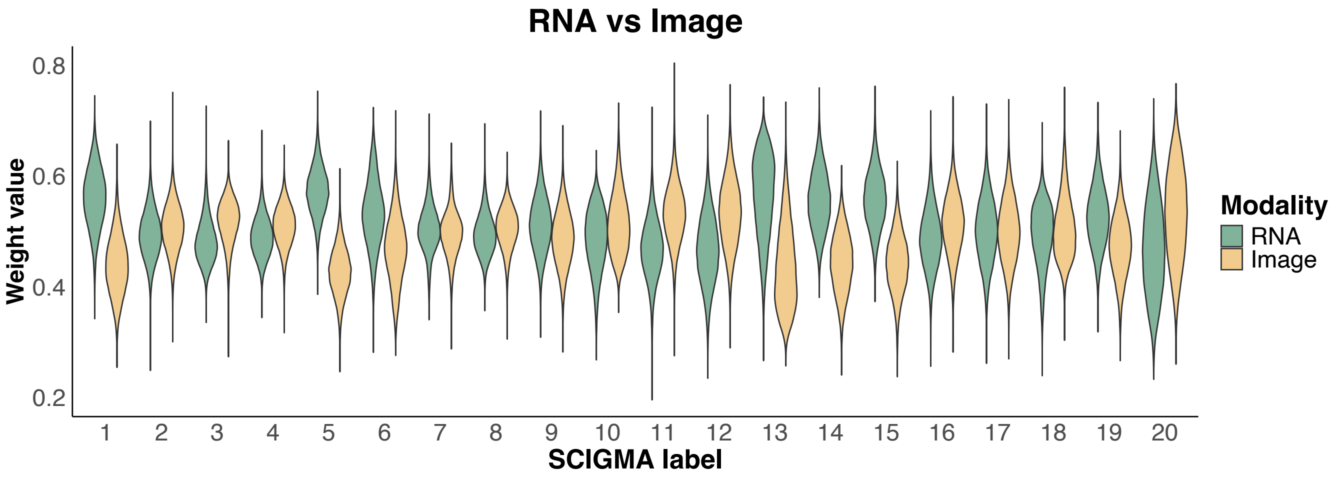

**Supplementary Figure 24. Spatial distribution of key marker genes in SCIGMA's reconstructed data versus original data for the 10xXeniumPrime human ovarian adenocarcinoma dataset.**

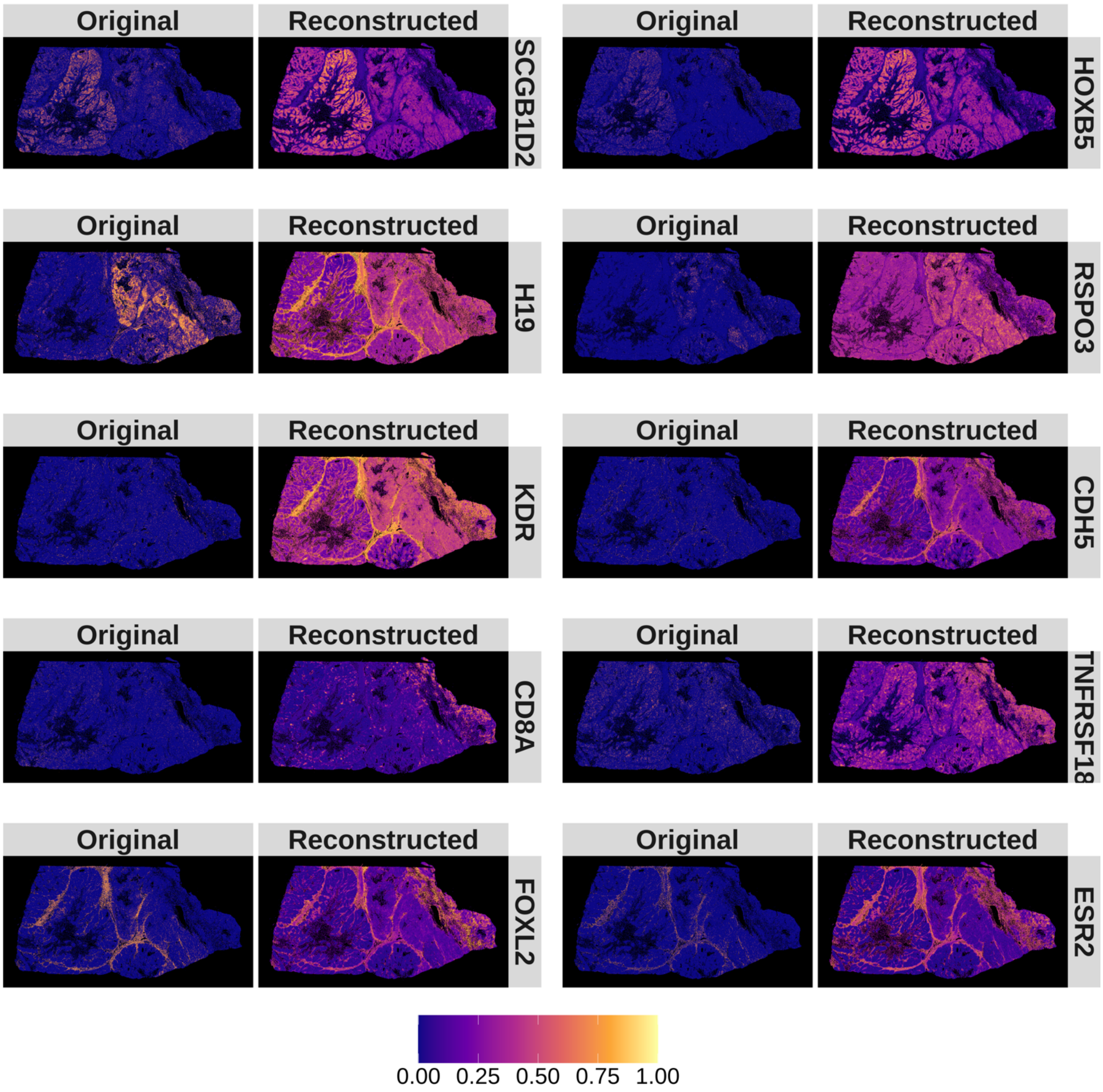

**Supplementary Figure 25. Interpretations of the uncertainty estimates for the 10xXeniumPrime human cervical cancer dataset. (A)** Spatial distribution of uncertainty estimates generated by SCIGMA. **(B)** Boxplot of the total scaled expression absolute difference between RNA and imaging modalities, comparing uncertain and normal spatial locations. The y-axis represents the absolute difference. **(C)** Boxplot of the RMSE between the reconstructed and original feature matrices for RNA and imaging modalities, comparing uncertain and normal spatial locations. The y-axis represents the RMSE. **(D)** Boxplot of the Jaccard index, measuring the similarity between RNA-specific and joint embeddings, as well as imaging-specific and joint embeddings, respectively. Comparisons are made between uncertain and normal spatial locations. The y-axis represents the Jaccard index. Here, the stars ("*") indicate the significance of a one-sided Wilcoxon rank-sum test between certain and uncertain regions, where the uncertain regions are defined as the top 1%, 5%, 10%, and 20% of uncertainty values, ranked from largest to smallest.

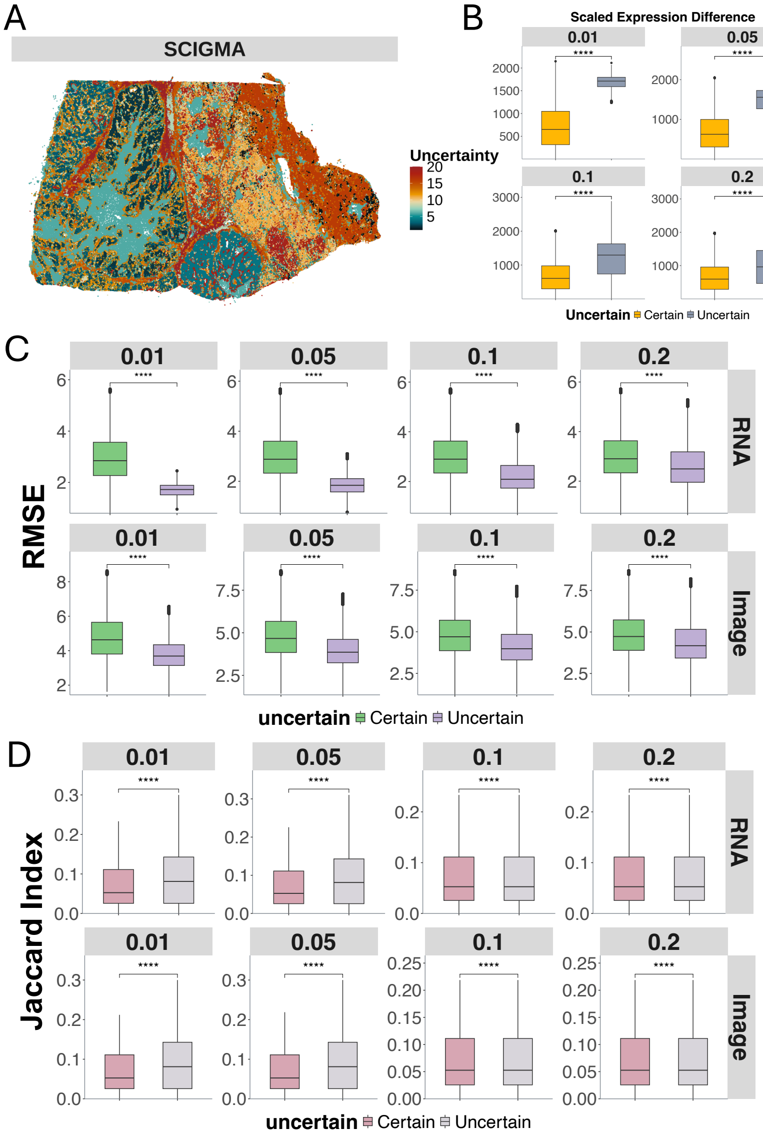

**Supplementary Figure 26. Spatial clusters identified by SCIGMA in 10xXeniumPrime human cervical cancer dataset.** This scatterplot displays the individual spatial clusters separately. The identified localization of the 20 spatial clusters by SCIGMA satisfactorily captured the tissue anatomic structure, including: tumor subregions (clusters 4, 6, 7, 9, 12, 13, 14, 15, 17) such as cancer associated fibroblasts (cluster 14) and epithelial tumors (cluster 13); immune regions (cluster 20); and epithelial cells (cluster 5).

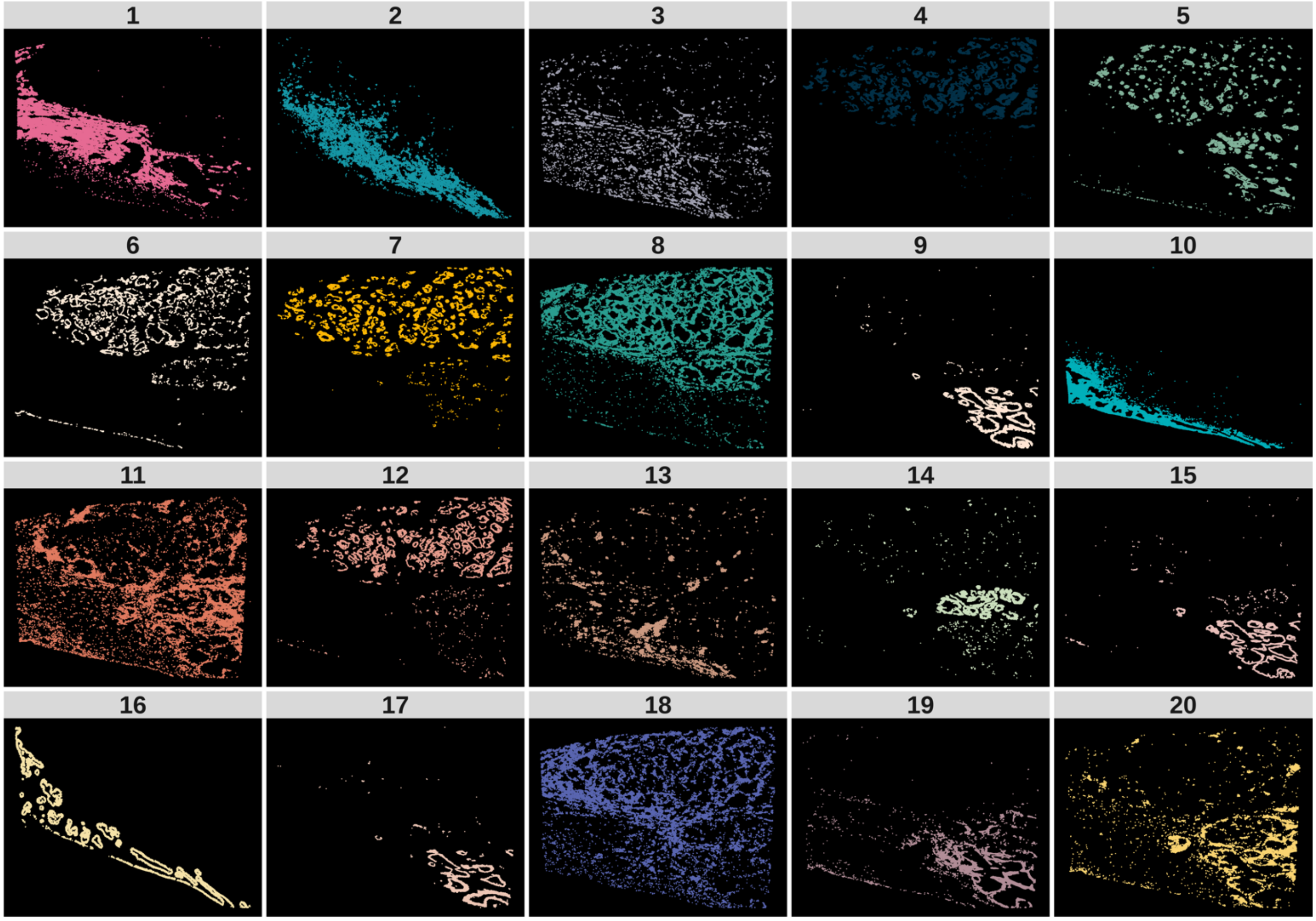

**Supplementary Figure 27. Differentially expressed genes analysis for 10xXeniumPrime human cervical cancer dataset.** Due to the space issue, we only display the top 10 selected DE genes. The red color represents a higher expression while the blue color represents a lower expression.

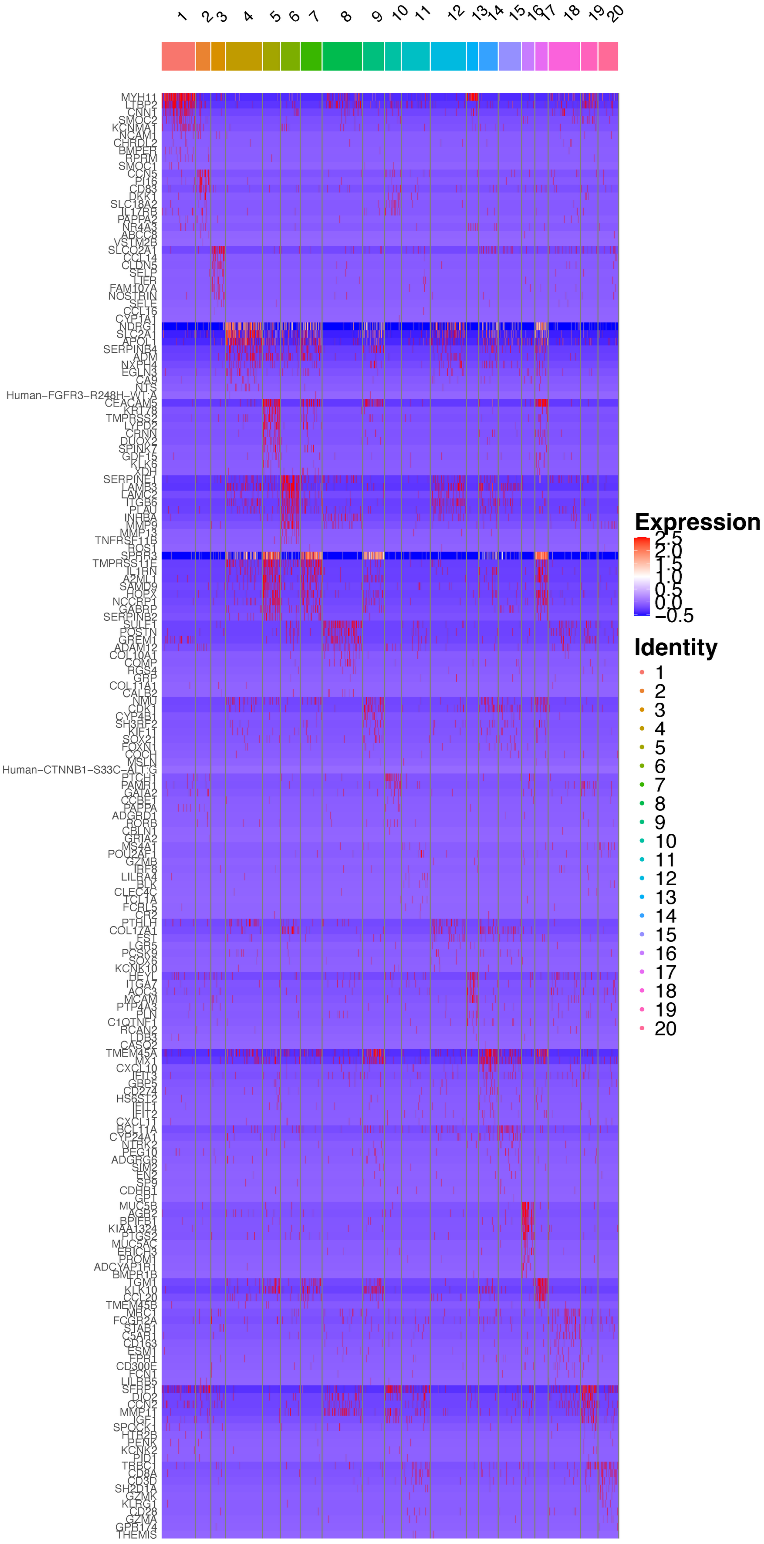

**Supplementary Figure 28.** **Scatter plots displaying the spatial distribution of important marker genes identified in the DE analysis in the 10xXeniumPrime human cervical cancer dataset.**

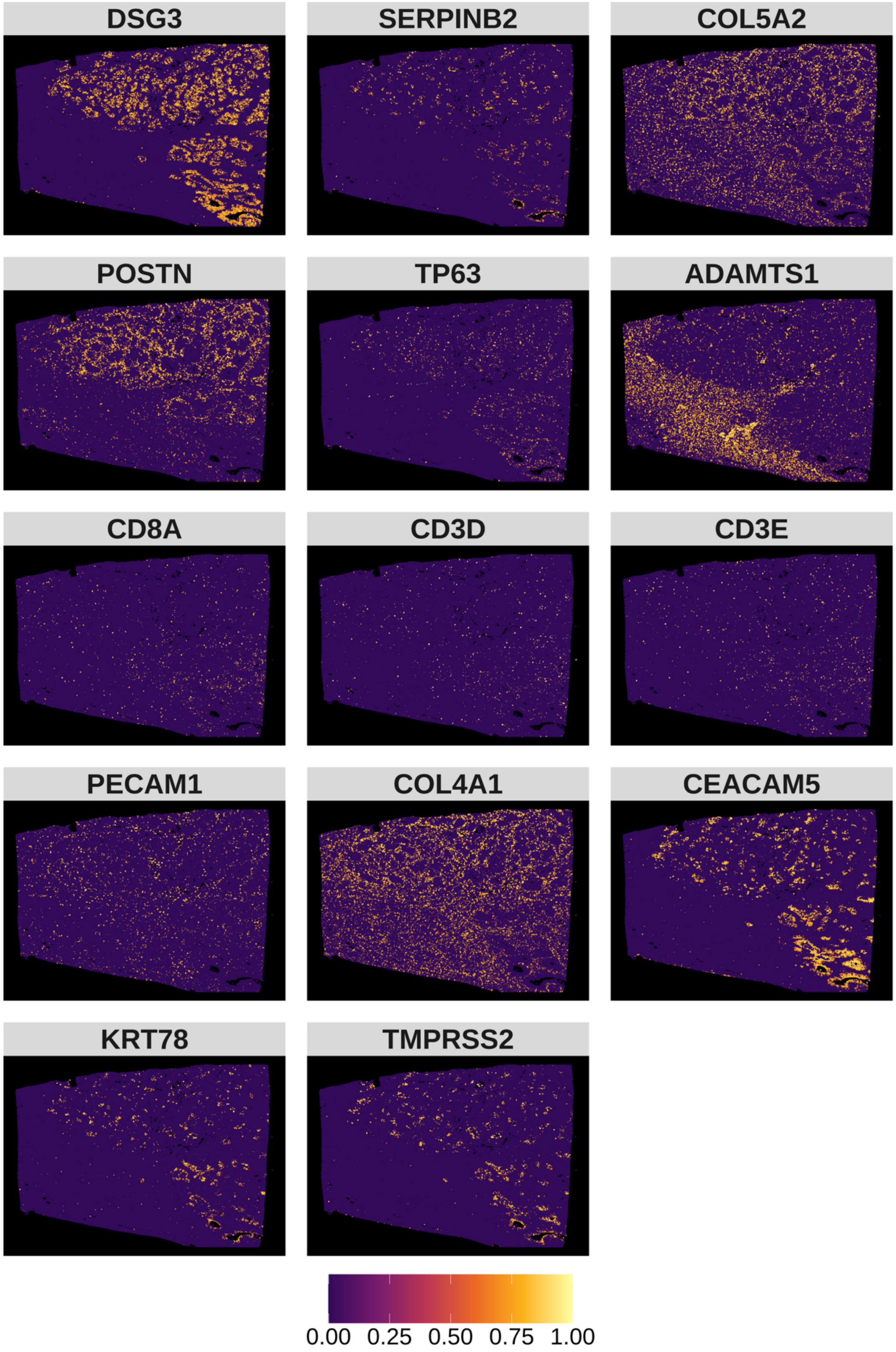

**Supplementary Figure 29. GSEA results for the 10xXeniumPrime human cervical cancer dataset.**
Top 10 (if available) gene set enrichment results.

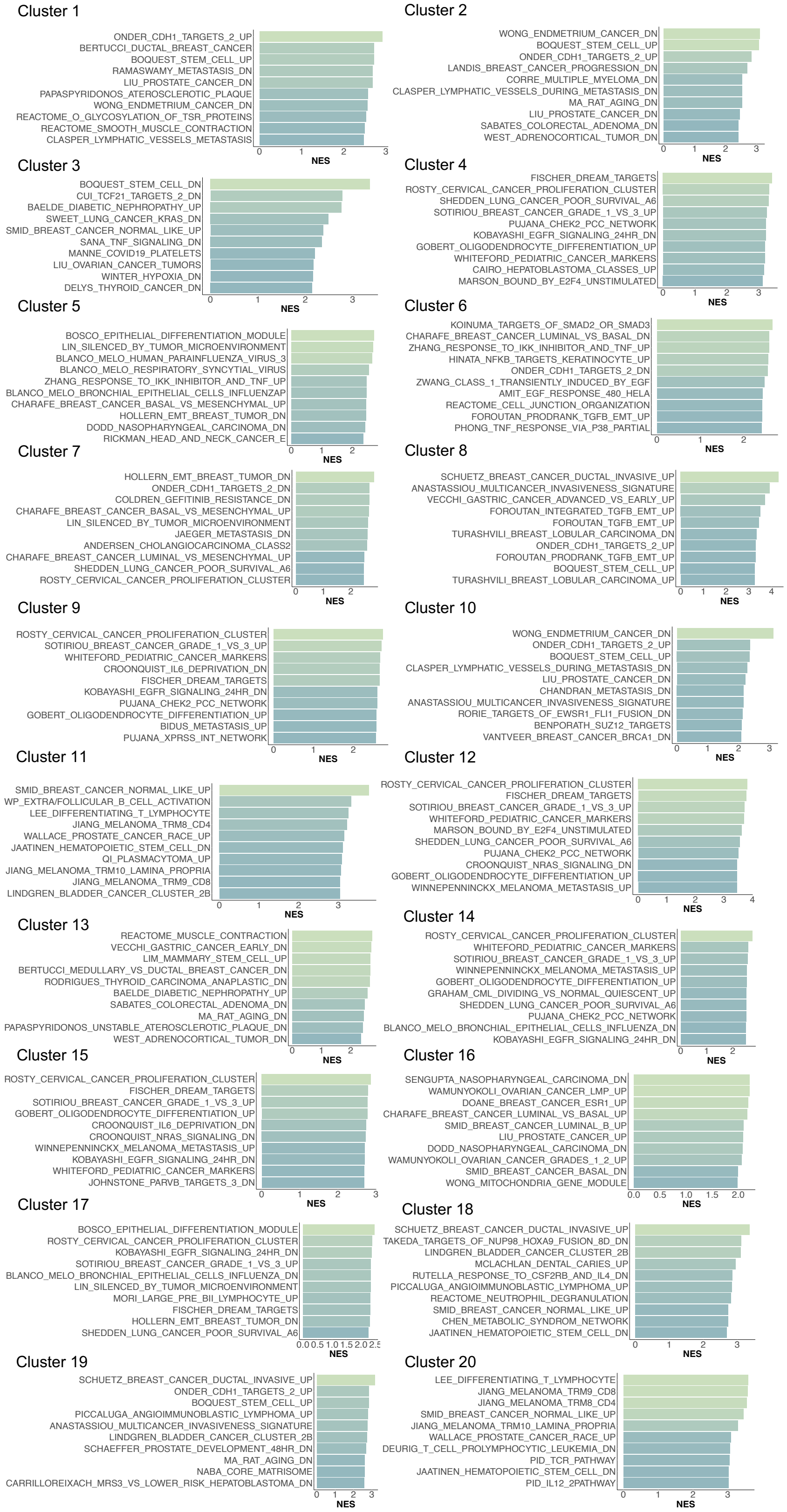

**Supplementary Figure 30. Modality specific weights for 10xXeniumPrime human cervical cancer dataset.** Here are the weights represent the Attention weights from SCIGMA's attention layer, with the higher weight representing that SCIGMA learned to attend to a specific modality more for a given cluster.

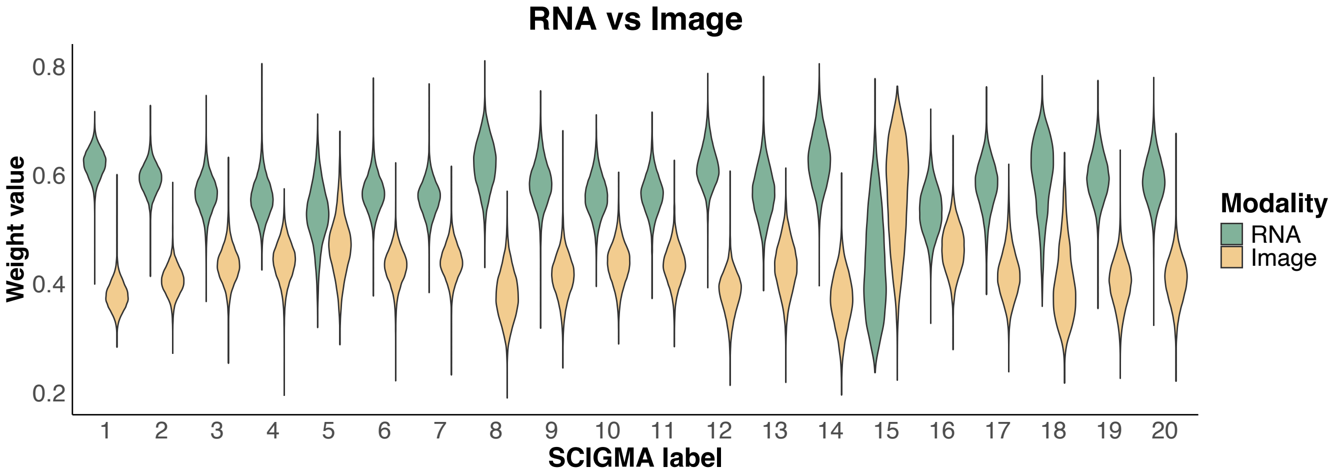

**Supplementary Figure 31. Spatial distribution of key marker genes in SCIGMA's reconstructed data versus original data for the 10xXeniumPrime human cervical cancer dataset.**

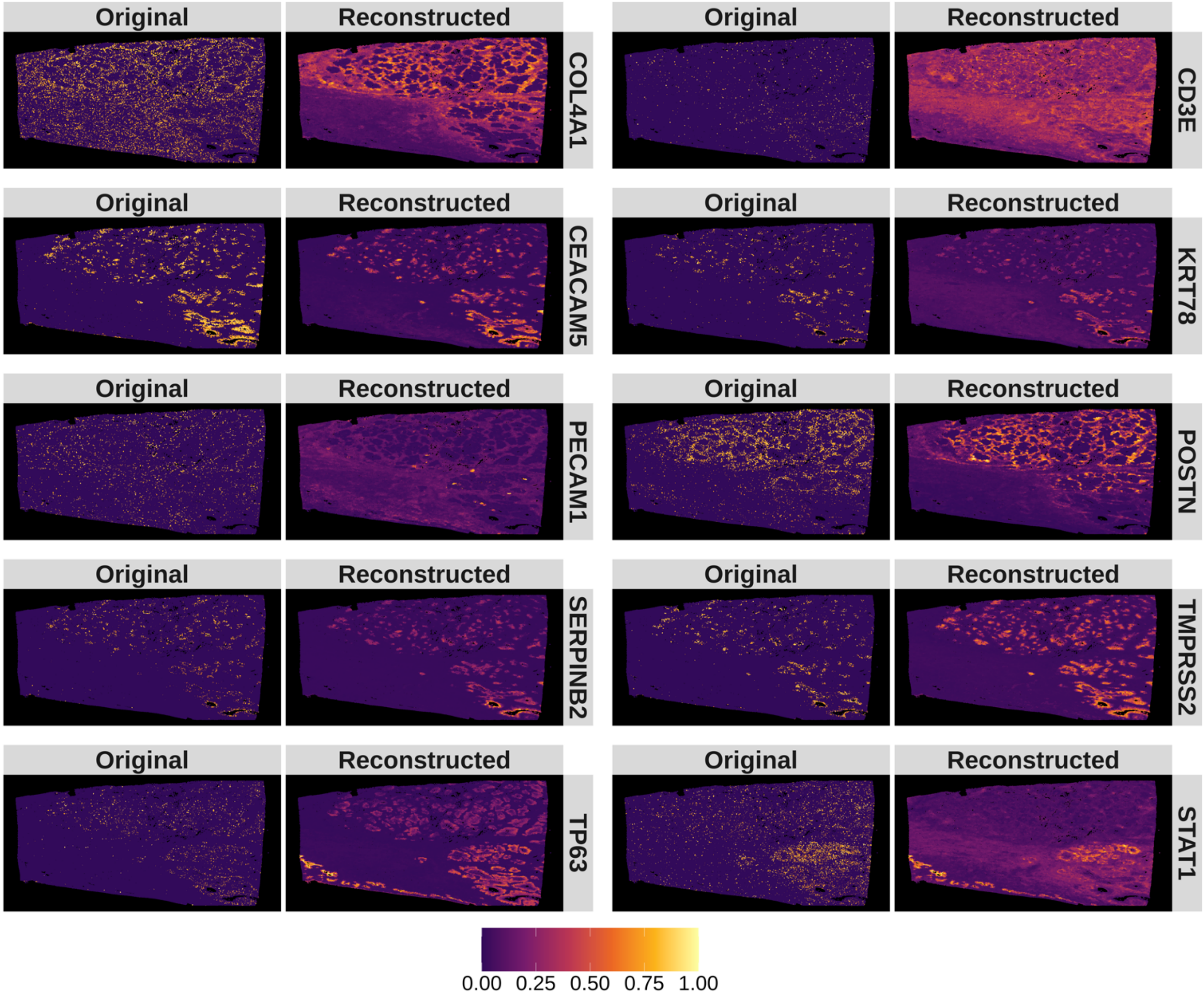

**Supplementary Figure 32. Interpretations of uncertainty estimates for the 10xXeniumPrime human cervical cancer dataset. (A)** Spatial distribution of uncertainty estimates generated by SCIGMA. **(B)** Boxplot of the total scaled expression absolute difference between RNA and imaging modalities, comparing uncertain and normal spatial locations. The y-axis represents the absolute difference. **(C)** Boxplot of the RMSE between the reconstructed and original feature matrices for RNA and imaging modalities, comparing uncertain and normal spatial locations. The y-axis represents the RMSE. **(D)** Boxplot of the Jaccard index, measuring the similarity between RNA-specific and joint embeddings, as well as imaging-specific and joint embeddings, respectively. Comparisons are made between uncertain and normal spatial locations. The y-axis represents the Jaccard index. Here, the stars ("*") indicate the significance of a one-sided Wilcoxon rank-sum test between certain and uncertain regions, where the uncertain regions are defined as the top 1%, 5%, 10%, and 20% of uncertainty values, ranked from largest to smallest.

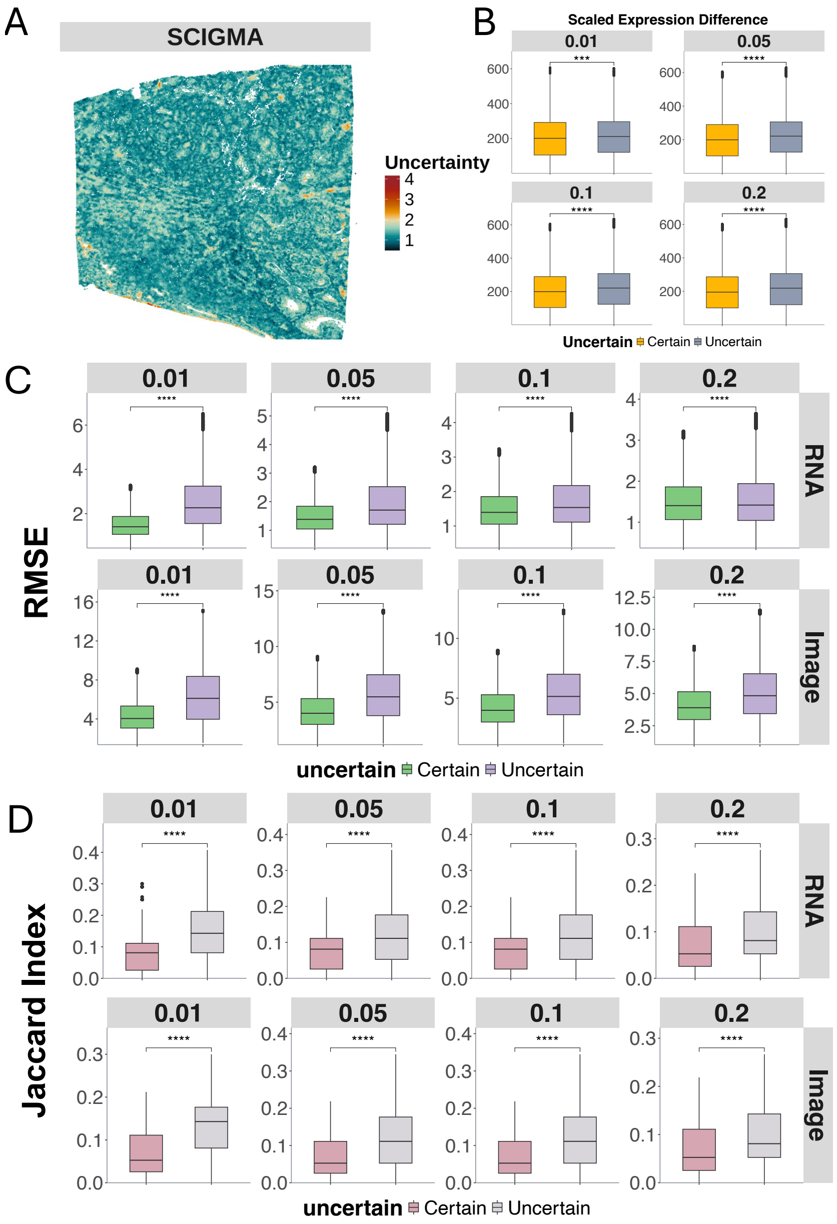

**Supplementary Figure 33. Spatial clusters identified by SCIGMA in 10xVisiumHD human colorectal cancer dataset.** This scatterplot displays the individual spatial clusters separately. The identified localization of the 20 spatial clusters by SCIGMA satisfactorily captured the tissue structure, including: tumor subregions (cluster 2, 5, 6) such as hypoxia-driven tumors (cluster 2), cancer stem cells (cluster 5) and immune evasion regions (cluster 6); and normal tissue regions such as immune cells (cluster 9).

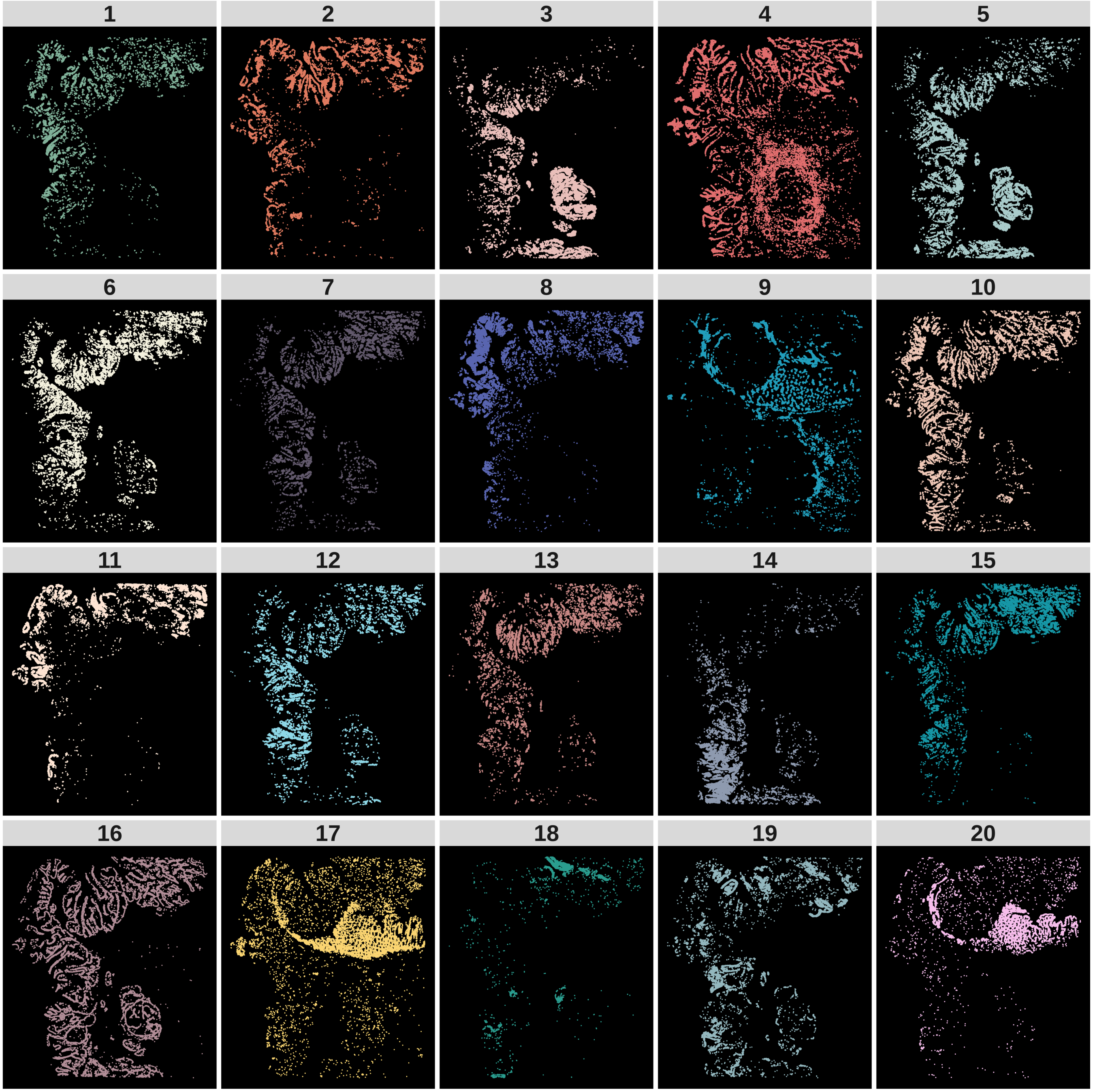

**Supplementary Figure 34. Differentially expressed genes analysis for 10xVisiumHD human colorectal cancer dataset.** Due to the space issue, we only display the top 10 selected DE genes. The red color represents a higher expression while the blue color represents a lower expression.

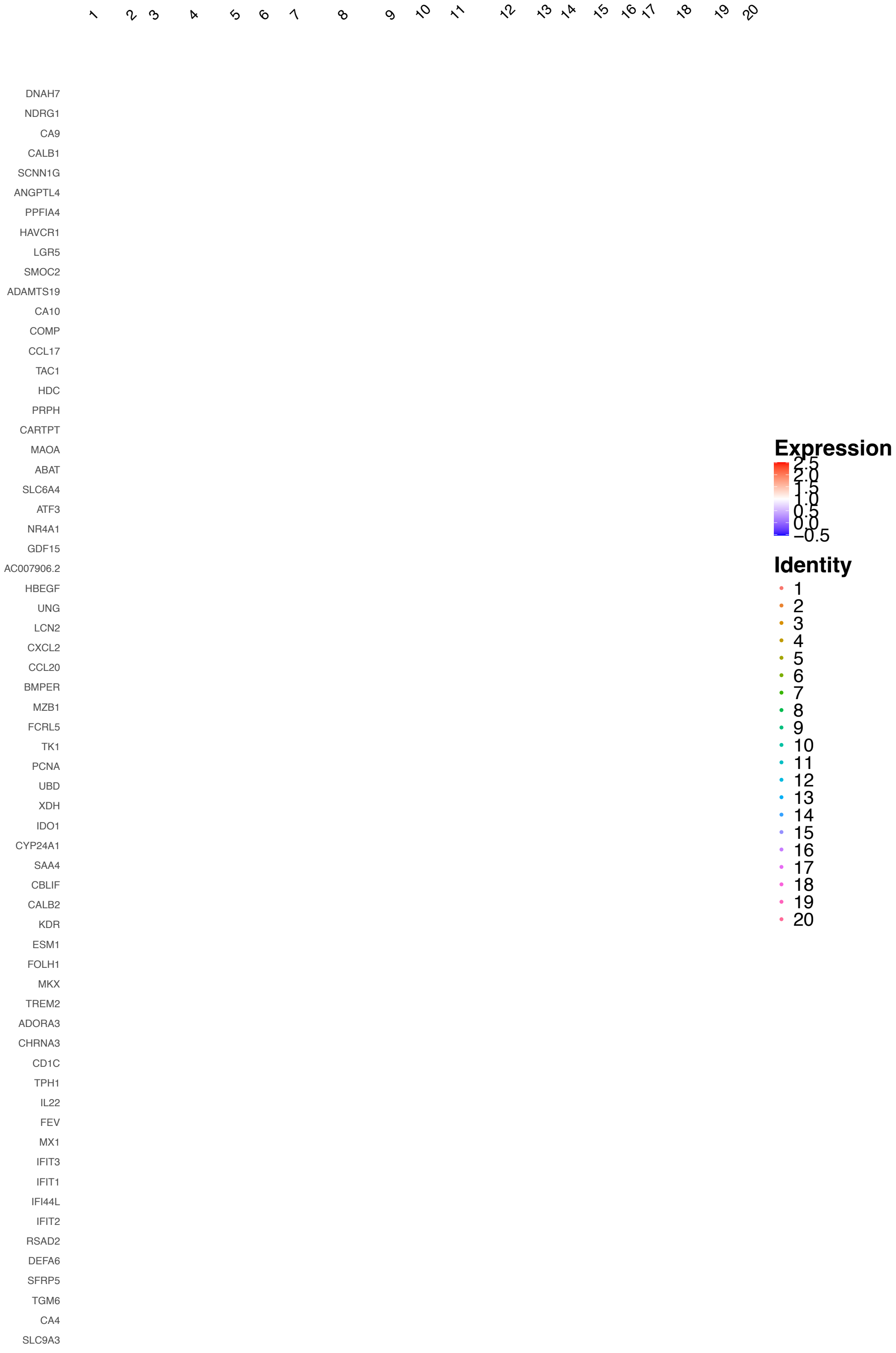

**Supplementary Figure 35. Scatter plots display the spatial distribution of important marker genes identified in the DE analysis in the 10xVisiumHD human colorectal cancer dataset.**

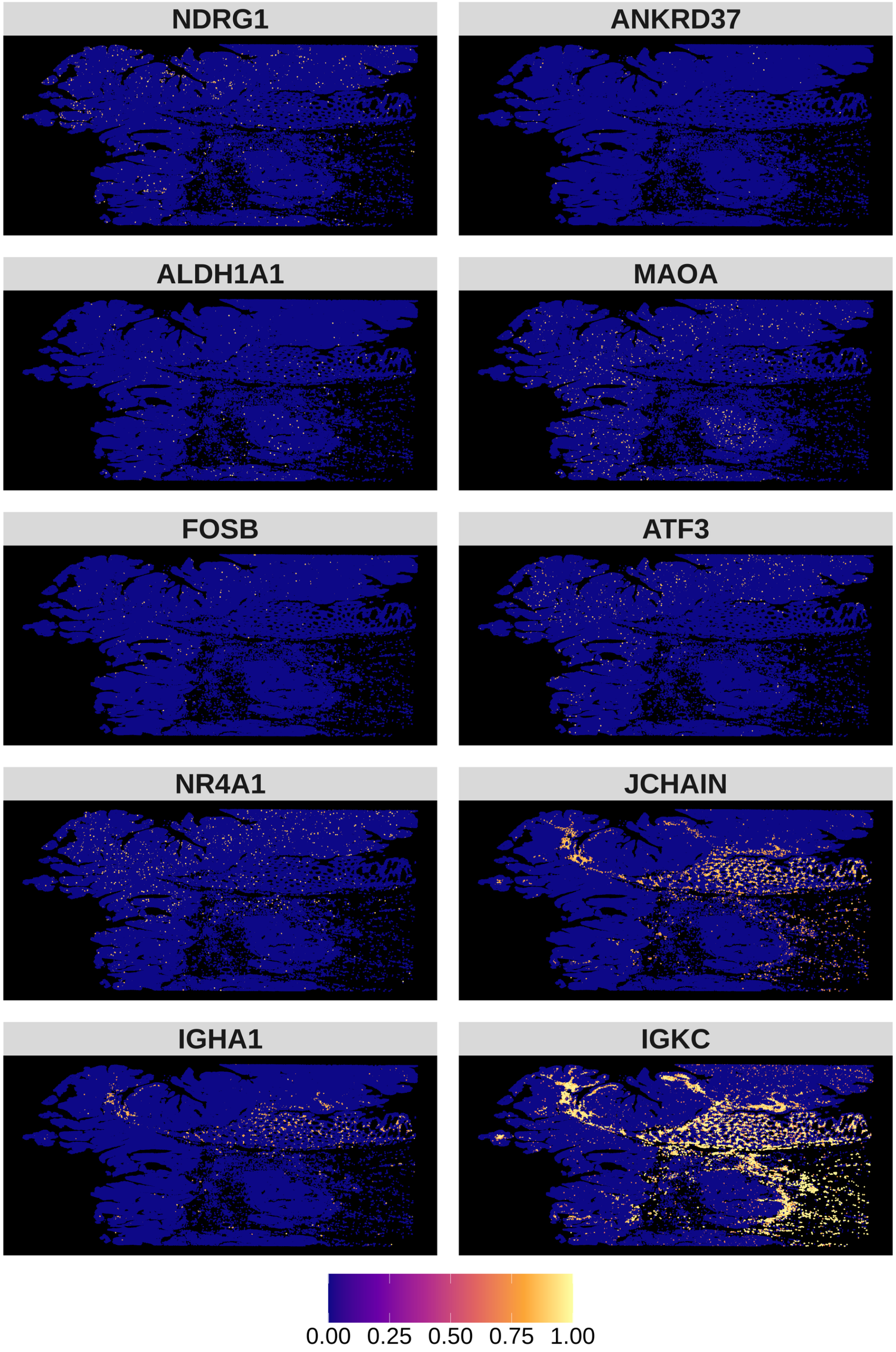

**Supplementary Figure 36. GSEA results for the 10xVisiumHD human colorectal cancer dataset.** Top 10 (if available) gene set enrichment results.

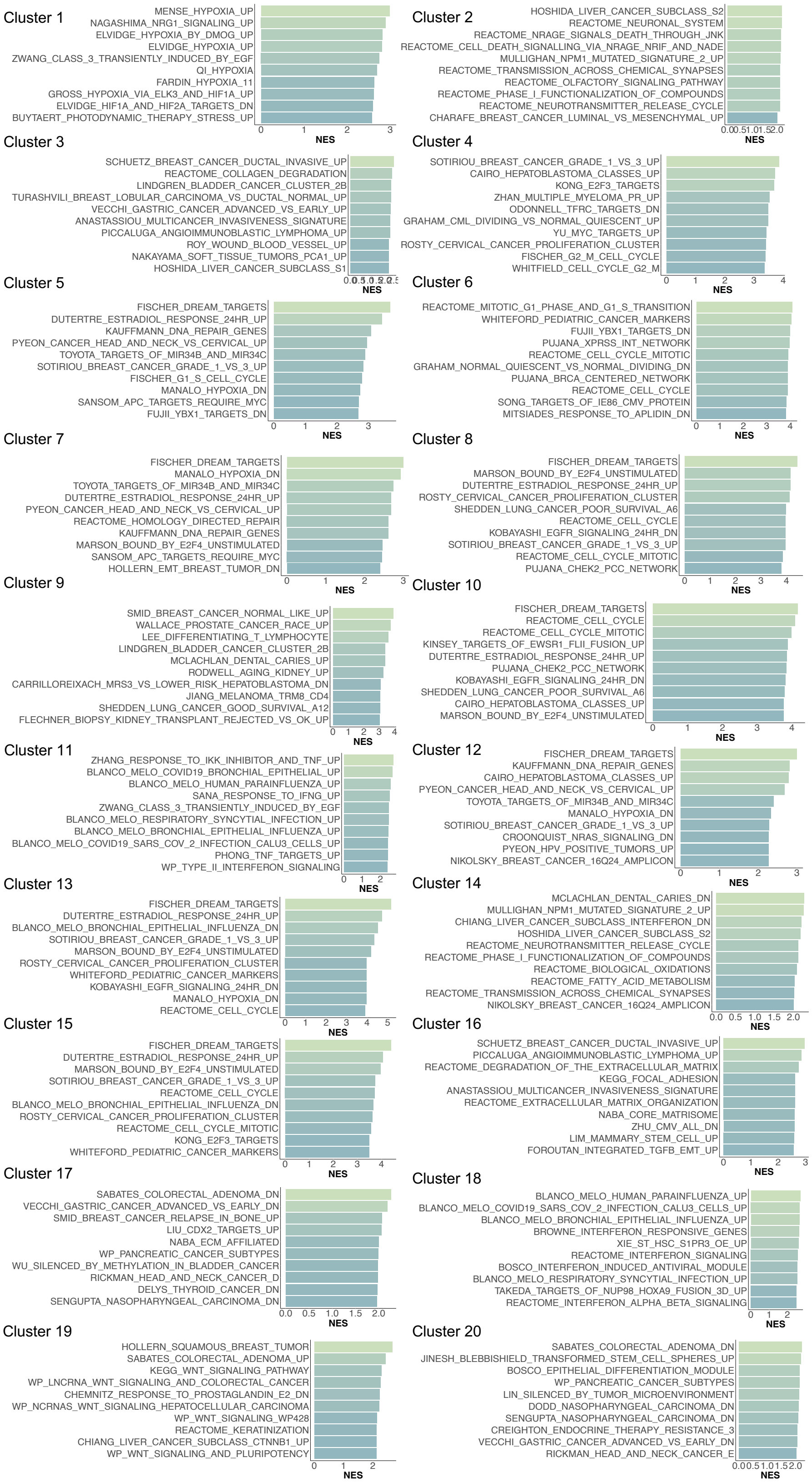

**Supplementary Figure 37. Modality specific weights for 10xVisiumHD human colorectal cancer dataset.** Here are the weights representing the Attention weights from SCIGMA's attention layer, with the higher weight representing that SCIGMA learned to attend to a specific modality more for a given cluster.

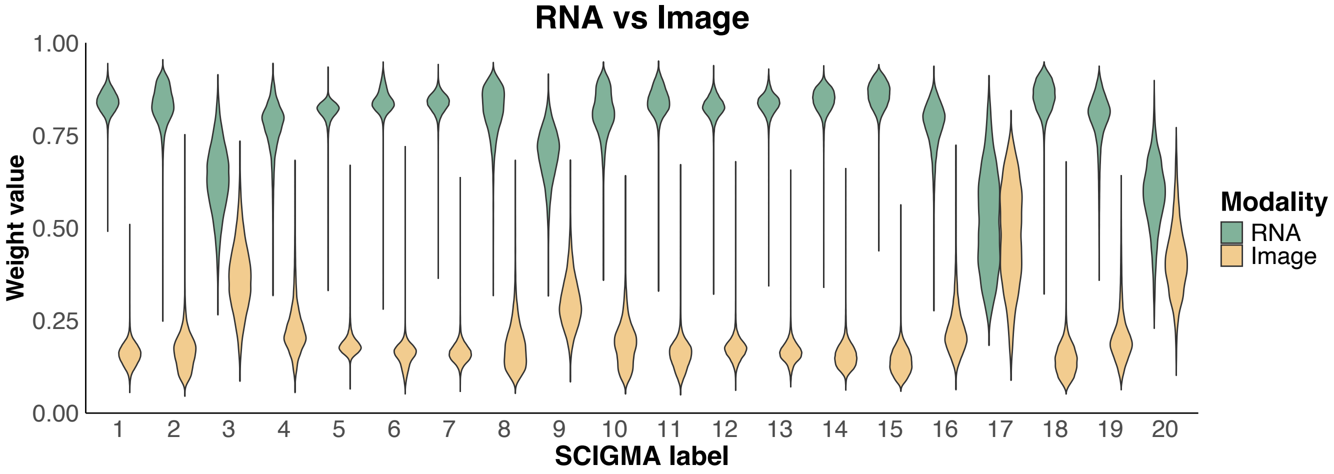

**Supplementary Figure 38. Spatial distribution of key marker features in SCIGMA's reconstructed data versus original data for the 10xXeniumPrime 10xVisiumHD human colorectal cancer dataset.**

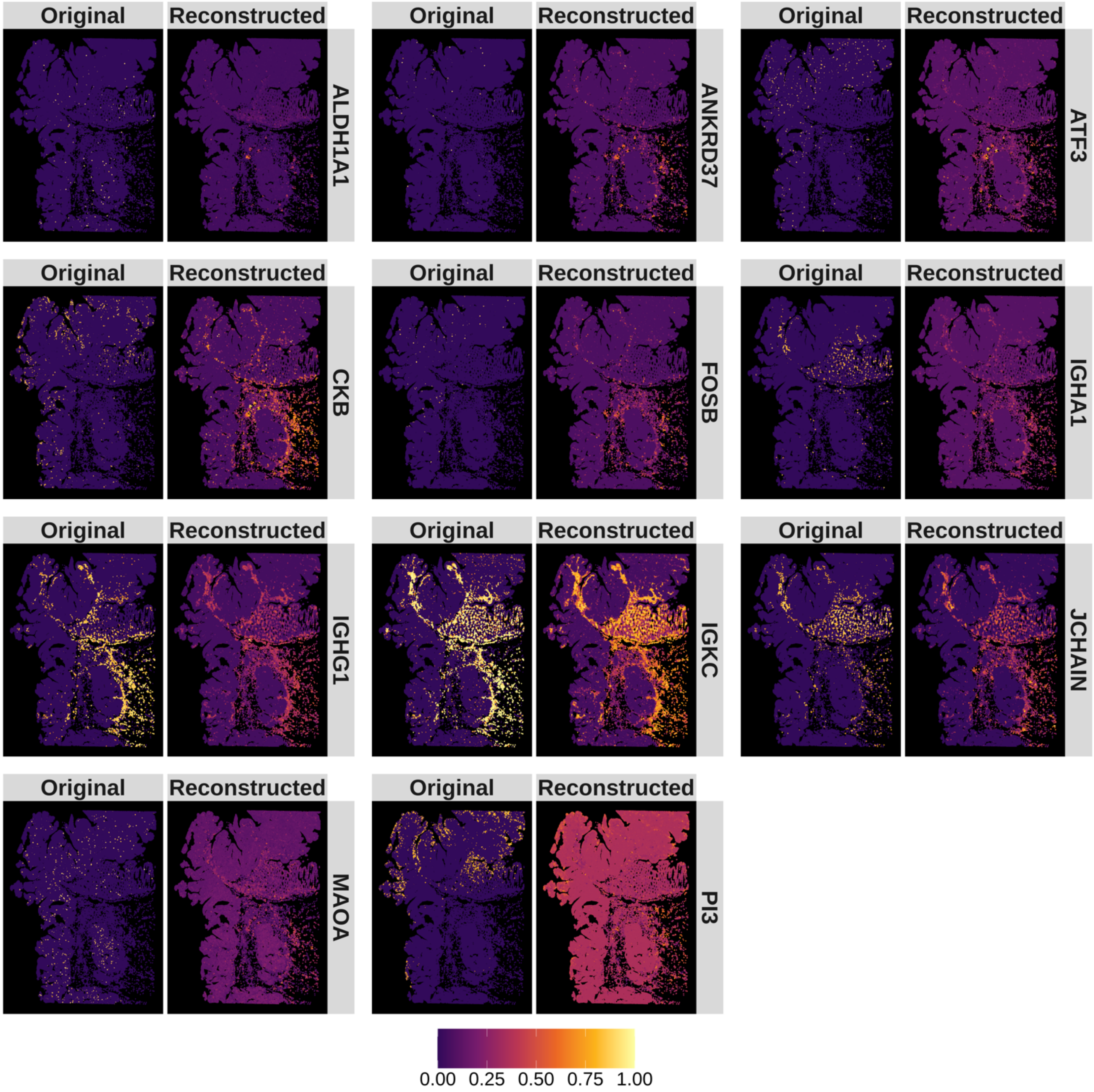

**Supplementary Figure 39. Interpretations of uncertainty estimates for the 10xVisiumHD human colorectal cancer dataset. (A)** Spatial distribution of uncertainty estimates generated by SCIGMA. **(B)** Boxplot of the total scaled expression absolute difference between RNA and imaging modalities, comparing uncertain and normal spatial locations. The y-axis represents the absolute difference. **(C)** Boxplot of the RMSE between the reconstructed and original feature matrices for RNA and imaging modalities, comparing uncertain and normal spatial locations. The y-axis represents the RMSE. **(D)** Boxplot of the Jaccard index, measuring the similarity between RNA-specific and joint embeddings, as well as imaging-specific and joint embeddings, respectively. Comparisons are made between uncertain and normal spatial locations. The y-axis represents the Jaccard index. Here, the stars ("*") indicate the significance of a one-sided Wilcoxon rank-sum test between certain and uncertain regions, where the uncertain regions are defined as the top 1%, 5%, 10%, and 20% of uncertainty values, ranked from largest to smallest.

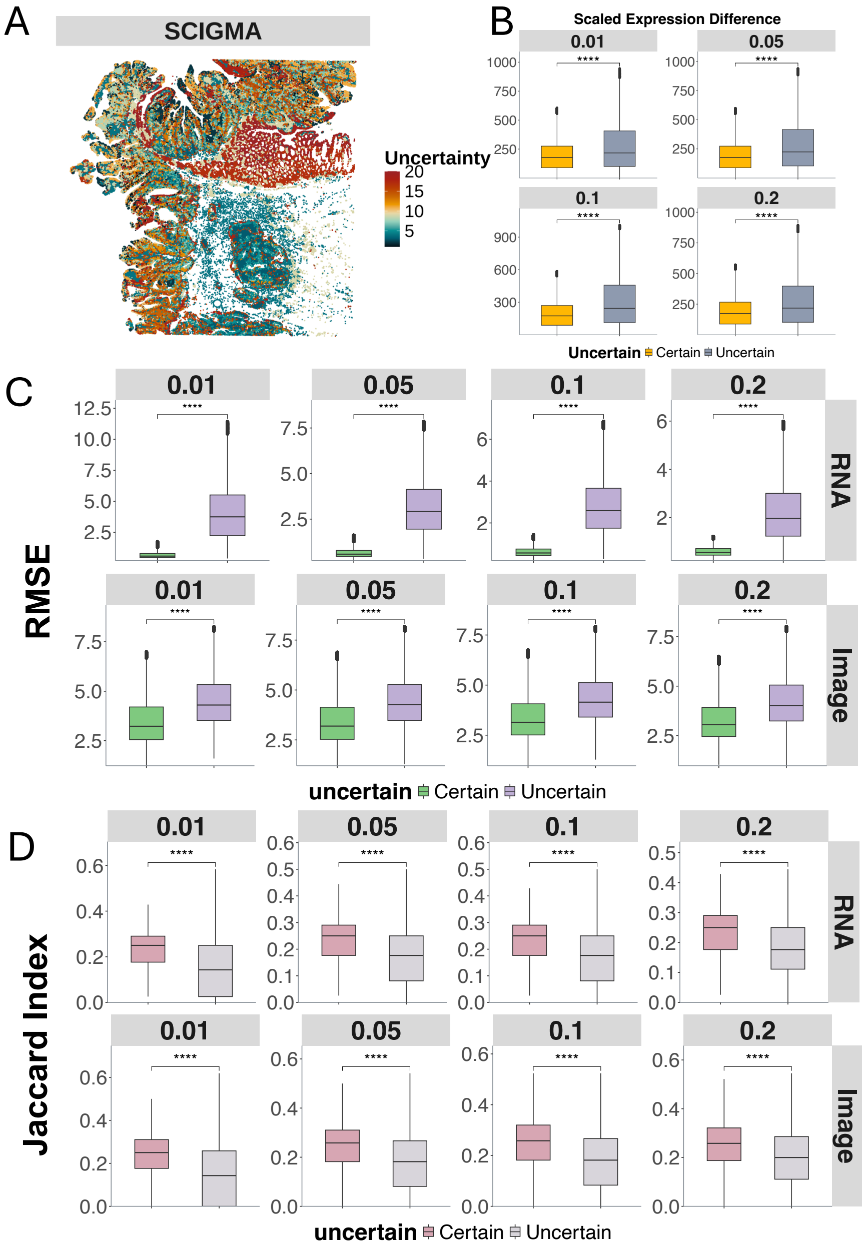

**Supplementary Figure 40. Spatial clusters identified by SCIGMA in 10xVisiumHD mouse small intestine dataset.** This scatterplot displays the individual spatial clusters separately. The identified localization of the 20 spatial clusters by SCIGMA satisfactorily captured the tissue structure, including: villus epithelium (cluster 13); lamina propria (cluster 6); and intestinal crypt (cluster 7, 15).

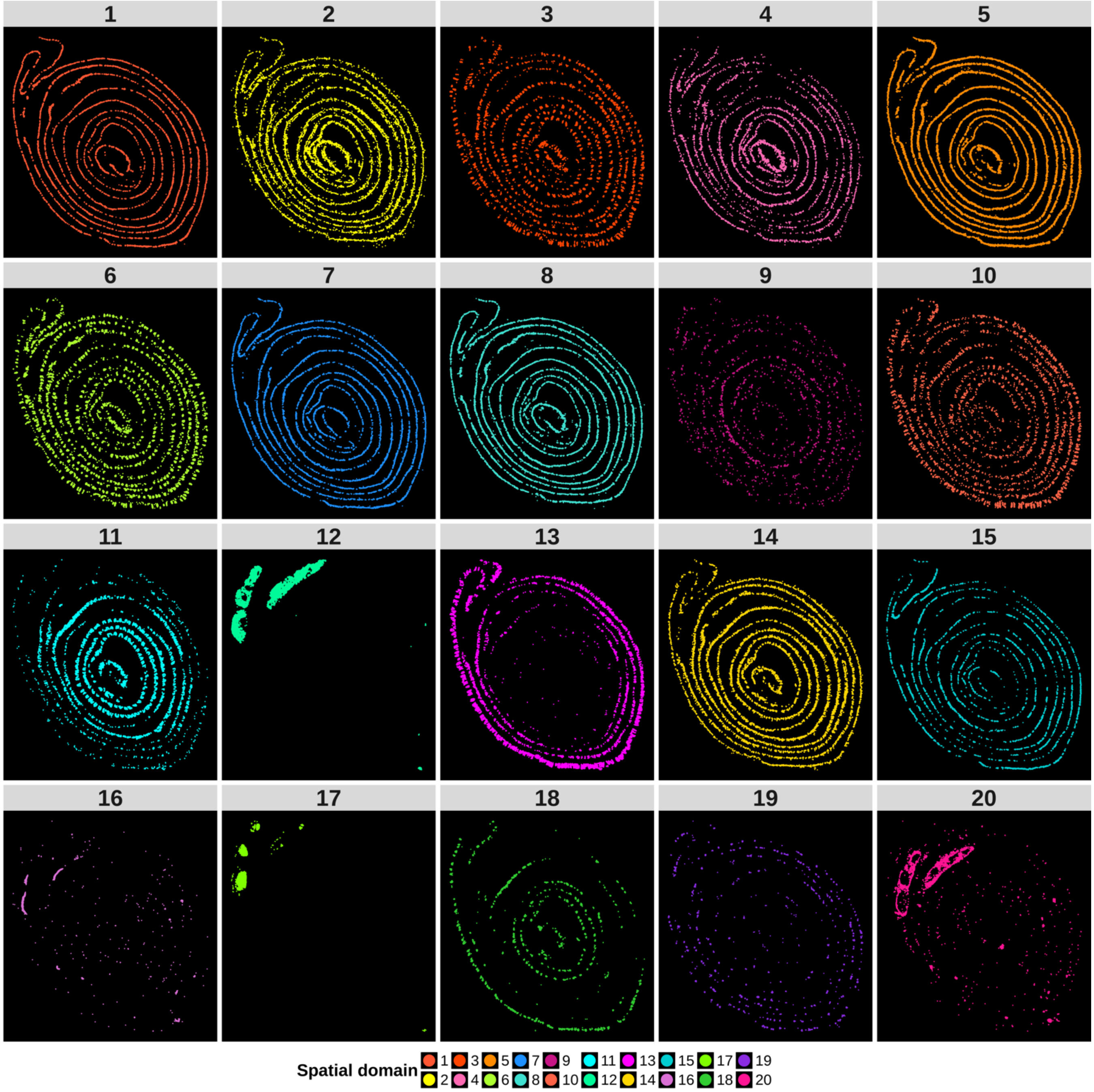

**Supplementary Figure 41. Differentially expressed genes analysis for 10xVisiumHD mouse small intestine dataset.** Due to the space issue, we only display the top 10 selected DE genes. The red color represents a higher expression while the blue color represents a lower expression.

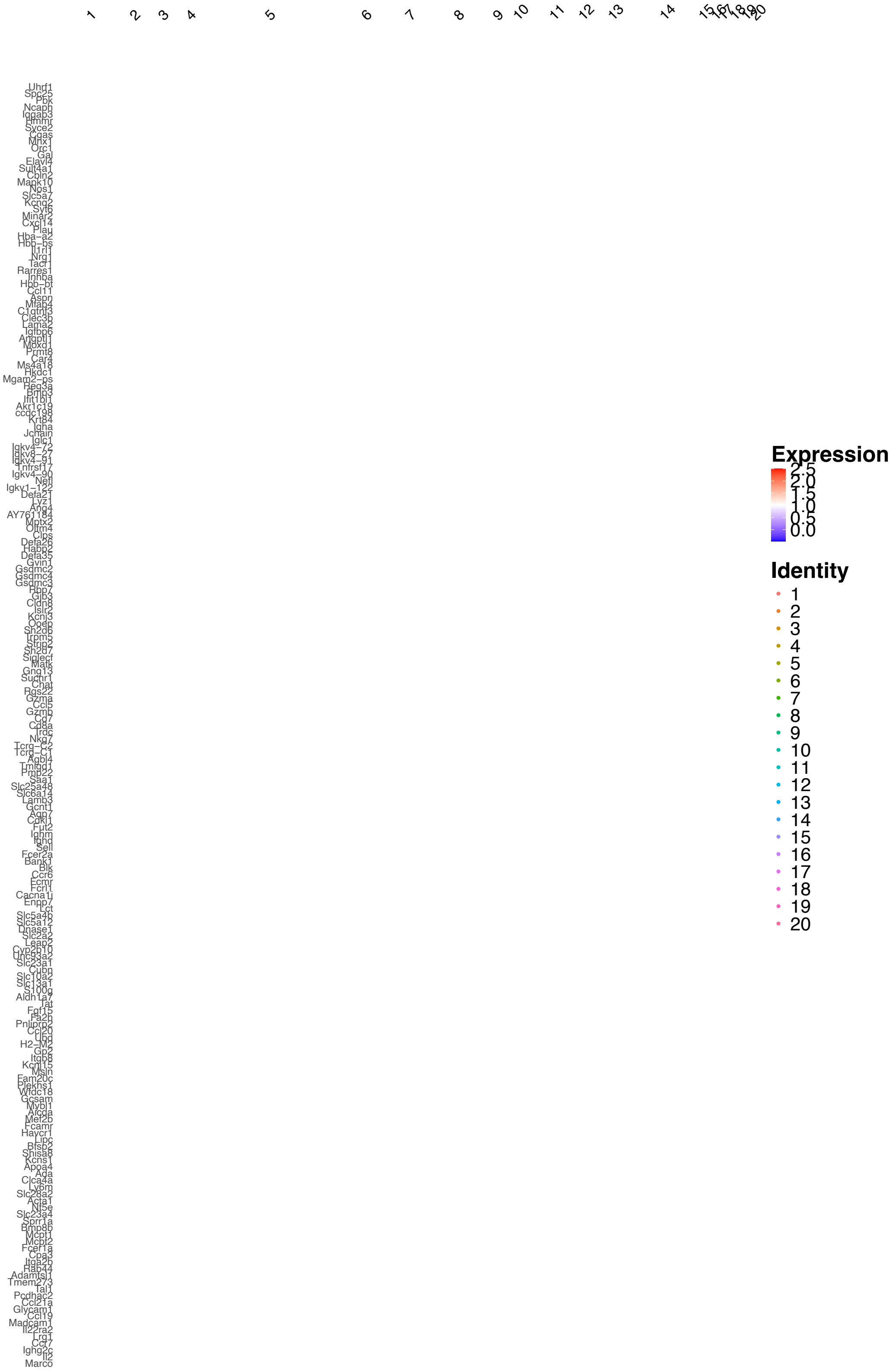

**Supplementary Figure 42. Scatter plots display the spatial distribution of important marker genes identified in the DE analysis in the 10xVisiumHD mouse small intestine dataset.**

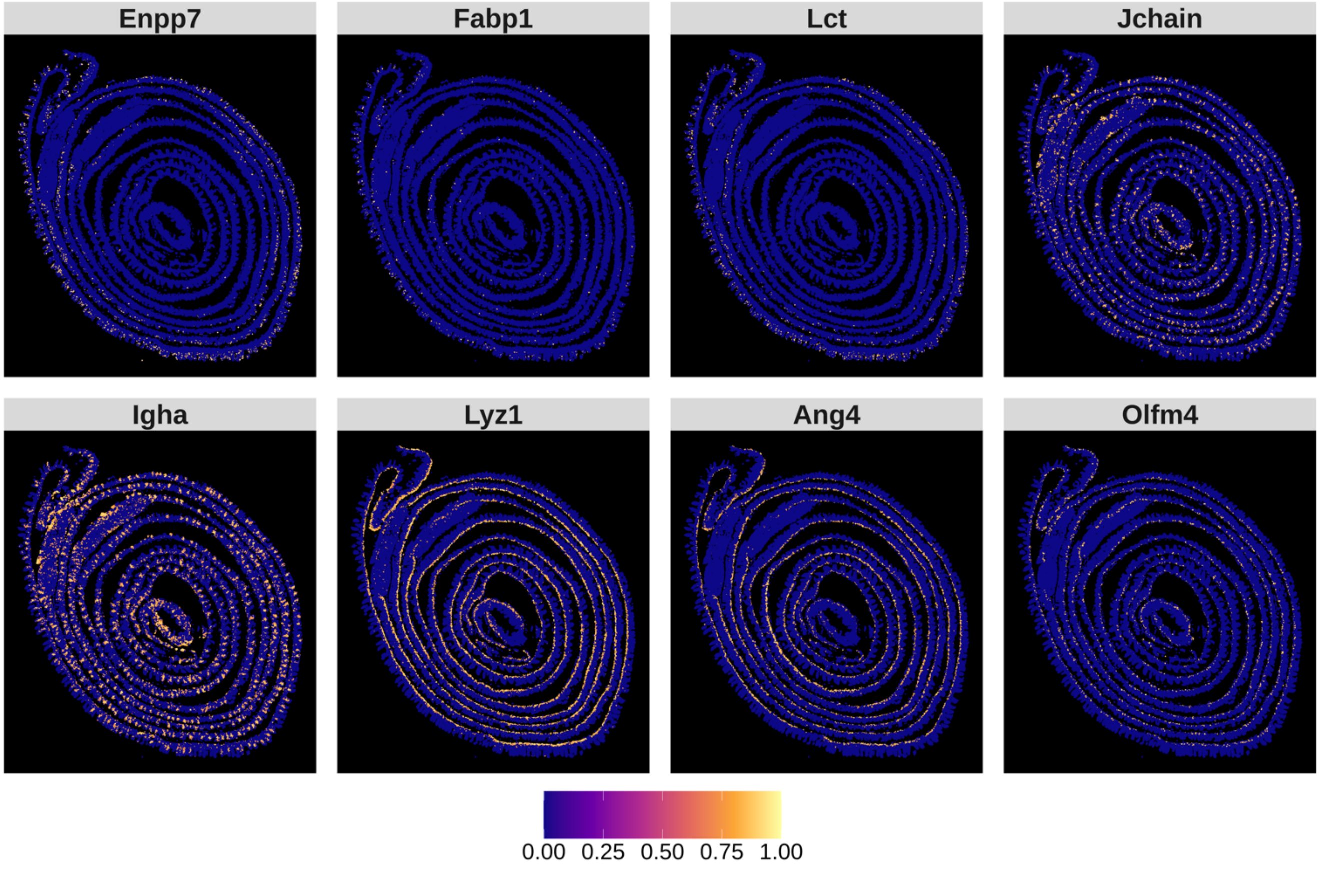

**Supplementary Figure 43. GSEA results for the 10xVisiumHD mouse small intestine cancer dataset.** Top 10 (if available) gene set enrichment results. Details on filtering provided in Supplementary Note 3.

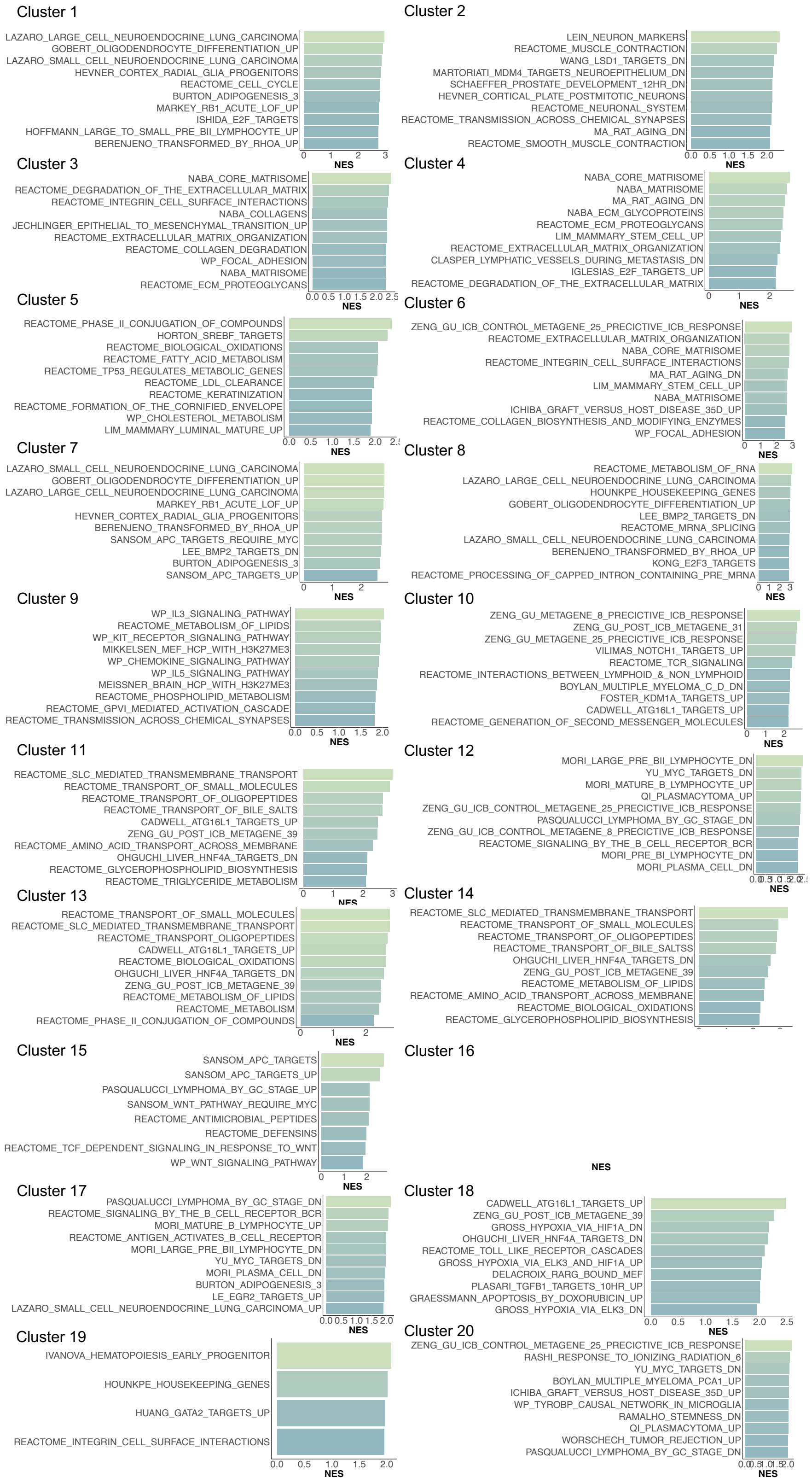

**Supplementary Figure 44. Modality specific weights for 10xVisiumHD mouse small intestine dataset.** Here are the weights representing the Attention weights from SCIGMA's attention layer, with the higher weight representing that SCIGMA learned to attend to a specific modality more for a given cluster.

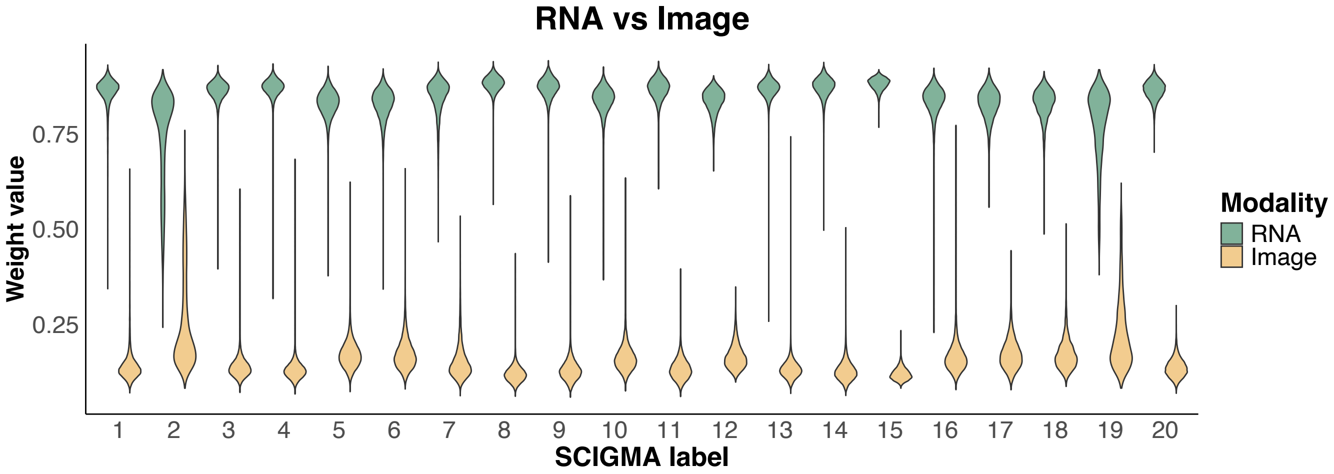

**Supplementary Figure 45. Spatial distribution of key marker features in SCIGMA's reconstructed data versus original data for the 10xVisiumHD mouse small intestine dataset.**

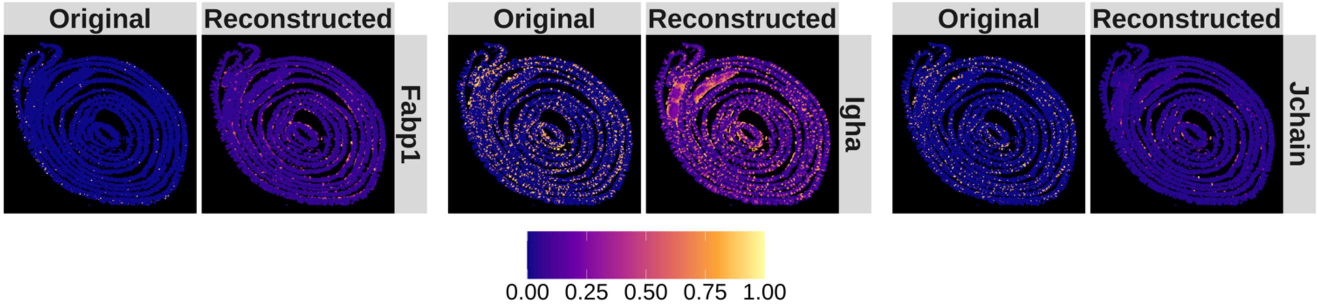

**Supplementary Figure 46. Uncertainty interpretability analysis for the 10xVisiumHD mouse small intestine dataset. (A)** Spatial distribution of uncertainty estimates generated by SCIGMA. **(B)** Boxplot of the total scaled expression absolute difference between RNA and imaging modalities, comparing uncertain and normal spatial locations. The y-axis represents the absolute difference. **(C)** Boxplot of the RMSE between the reconstructed and original feature matrices for RNA and imaging modalities, comparing uncertain and normal spatial locations. The y-axis represents the RMSE. **(D)** Boxplot of the Jaccard index, measuring the similarity between RNA-specific and joint embeddings, as well as imaging-specific and joint embeddings, respectively. Comparisons are made between uncertain and normal spatial locations. The y-axis represents the Jaccard index. Here, the stars ("*") indicate the significance of a one-sided Wilcoxon rank-sum test between certain and uncertain regions, where the uncertain regions are defined as the top 1%, 5%, 10%, and 20% of uncertainty values, ranked from largest to smallest.

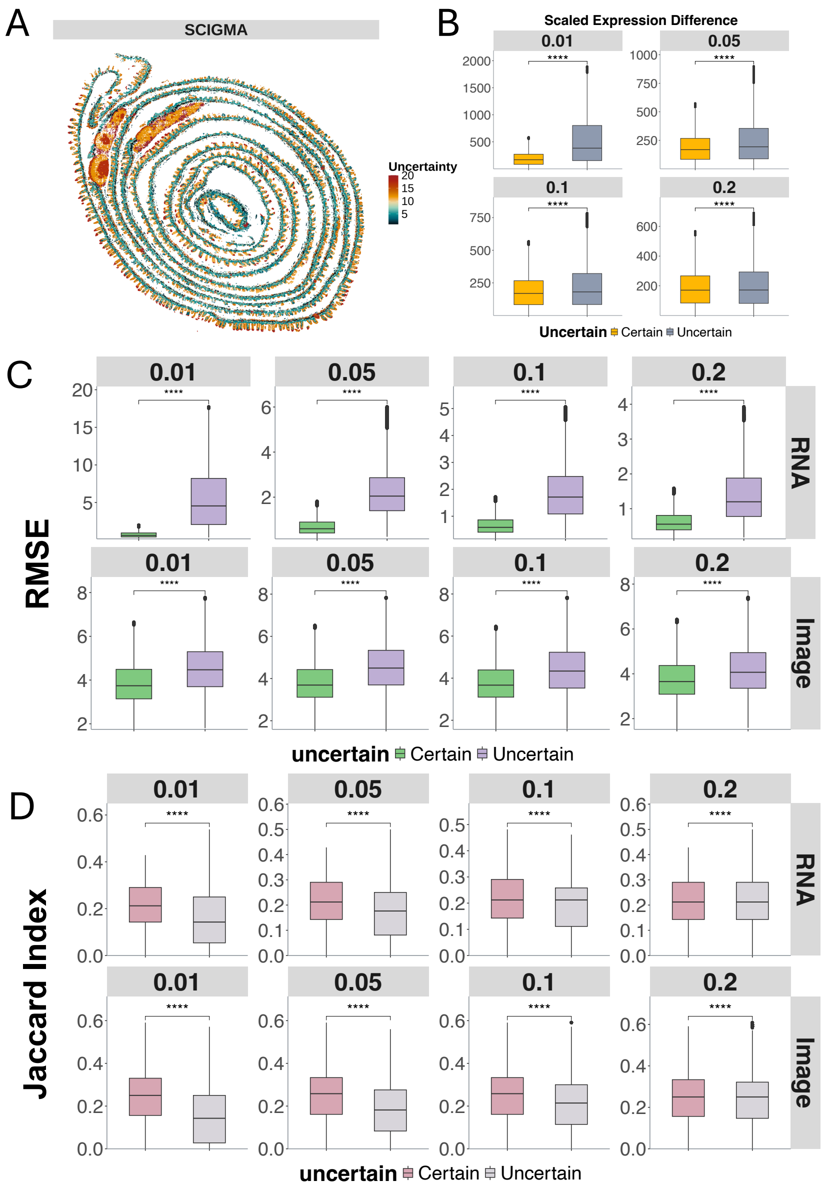

**Supplementary Figure 47. Spatial clusters identified by SCIGMA in spatial mux-seq 5-month mouse brain dataset.** This scatterplot displays the individual spatial clusters separately. The identified localization of the 17 spatial clusters by SCIGMA satisfactorily captured the tissue structure, including: cortex layers (clusters 3, 6, 8, 12, 13, 16), CCG (cluster 9), CP (cluster 5, 11), ACB (cluster 2), LSR (cluster 1), and ACO (cluster 10).

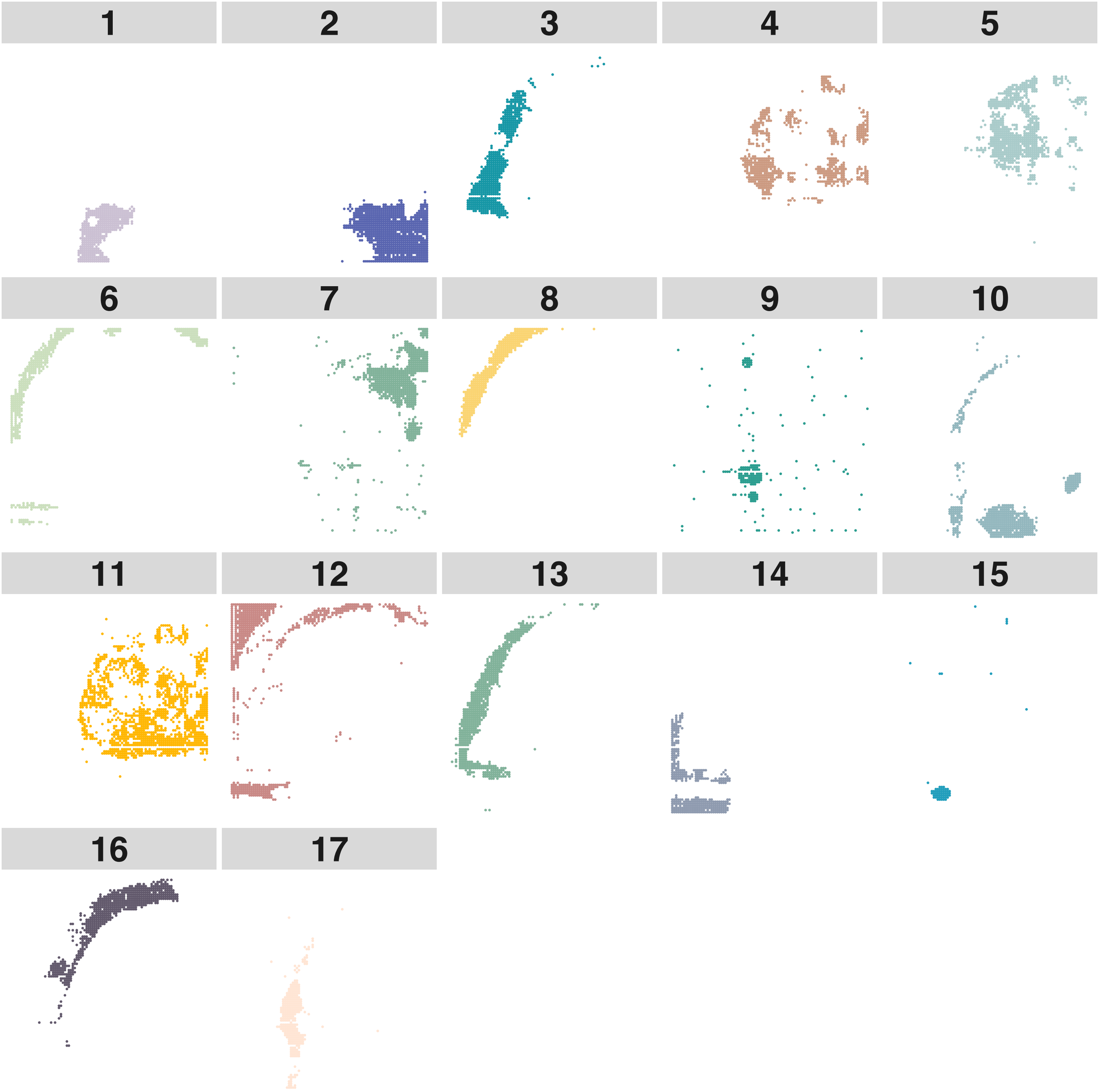

**Supplementary Figure 48. DEG results for spatial mux-seq mouse brain dataset.** Due to the space issue, we only display the top 10 selected DE genes. The red color represents a higher expression while the blue color represents a lower expression.

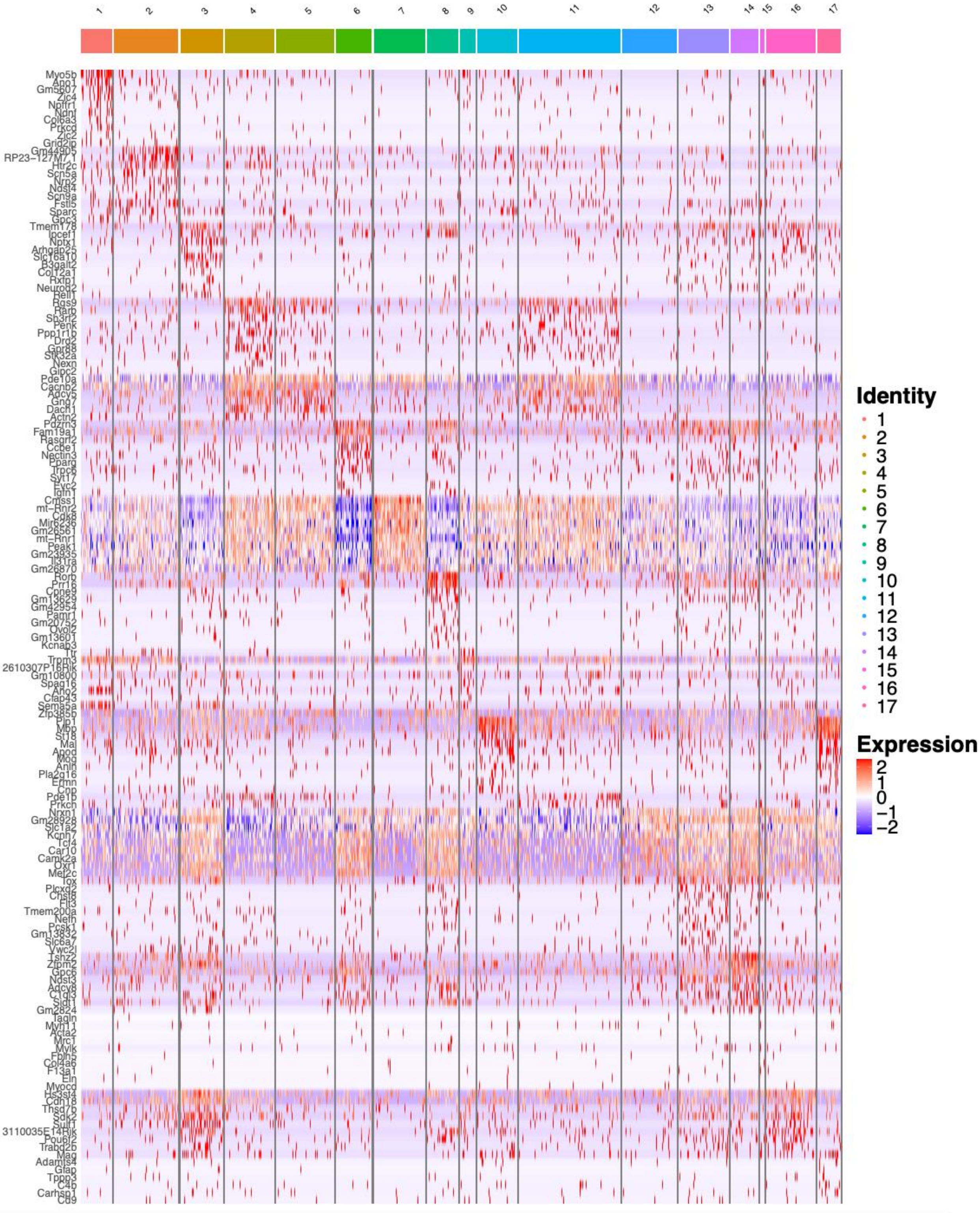

**Supplementary Figure 49. Scatter plots displaying the spatial distribution of important marker genes identified in the DE analysis in the spatial mux-seq mouse brain dataset.**

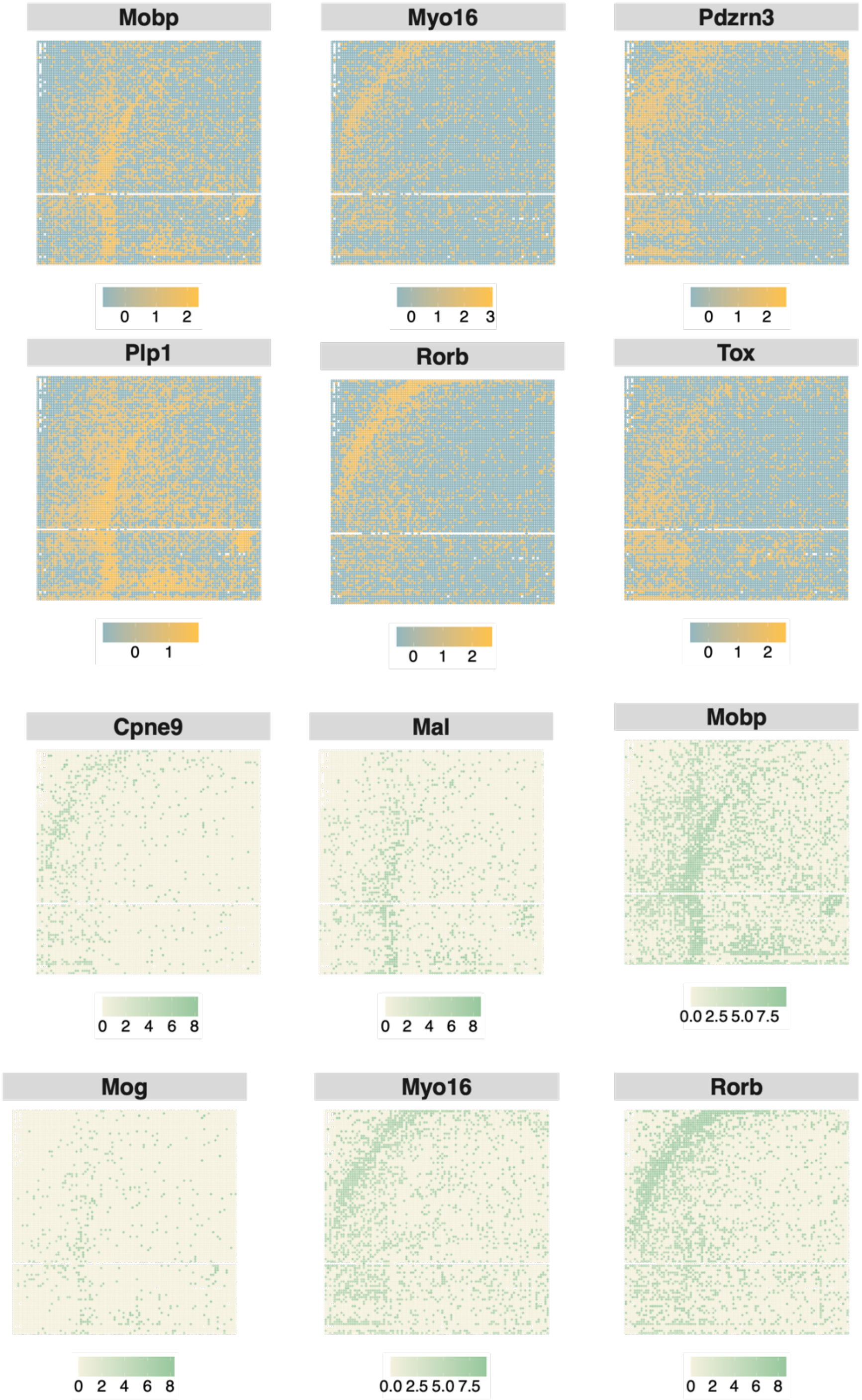

**Supplementary Figure 50. GSEA results for the 10xVisiumHD spatial mux-seq mouse brain dataset.** Top 10 (if available) gene set enrichment results. Details on filtering provided in Supplementary Note 3.

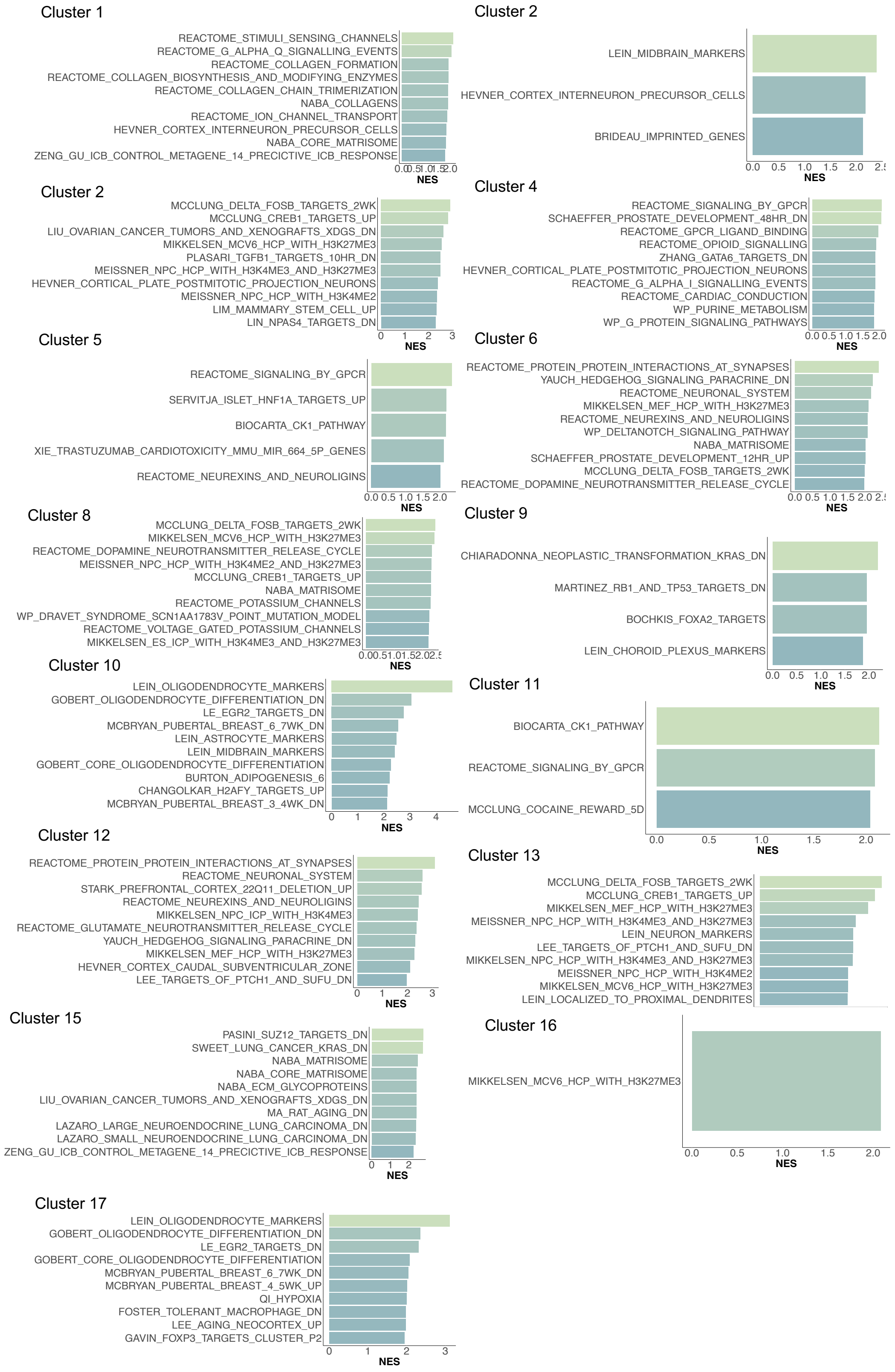

**Supplementary Figure 51. Modality specific weights for spatial mux-seq mouse brain dataset.** Here are the weights representing the Attention weights from SCIGMA's attention layer, with the higher weight representing that SCIGMA learned to attend to a specific modality more for a given cluster.

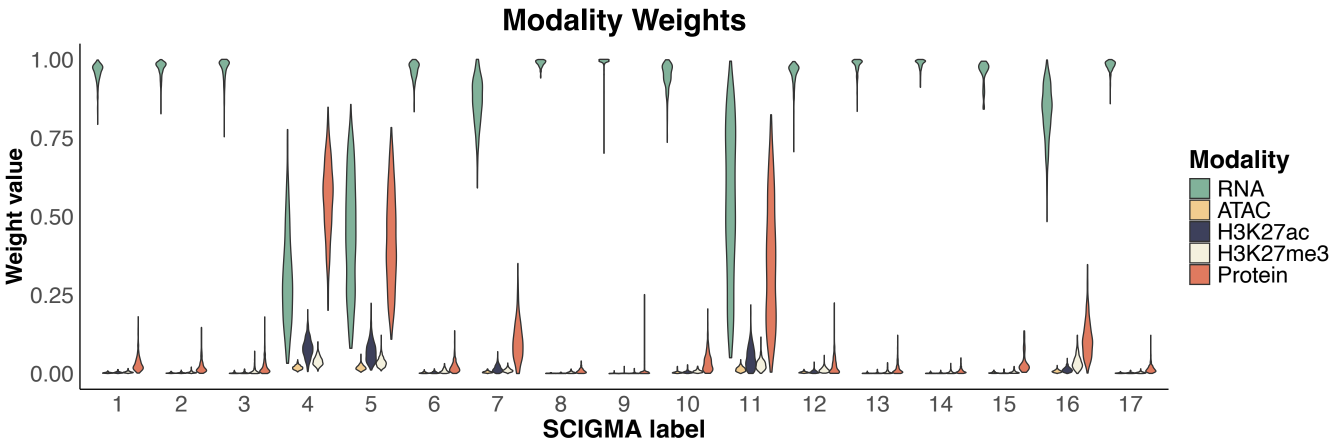

**Supplementary Figure 52. Spatial distribution of key marker features (eg., genes and proteins) in SCIGMA's reconstructed data versus original data for the spatial mux-seq mouse brain dataset.**

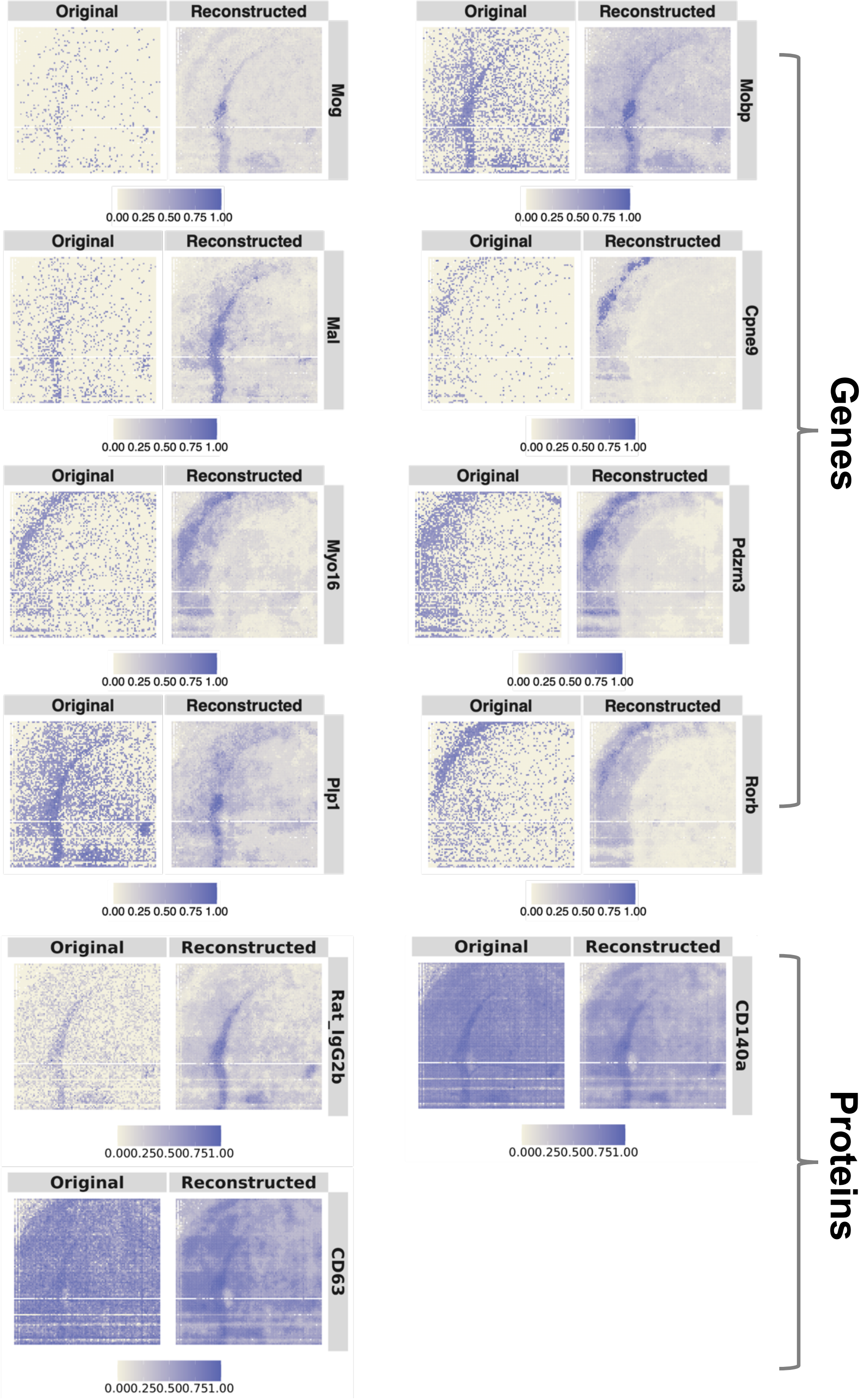

**Supplementary Figure 53. Interpretations of uncertainty estimates for the spatial mux-seq mouse brain dataset. (A)** Spatial distribution of uncertainty estimates generated by SCIGMA. **(B)** Boxplot of the total scaled expression absolute difference between RNA and protein modalities, comparing uncertain and normal spatial locations. The y-axis represents the absolute difference. **(C)** Boxplot of the RMSE between the reconstructed and original feature matrices for RNA and protein modalities, comparing uncertain and normal spatial locations. The y-axis represents the RMSE. **(D)** Boxplot of the Jaccard index, measuring the similarity between RNA-specific and joint embeddings, as well as protein-specific and joint embeddings, respectively. Comparisons are made between uncertain and normal spatial locations. The y-axis represents the Jaccard index. Here, the stars ("*") indicate the significance of a one-sided Wilcoxon rank-sum test between certain and uncertain regions, where the uncertain regions are defined as the top 1%, 5%, 10%, and 20% of uncertainty values, ranked from largest to smallest.

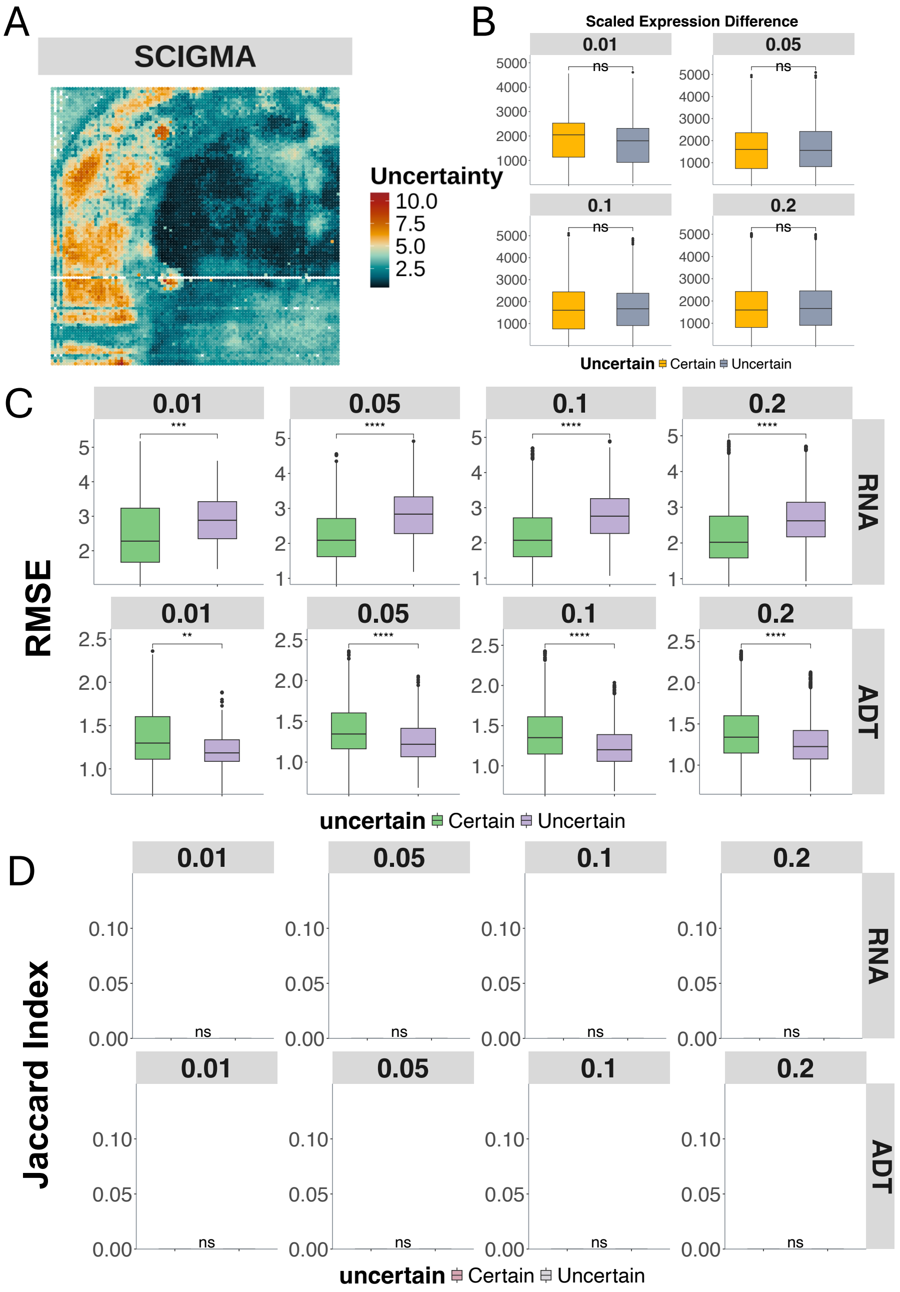

**Supplementary Figure 54. Analysis of the spatial epigenome-transcriptome (CUT&Tag-seq) H3K27ac dataset.** (**A**) Spatial clusters identified by single-modality and multi-modal integration methods, including Seurat, MOFA+, MultiVI, SpatialGlue, and SCIGMA. (**B**) Boxplots displaying Moran's I values across methods, quantifying spatial autocorrelation. Each box spans the first to third quartiles, with the median represented by a horizontal line; whiskers extend to 1.5 times the interquartile range. (**C**) Jaccard similarity scores comparing preservation of modality-specific relationships in the joint representation across different methods. (**D**) Boxplots displaying pairwise ARI and NMI, assessing the consistency of clustering results across 20 random seeds for deep learning methods. SCIGMA successfully identified the cortex layers (clusters 5, 3, 13), CCG (cluster 6), VL (cluster 14), CP (cluster 1), ACB (cluster 8), and LSR (cluster 11) regions, showing improvements over all of the benchmarked methods. SCIGMA displayed the highest Moran's I and Jaccard similarity, and it showed the best consistency among the deep learning frameworks.

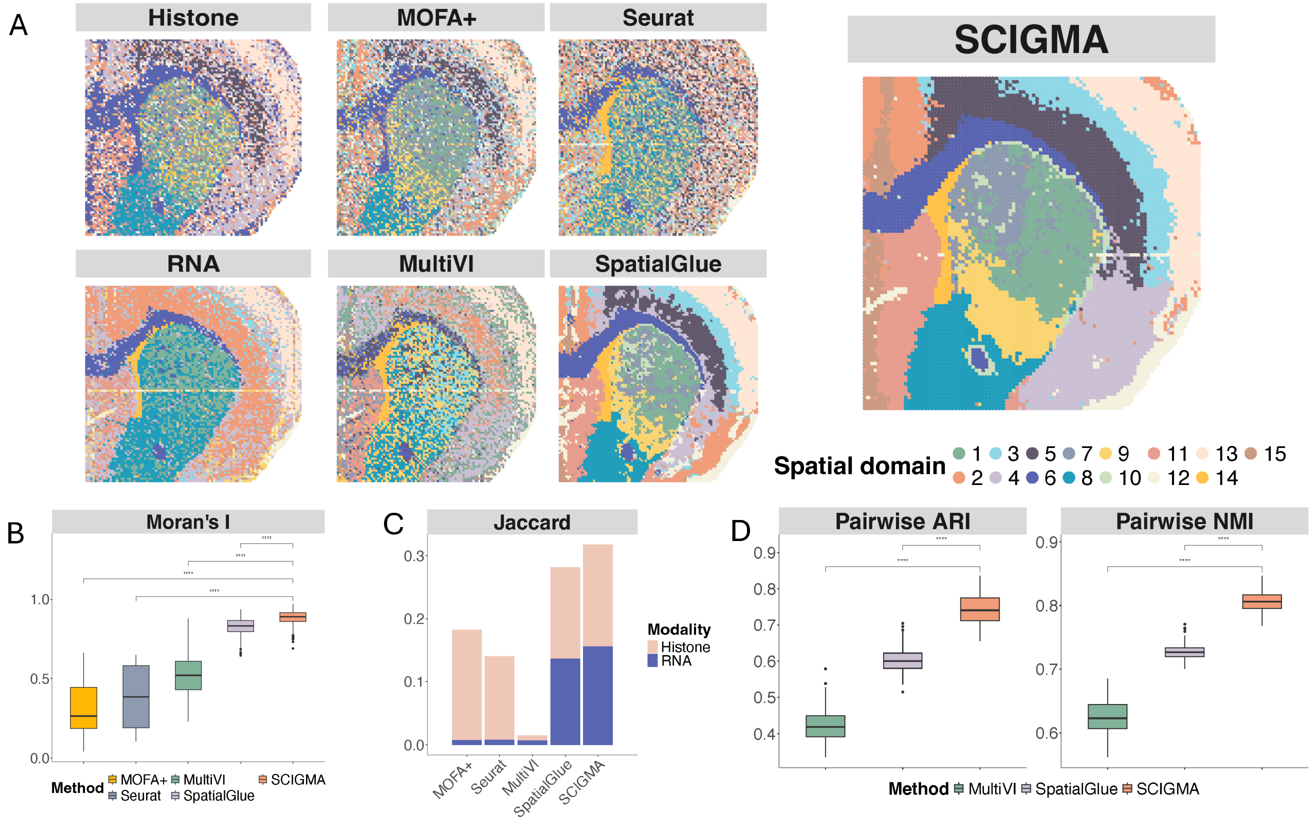

**Supplementary Figure 55. Analysis of the spatial epigenome-transcriptome (CUT&Tag-seq) H3K27me3 dataset.** (A) Spatial clusters identified by single-modality and multi-modal integration methods, including Seurat, MOFA+, MultiVI, SpatialGlue, and SCIGMA. (**B**) Boxplots displaying Moran's I values across methods, quantifying spatial autocorrelation. Each box spans the first to third quartiles, with the median represented by a horizontal line; whiskers extend to 1.5 times the interquartile range. (**C**) Jaccard similarity scores comparing preservation of modality-specific relationships in the joint representation across different methods. (**D**) Boxplots displaying pairwise ARI and NMI, assessing the consistency of clustering results across 20 random seeds for each method. SCIGMA successfully identified the cortex layers (clusters 15, 5, 7, 6), CCG (cluster 10), VL (cluster 8), CP (cluster 7), ACB (cluster 9), and LSR (cluster 3) regions, showing improvements over all of the benchmarked methods. SCIGMA displayed the highest Moran's I and Jaccard similarity, and it showed the best consistency among the deep learning frameworks.

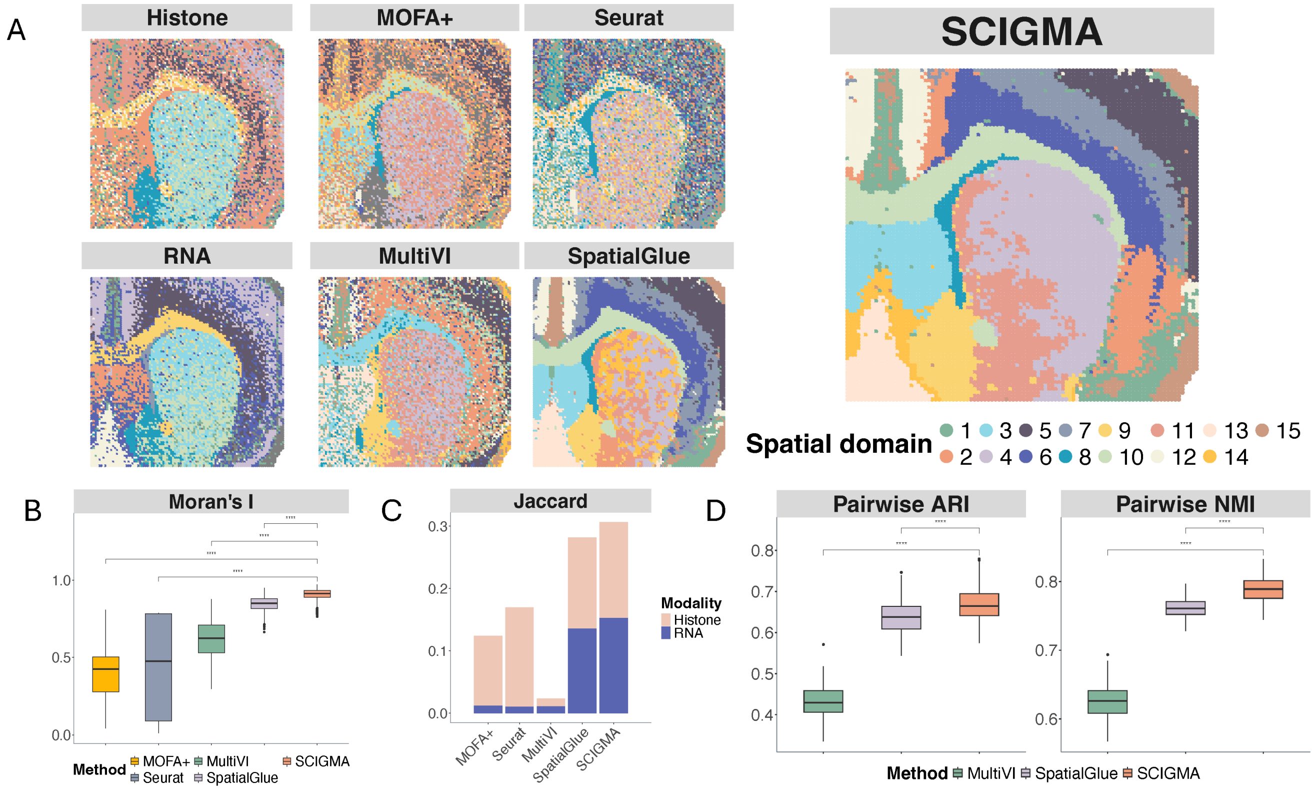

**Supplementary Figure 56. Analysis of the spatial epigenome-transcriptome (CUT&Tag-seq) H3K4me3 dataset.** (**A**) Spatial clusters identified by single-modality and multi-modal integration methods, including Seurat, MOFA+, MultiVI, SpatialGlue, and SCIGMA. (**B**) Boxplots displaying Moran's I values across methods, quantifying spatial autocorrelation. Each box spans the first to third quartiles, with the median represented by a horizontal line; whiskers extend to 1.5 times the interquartile range. (**C**) Jaccard similarity scores comparing preservation of modality-specific relationships in the joint representation across different methods. (**D**) Boxplots displaying pairwise ARI and NMI, assessing the consistency of clustering results across 20 random seeds for each method. SCIGMA successfully identified the cortex layers (clusters 5, 9, 2), CCG (cluster 4), VL (cluster 11), CP (cluster 5), ACB (cluster 14), and LSR (cluster 3) regions, showing improvements over all of the benchmarked methods. SCIGMA displayed the highest Moran's I and Jaccard similarity, and it showed the best consistency among the deep learning frameworks.

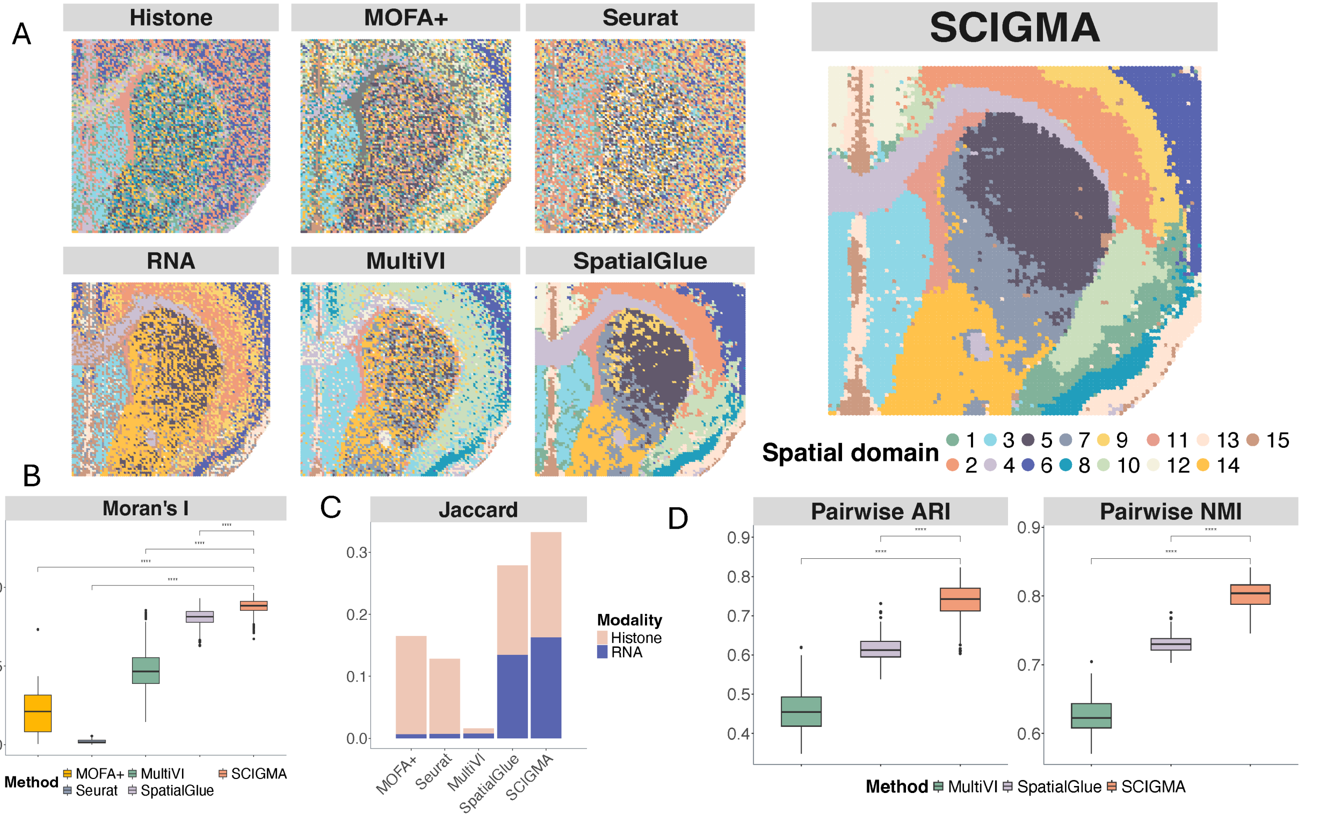

**Supplementary Figure 57. Analysis of the Stereo-CITE-seq mouse thymus dataset.** (**A)** Spatial clusters identified by single-modality and multi-modal integration methods, including Seurat, MOFA+, MultiVI, SpatialGlue, and SCIGMA. **(B)** Boxplots displaying Moran's I values across methods, quantifying spatial autocorrelation. Each box spans the first to third quartiles, with the median represented by a horizontal line; whiskers extend to 1.5 times the interquartile range. (**C**) Jaccard similarity scores comparing preservation of modality-specific relationships in the joint representation across different methods. (**D**) Boxplots displaying pairwise ARI and NMI, assessing the consistency of clustering results across 20 random seeds for each method. SCIGMA successfully identified spatial structures in the mouse thymus, uniquely detecting the capsule (cluster 6, marked by a dark blue arrow), the subcapsular cortex (cluster 10, marked by a green arrow), and blood vessels (cluster 3, marked by a blue arrow). These critical features were missed by other computational methods, highlighting SCIGMA's superior ability to resolve distinct anatomical regions.

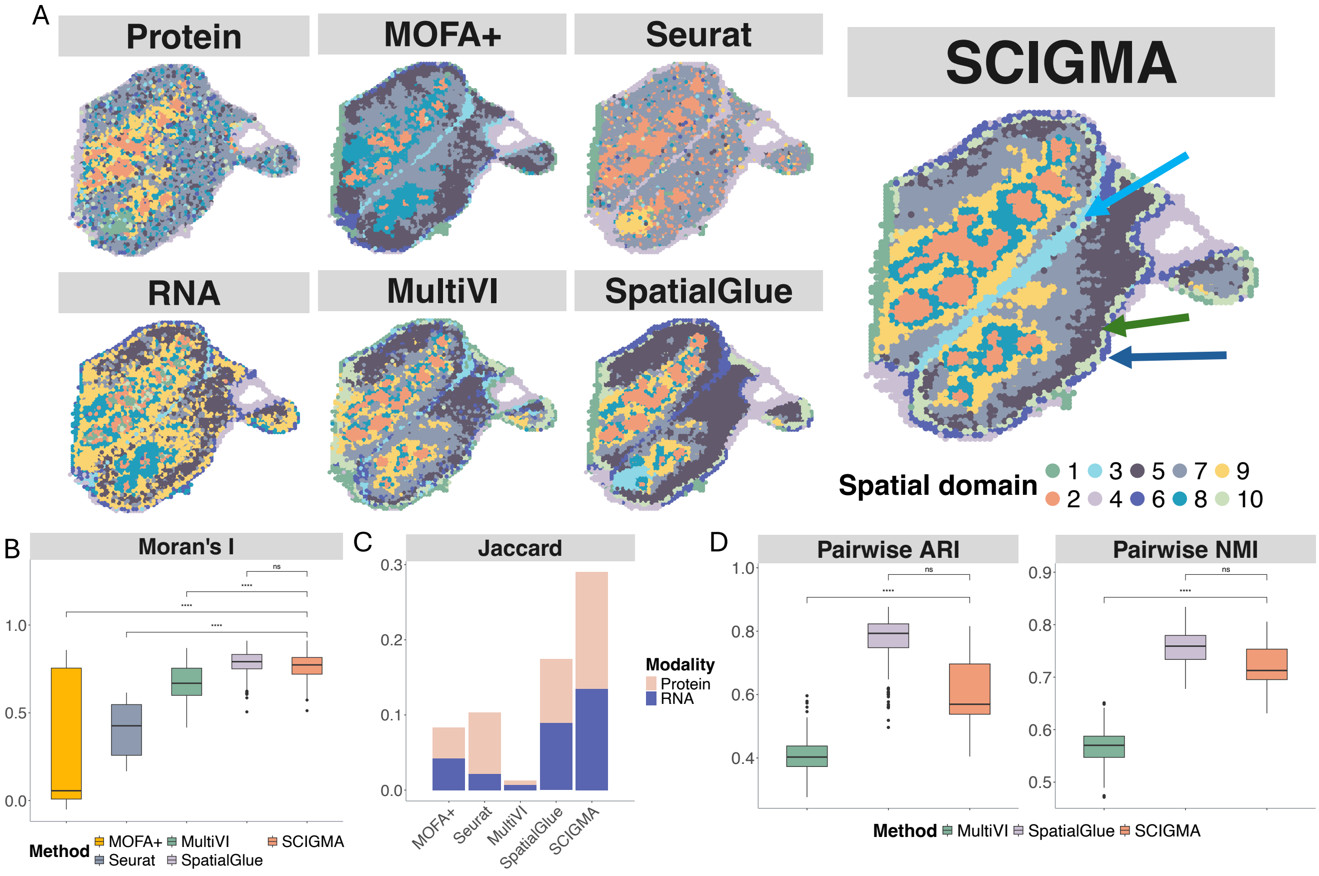

**Supplementary Figure 58. Analysis of the 10xXenium human breast cancer dataset. (A)** Spatial clusters identified by single-modality, and SCIGMA in the replicate 1 of the human breast cancer dataset **(B)** Spatial clusters identified by single-modality, and SCIGMA in the two replicates of the human breast cancer dataset. SCIGMA clearly identified distinct tumor subdomains, including the ductal carcinoma in situ (DCIS) domains that represent noninvasive forms of BC (cluster 1) and the invasive ductal carcinoma (IDC) domains (cluster 11 and 14), along with the other clusters such as immune-related regions (cluster 13) in the tumor microenvironment.

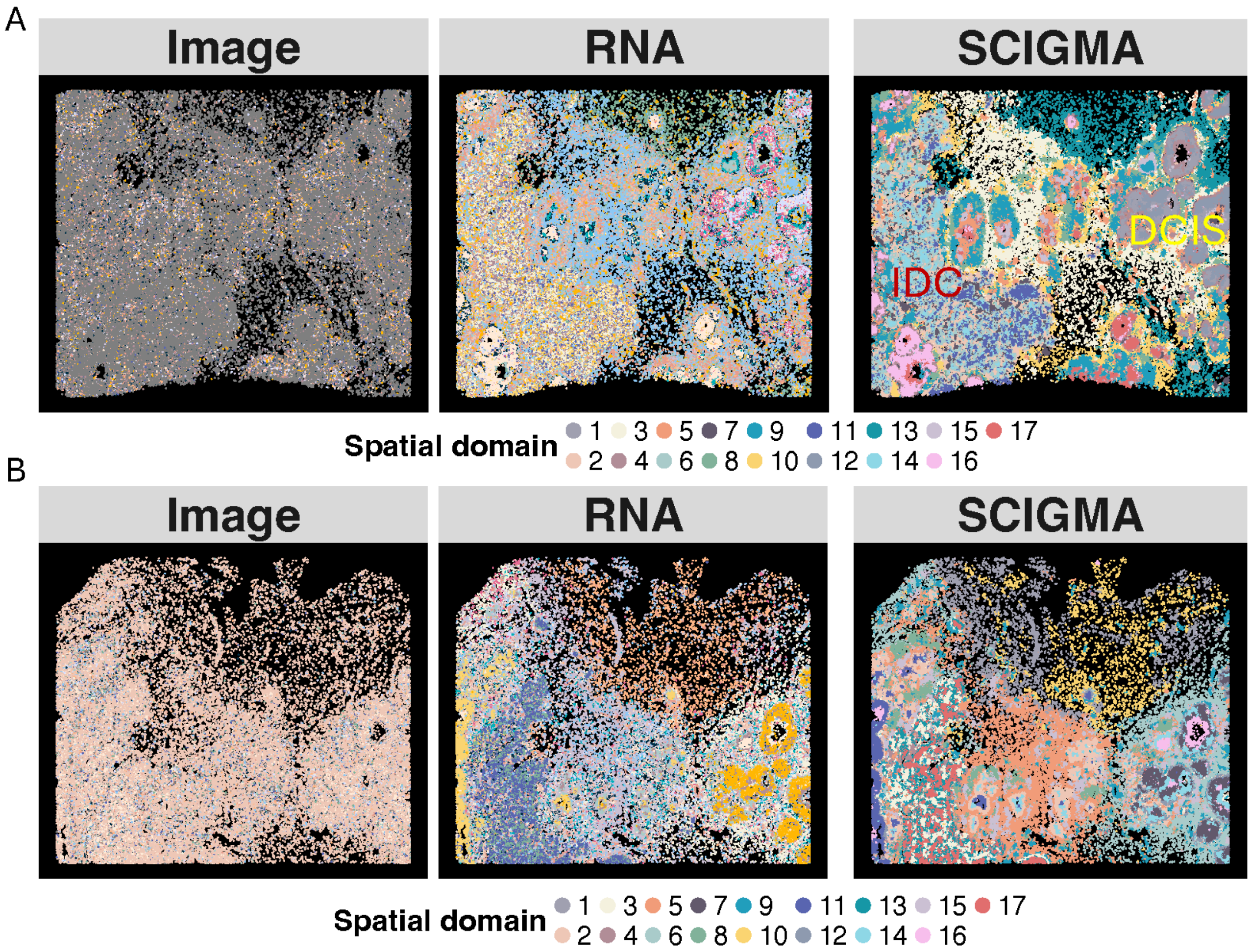

**Supplementary Figure 59. Analysis of the spatial metabolome-transcriptome dataset. Here, the plots display the** spatial clusters identified by single-modality, and SCIGMA in each dataset including (**A**) mouse brain striatum sample 1, (**B**) mouse brain striatum sample 2, (**C**) mouse brain striatum sample 3, (**D**) mouse brain subnigra sample 1 (**E**) mouse brain subnigra sample 2, (**F**) mouse brain subnigra sample 3. SCIGMA clearly identified distinct anatomical structures of the mouse brain, including clear striatum regions (marked by the red circles) and the substantia nigra (marked by the light blue circles) from the striatum and subnigra samples, respectively.

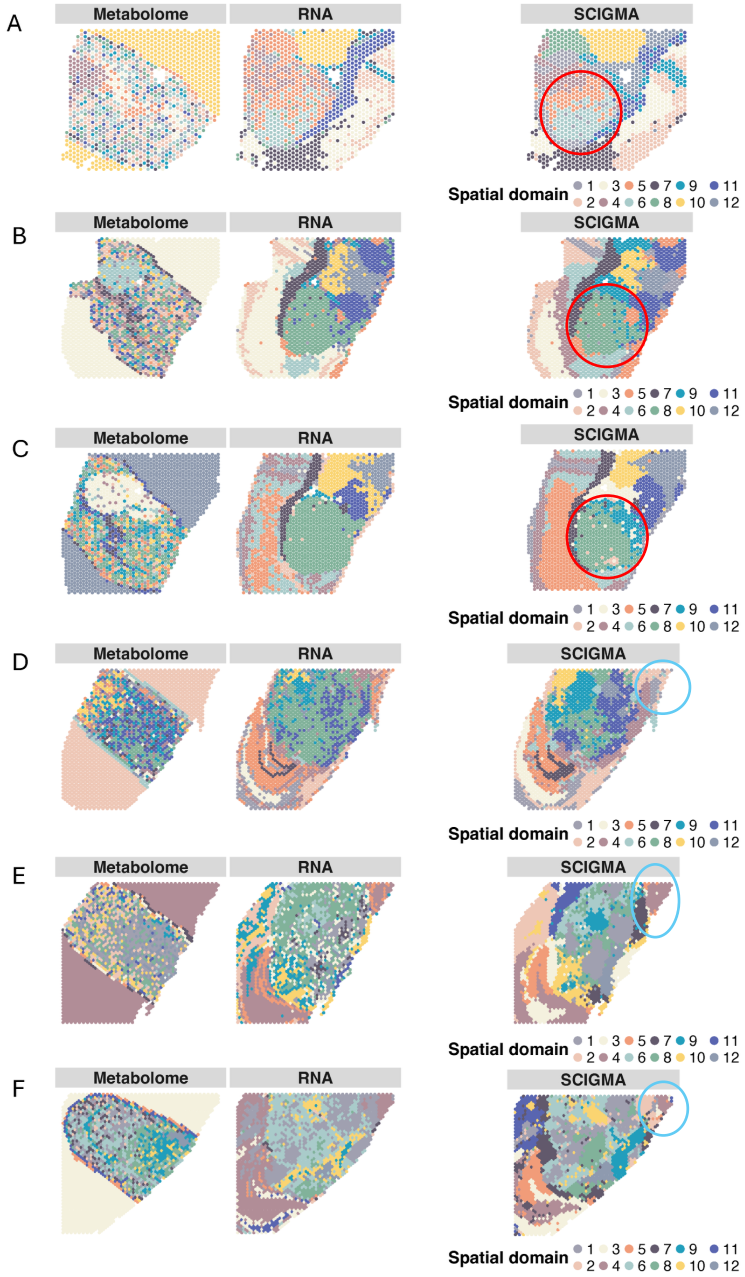

### Supplementary Tables

**Supplementary Table 1. List of datasets analyzed.** We analyze a variety of multi omics modalities across different types of tissues, including 19 datasets from 10 different tissues, 9 different technology platforms, and 8 different modalities, with number of spatial locations ranging from 2653 to 1.15 million. The table contains dataset name (column 1), technology/platform (column 2), modalities measured (column 3), dataset size (column 4). See Data Availability for downloading the corresponding datasets.

| Name | Platform | Modalities | # Spatial locations |
|---|---|---|---|
| P22 Mouse brain (RNA/ATAC) | Spatial-transcriptome-epigenome | RNA/ATAC | 9215 |
| Mouse spleen | SPOTS | RNA/Protein | 2653 |
| FF Human ovarian adenocarcinoma | 10x Xenium Prime 5K | RNA/Image | 1157659 |
| FFPE Human cervical cancer | | RNA/Image | 840397 |
| FFPE Mouse small intestine | 10x VisiumHD | RNA/Image | 748762 |
| FFPE Human colorectal cancer | | RNA/Image | 1052488 |
| 5-month Mouse brain (RNA/ATAC/H27K3me3/H27K3ac/Protein) | Spatial-Mux-Seq | RNA/ATAC/H27K3me3/H27K3ac/Protein | 10000 |
| P22 Mouse Brain RNA/H3K27ac | Spatial-transcriptome-epigenome | RNA/H3K27ac | 9370 |
| P22 Mouse Brain RNA/H3K27me3 | | RNA/H3K27me3 | 9752 |
| P22 Mouse Brain RNA/H3K4me3 | | RNA/H3K4me3 | 9548 |
| Mouse thymus | Stereo-CITE-seq | RNA/Protein | 4697 |
| Human breast cancer replicate 1 | 10x Xenium | RNA/Image | 167780 |
| Human breast cancer replicate 2 | | RNA/Image | 118752 |
| Mouse brain striatum sample 1 | | RNA/MSI | 1296 |

| | | | |
|---|---|---|---|
| Mouse brain striatum sample 2 | Spatial-transcriptome-metabolome | RNA/MSI | 1848 |
| Mouse brain striatum sample 3 | | RNA/MSI | 1791 |
| Mouse brain subnigra sample 1 | Spatial-transcriptome-metabolome | RNA/MSI | 1908 |
| Mouse brain subnigra sample 2 | | RNA/MSI | 2442 |
| Mouse brain subnigra sample 3 | | RNA/MSI | 2362 |

**Supplementary Table 2. Method applicability for each dataset.** We list the applicability of each method to the datasets analyzed. If a method 1) is designed for the measured modality types and 2) able to run on a cluster with at most 400gb of RAM, 24Gb of GPU memory, and 48 hours of runtime. We observe that SCIGMA is the only multi omics model that can handle all the available modalities including imaging modality obtained from existing platforms as well as datasets scaling into millions of spatial locations.

| Name | Platform | Seurat | MOFA+ | MultiVI | Spatial Glue | SCIGMA |
|---|---|---|---|---|---|---|
| P22 Mouse brain (RNA/ATAC) | Spatial-transcriptome-epigenome | ✓ | ✓ | ✓ | ✓ | ✓ |
| Mouse spleen | SPOTS | ✓ | ✓ | ✓ | ✓ | ✓ |
| FF Human ovarian adenocarcinoma | 10x Xenium Prime 5K | ✗ | ✗ | ✗ | ✗ | ✓ |
| FFPE Human cervical cancer | | ✗ | ✗ | ✗ | ✗ | ✓ |
| FFPE Mouse small intestine | 10x VisiumHD | ✗ | ✗ | ✗ | ✗ | ✓ |
| FFPE Human colorectal cancer | | ✗ | ✗ | ✗ | ✗ | ✓ |
| 5-month Mouse brain (RNA/ATAC/H27K3me3/H27K3ac/Protein) | Spatial-Mux-Seq | ✗ | ✗ | ✗ | ✗ | ✓ |
| P22 Mouse Brain RNA H3K27ac | Spatial-transcriptome-epigenome | ✓ | ✓ | ✓ | ✓ | ✓ |
| P22 Mouse Brain RNA H3K27me3 | | ✓ | ✓ | ✓ | ✓ | ✓ |
| P22 Mouse Brain RNA H3K4me3 | | ✓ | ✓ | ✓ | ✓ | ✓ |
| Mouse thymus | Stereo-CITE-seq | ✓ | ✓ | ✓ | ✓ | ✓ |
| Human breast cancer replicate 1 | 10x Xenium | ✗ | ✗ | ✗ | ✗ | ✓ |
| Human breast cancer replicate 2 | | ✗ | ✗ | ✗ | ✗ | ✓ |
| Mouse brain striatum sample 1 | Spatial-transcriptome-metabolome | ✗ | ✗ | ✗ | ✗ | ✓ |
| Mouse brain striatum sample 2 | | ✗ | ✗ | ✗ | ✗ | ✓ |
| Mouse brain striatum sample 3 | | ✗ | ✗ | ✗ | ✗ | ✓ |
| Mouse brain subnigra sample 1 | | ✗ | ✗ | ✗ | ✗ | ✓ |
| Mouse brain subnigra sample 2 | | ✗ | ✗ | ✗ | ✗ | ✓ |
| Mouse brain subnigra sample 3 | | ✗ | ✗ | ✗ | ✗ | ✓ |

**Supplementary Table 3. Quantitative metrics for the spatial ATAC-seq mouse brain and SPOTS mouse spleen datasets. (A)** Quantitative metric results for the spatial ATAC-seq mouse brain **(B)** Quantitative metric results SPOTS mouse spleen datasets for SCIGMA and the benchmarked models. Here, we assess the deep learning model's stability and consistency by calculating the pairwise adjusted Rand index (ARI) and Normalized Mutual Information (NMI) across 20 random seeds. We also calculated the Moran's I to assess the spatial autocorrelation of detected clusters, reflecting how well spatial structure is preserved, and Jaccard similarity coefficients to assess the consistency between the original modality-specific features and the model's learned joint representation. NA represents that the metric is not applicable to the corresponding method. For each metric, the value representing the best performance is highlighted in red.

**A.**

| Metric | Seurat | MOFA+ | MultiVI | SpatialGlue | SCIGMA |
|---|---|---|---|---|---|
| Median Pairwise ARI | NA | NA | 0.3329 | 0.6613 | 0.7102 |
| Median Pairwise NMI | NA | NA | 0.5267 | 0.7736 | 0.8105 |
| Median Moran's I | 0.7560 | 0.0373 | 0.6074 | 0.7368 | 0.8619 |
| Mean Jaccard index (RNA) | 0.0112 | 0.0112 | 0.00731 | 0.126 | 0.219 |
| Mean Jaccard index (ATAC) | 0.0230 | 0.0151 | 0.00759 | 0.151 | 0.227 |

**B.**

| Metric | Seurat | MOFA+ | MultiVI | SpatialGlue | SCIGMA |
|---|---|---|---|---|---|
| Median Pairwise ARI | NA | NA | 0.0133 | 0.4779 | 0.5877 |
| Median Pairwise NMI | NA | NA | 0.0274 | 0.5769 | 0.6686 |
| Median Moran's I | 0.6679 | 0.1651 | 0.3565 | 0.6187 | 0.6851 |
| Mean Jaccard index (RNA) | 0.0186 | 0.0326 | 0.0044 | 0.0718 | 0.123 |
| Mean Jaccard index (ATAC) | 0.0659 | 0.0557 | 0.0049 | 0.0895 | 0.144 |

**Supplementary Table 4. Datasets preprocessing details.** We list important parameters for preprocessing the datasets for input to SCIGMA. For each dataset (column 1), we list highly variable feature selection (column 2), dimension for PCA (column 3), number of spatial neighbors chosen for spatial graph construction (column 4), and number of feature neighbors chosen for feature graph construction (column 5). We use "/" as a delimiter between values for different modalities. NA represents no filtering of the features.

| Name | Highly variable features | PCA Dim | # Spatial Neighbors | # Feature Neighbors |
|---|---|---|---|---|
| P22 Mouse brain (RNA/ATAC) | 3000/3000 | 50/50 | 18 | 6 |
| Mouse spleen | 3000/NA | 20/20 | 6 | 6 |
| FF Human ovarian adenocarcinoma | 3000/NA | 30/30 | 18 | 6 |
| FFPE Human cervical cancer | 3000/NA | 30/30 | 18 | 6 |
| FFPE Mouse small intestine | 3000/NA | 40/40 | 18 | 6 |
| FFPE Human colorectal cancer | 3000/NA | 40/40 | 18 | 6 |
| 5-month Mouse brain (RNA/ATAC/H27K3me3/H27K3ac/Protein) | 3000/3000/3000/3000/NA | 30/30/30/30/30 | 18 | 6 |
| P22 Mouse Brain RNA H3K27ac | 3000/3000 | 50/50 | 18 | 6 |
| P22 Mouse Brain RNA H3K27me3 | 3000/3000 | 50/50 | 18 | 6 |
| P22 Mouse Brain RNA H3K4me3 | 3000/3000 | 50/50 | 18 | 6 |
| Mouse thymus | 3000/NA | 50/50 | 6 | 6 |
| Human breast cancer replicate 1 | 3000/NA | 30/30 | 18 | 6 |
| Human breast cancer replicate 2 | 3000/NA | 30/30 | 18 | 6 |
| Mouse brain striatum sample 1 | 3000/NA | 30/30 | 6 | 6 |
| Mouse brain striatum sample 2 | 3000/NA | 30/30 | 6 | 6 |
| Mouse brain striatum sample 3 | 3000/NA | 30/30 | 6 | 6 |
| Mouse brain subnigra sample 1 | 3000/NA | 30/30 | 6 | 6 |
| Mouse brain subnigra sample 2 | 3000/NA | 30/30 | 6 | 6 |
| Mouse brain subnigra sample 3 | 3000/NA | 30/30 | 6 | 6 |

**Supplementary Table 5. SCIGMA hyperparameters**. This table lists the hyperparameters for SCIGMA, optimized through a grid search. The hyperparameters include the latent dimension, which is the size of the joint representation used by SCIGMA; the batch size, representing the number of samples in each minibatch during training; the reconstruction weight and contrastive weight, which are the weights assigned to the reconstruction and contrastive loss terms, respectively; the number of epochs, indicating the total number of training epochs for SCIGMA; and the learning rate, which is the rate used by the optimizer.

| Name | Latent dimension | Batch size | Recon weight | Contrastive weight | Epoch | Learning Rate |
|---|---|---|---|---|---|---|
| P22 Mouse brain (RNA/ATAC) | 40 | 8000 | 1/1 | 1e-2/1e-2 | 600 | 1e-3 |
| Mouse spleen | 20 | 2000 | 1/1 | 1/1 | 300 | 1e-3 |
| FF Human ovarian adenocarcinoma | 20 | 8000 | 1/1 | 1/1 | 600 | 1e-3 |
| FFPE Human cervical cancer | 20 | 8000 | 1/1 | 1/1 | 600 | 1e-3 |
| FFPE Mouse small intestine | 36 | 8000 | 1/1e-1 | 1e-2/1e-2 | 400 | 1e-3 |
| FFPE Human colorectal cancer | 36 | 8000 | 1/1e-1 | 1e-2/1e-2 | 400 | 1e-3 |
| P22 Mouse brain (RNA/ATAC/H27K3me3/ H27K3ac/Protein) | 64 | 2000 | 1/1/1/1/1 | 1e-1/1e-2/1e-2/ 1e-2/1e-2 | 800 | 1e-3 |
| P22 Mouse Brain RNA H3K27ac | 40 | 8000 | 1/1 | 1e-2/1e-2 | 300 | 1e-3 |
| P22 Mouse Brain RNA H3K27me3 | 40 | 8000 | 1/1 | 1e-2/1e-2 | 300 | 1e-3 |
| P22 Mouse Brain RNA H3K4me3 | 40 | 8000 | 1/1 | 1e-2/1e-2 | 300 | 1e-3 |
| Mouse thymus | 20 | 2000 | 1/1 | 1/1 | 600 | 1e-3 |
| Human breast cancer replicate 1 | 20 | 8000 | 1/1e-1 | 1e-1/1e-2 | 300 | 1e-3 |
| Human breast cancer replicate 2 | 20 | 8000 | 1/1e-1 | 1e-2/1e-2 | 300 | 1e-3 |
| Mouse brain striatum sample 1 | 20 | All | 1/1e-1 | 1e-1/1e-2 | 500 | 1e-3 |
| Mouse brain striatum sample 2 | 20 | All | 1/1e-1 | 1e-1/1e-2 | 500 | 1e-3 |
| Mouse brain striatum sample 3 | 20 | All | 1/1e-1 | 1e-1/1e-2 | 500 | 1e-3 |
| Mouse brain subnigra sample 1 | 20 | All | 1/1e-1 | 1e-1/1e-2 | 500 | 1e-3 |
| Mouse brain subnigra sample 2 | 20 | All | 1/1e-1 | 1e-1/1e-2 | 500 | 1e-3 |
| Mouse brain subnigra sample 3 | 20 | All | 1/1e-1 | 1e-1/1e-2 | 500 | 1e-3 |

**Supplementary Table 6. SCIGMA training runtime for each dataset.** We list approximate training runtime for SCIGMA on each of the datasets. These benchmarks were conducted using a single GPU with 24GB of memory, a standard computational resource, demonstrating the efficiency of our method without requiring large-scale computing infrastructure.

| Name | Technology | # of Spatial locations | Runtime |
|---|---|---|---|
| P22 Mouse brain (RNA/ATAC) | Spatial-transcriptome-epigenome | 9196 | ~20min |
| Mouse Spleen | SPOTS | 2653 | ~2min |
| FF Human ovarian adenocarcinoma | 10x Xenium Prime 5K | 1144582 | ~120min |
| FFPE Human cervical cancer | | 480346 | ~120min |
| FFPE Mouse small intestine | 10x VisiumHD | 748272 | ~120min |
| FFPE Human colorectal cancer | | 1052488 | ~120min |
| P22 Mouse brain (RNA/ATAC/H27K3me3/H27K3ac/Protein) | Spatial-Mux-Seq | 9879 | ~11min |
| P22 Mouse Brain RNA H3K27ac | Spatial-transcriptome-epigenome | 9370 | ~12min |
| P22 Mouse Brain RNA H3K27me3 | | 9752 | ~12min |
| P22 Mouse Brain RNA H3K4me3 | | 9548 | ~12min |
| Mouse thymus | Stereo-CITE-seq | 4468 | ~5min |
| Human breast cancer replicate 1 | 10x Xenium | 123272 | ~25min |
| Human breast cancer replicate 2 | | 88803 | ~25min |
| Mouse brain striatum sample 1 | Spatial-transcriptome-metabolome | 1296 | ~3min |
| Mouse brain striatum sample 2 | | 1848 | ~3min |
| Mouse brain striatum sample 3 | | 1791 | ~3min |
| Mouse brain subnigra sample 1 | | 1908 | ~3min |
| Mouse brain subnigra sample 2 | | 2442 | ~3min |
| Mouse brain subnigra sample 3 | | 2362 | ~3min |

**Supplementary Table 7. XGBoost hyperparameters**. This table lists the hyperparameters for XGBoost models used for reconstruction, optimized through a grid search. The hyperparameters include max depth, representing the depth of the model; n_estimators, which is the number of epochs; eta, which is the learning rate; and early_stopping, or the early stopping criteria.

| Name | Max_depth | N_estimators | Eta | Early Stopping |
|---|---|---|---|---|
| P22 Mouse brain (RNA/ATAC) | 3 | 250 | 0.05 | 3 |
| Mouse spleen | 3 | 250 | 0.05 | 3 |
| FF Human ovarian adenocarcinoma | 8 | 250 | 0.05 | 3 |
| FFPE Human cervical cancer | 8 | 250 | 0.05 | 3 |
| FFPE Mouse small intestine | 8 | 250 | 0.05 | 3 |
| FFPE Human colorectal cancer | 8 | 250 | 0.05 | 3 |
| P22 Mouse brain (RNA/ATAC/H27K3me3/ H27K3ac/Protein) | 5 | 250 | 0.03 | 3 |

### Supplementary Notes

#### 1. SCIGMA Details

**1.1 Overview.** This section provides an overview of the notation used throughout the Supplementary Notes and a summary of the architecture of SCIGMA.

SCIGMA is a multi-view graph-based deep learning model optimized via an unsupervised contrastive learning framework. Given a spatial multi-modal dataset with $M$ different modalities, each with distinct feature set $\boldsymbol{X}_m \in R^{N\times p_m}$, where $N$ represents the number of spatial locations in the tissue, and $p_m$ denotes the number of features in modality $m \in \{1,2,\dots,M\}$. For each spatial location $i$, the modality-specific feature data is denoted as $\boldsymbol{X}_{im}$, and the corresponding 2-dimensional spatial coordinate matrix is denoted as $\boldsymbol{S}$, where $\boldsymbol{S} \in \mathrm{R}^{\mathrm{N}\times 2}$. For example, in spatial-ATAC-RNA-seq[1], $p_1$ and $p_2$ correspond to genes and chromatin regions, respectively, while in Spatial-CITE-seq[2] and 10x VisiumHD[3], they refer to genes co-profiled with proteins and H&E image pixels, respectively. Details of the multimodal measurements from the state-of-the-art spatial multi-modal technologies are listed in Supplementary Table 1.

SCIGMA constructs a combined graph of for each modality as $\boldsymbol{G}^{(c_m)} = \boldsymbol{G}^{(spa)} \cup \boldsymbol{G}^{(p_m)}$, where $\boldsymbol{G}^{(spa)}$ is the spatial graph constructed from the spatial coordinates and $\boldsymbol{G}^{(p_m)}$ is the feature graph constructed from the dimensionality-reduced feature matrices. From the constructed graph and dimensionality-reduced feature matrices $\boldsymbol{X}^{(m)}$, SCIGMA generates modality-specific embeddings $\boldsymbol{Y}^{(m)}$ using a graph attention encoder layer for each modality. Using cross-attention, SCIGMA fuses these embeddings into a joint representation $\boldsymbol{Z}$. The joint representation is passed through a decoder comprising a linear layer and modality-specific graph attention decoder layers to reconstruct the dimensionality-reduced features $\widehat{\boldsymbol{X}}^{(m)}$. Specifically, SCIGMA is optimized using a composite objective function that includes: (1) Reconstruction loss to preserve information (2) Contrastive learning loss to enhance modality alignment.

##### 1.2 CrossCLR – a variant of contrastive loss

For the second objective, we implement CrossCLR[4], a variant of contrastive loss. In standard contrastive learning, we define a paired sample from the output of two different encoder backbones as a 'positive' pair, and all other joint samples are defined as negative pairs. The contrastive loss function encourages similar samples to be closer while pushing dissimilar samples apart. The standard contrastive loss is formulated as:

$$L = -\log \frac{\delta(x_i, y_i)}{\delta(x_i, y_i) + \sum_{y_j \in N_i} \delta(x_i, y_j)} \tag{1}$$

Where $x_i$ and $y_i$ form a positive pair, meaning they are two views of the same sample, $N_i$ is the set of negative samples. $\delta(x_i, y_j)$ is a similarity function that quantifies how close two embeddings are in the shared space. Unlike standard contrastive learning, CrossCLR extends this framework by defining negative samples within and between different latent spaces[4]. This approach is particularly beneficial for integrating heterogeneous modalities such as RNA, protein, ATAC-seq, imaging, and etc. Specifically, in SCIGMA, CrossCLR is applied between the joint representation $Z$, which captures integrated information across all modalities, and the modality-specific representations $Y^{(m)}$, which retain the unique structure of each modality. By aligning the joint representation with each modality-specific representation separately, SCIGMA ensures that the joint embedding captures shared multi-omics information while allowing individual modalities to preserve their modality-specific characteristics. This design prevents excessive homogenization of embeddings across modalities, which could obscure meaningful modality-specific differences. For each spatial location $i$, let $Z_i$ denote the joint representation and $Y_i^{(m)}$ denote the modality-specific representation for modality $m$. The similarity between these embeddings is defined as:

$$\delta\left(Z_i, y_i^{(m)}\right) = \exp\left(\frac{Z_i^{\intercal} y_i^{(m)}}{\tau}\right) \tag{2}$$

Where $Z_i^{\intercal} y_i^{(m)}$ represents the dot product, capturing alignment between the vectors in the joint and modality-specific embedding space, and $\tau$ is a temperature hyperparameter controlling the sensitivity of the alignment. The exponential transformation ensures similarity values remain positive and enhances contrast between similar and dissimilar pairs. To optimize alignment between the joint and modality-specific representations, SCIGMA applies a separate CrossCLR loss to each modality. The total contrastive loss for modality $m$ is defined as:

$$\mathcal{L}_{contrast}^{(m)} = \gamma_m \frac{L(Z_i) + L\left(y_i^{(m)}\right)}{2} \tag{3}$$

where $\gamma_m$ is a hyperparameter that balances the contrastive alignment for modality $m$. The contrastive loss for the joint representation $Z_i$ is given by:

$$L(Z_i) = -\log \frac{\delta\left(Z_i, y_i^{(m)}\right)}{\delta\left(Z_i, y_i^{(m)}\right) + \sum_{y_j^{(m)} \in N_i^E} \delta\left(Z_i, y_j^{(m)}\right) + \sum_{Z_j \in N_i^R} \delta\left(Z_i, Z_j\right)} \tag{4}$$

Similarly, the contrastive loss for the modality-specific representation $y_i^{(m)}$ is defined as:

$$L\left(y_i^{(m)}\right) = -\log \frac{\delta\left(y_i^{(m)}, Z_i\right)}{\delta\left(y_i^{(m)}, Z_i\right) + \sum_{Z_j \in N_i^E} \delta\left(y_i^{(m)}, Z_j\right) + \sum_{y_j^{(m)} \in N_i^R} \delta\left(y_i^{(m)}, y_j^{(m)}\right)} \tag{5}$$

Here, $N_i^E$ represents the set of inter-modality negative samples, and $N_i^R$ represents the set of intra-modality negative samples. The denominator in both loss terms normalize the similarity scores, ensuring that positive pairs receive higher probabilities than negative pairs. A positive pair is defined as the joint and modality-specific latent representations corresponding to the same spatial location. Negative samples are defined as representations from all other spatial locations, both within the same modality (intra-modality) and across different modalities (inter-modality). Based on this setup, we define the overall loss function of SCIGMA when $m \in \{1,2\}$ as

$$\mathcal{L} = \mathcal{L}_{recon} + \mathcal{L}_{contrast}^{(1)} + \mathcal{L}_{contrast}^{(2)} \tag{6}$$

Equation (6) presents a representative loss function for integrating two modalities. SCIGMA's modeling framework extends naturally to more than two modalities by computing contrastive alignment between the joint embedding $Z_i$ and each modality-specific embedding $y_i^{(m)}$ without requiring explicit pairwise comparisons among all modalities. This enables SCIGMA to scale efficiently as the number of modalities increases. For example, SCIGMA has been successfully applied to Spatial-MUX-Seq[5] technology, which include modalities such as RNA, ATAC, histone, and protein. Details on the generalization to multiple modalities (more than two) are provided in the Supplementary Note 1.3 below. The ability to model multiple modalities while preserving the distinct structure of each further demonstrates the flexibility of the SCIGMA framework.

**1.3 Loss function for more than two modalities**. SCIGMA extends naturally to multi-omics datasets with more than two modalities by modifying both the architecture and loss function to accommodate additional modalities. Let $M$ denote the total number of modalities in the dataset. For each modality, SCIGMA employs an encoder backbone consisting of a series of Graph Attention Networks (GATs). As defined above, each encoder takes as input the preprocessed feature matrix $X^{(m)}$, the combined graph $G^{(c_m)}$, which is represented as both a sparse adjacency matrix and an edge list, and outputs a modality-specific latent representation, denoted as $\mathrm{Y}^{(\mathrm{m})}$. The loss function consists of two primary components: reconstruction loss and contrastive loss. First, the reconstruction loss is extended to account for the mean squared error (MSE) across all modalities:

$$\mathcal{L}_{recon} = \sum_{m=1}^{M} \beta_m \frac{1}{N} \sum_{i=1}^{N} \left\| x_i^{(m)} - \hat{x}_i^{(m)} \right\|_2^2 \tag{7}$$

Where $\beta_m$ is a weighting coefficient for each modality, and $\hat{x}_i^{(m)}$ represents the reconstructed features for modality $m$ at spatial location $i$. Similarly, the contrastive loss is extended to align the joint embedding $Z_i$ with each modality-specific embedding $\mathrm{Y}_\mathrm{i}^{(\mathrm{m})}$:

$$\mathcal{L}_{contrast} = \sum_{m=1}^{M} \mathcal{L}_{contrast}^{(m)} \tag{8}$$

Where $\mathcal{L}_{contrast}^{(m)}$ enforces alignment between the joint representation and each individual modality $m$. The overall loss function for SCIGMA is then formulated as:

$$\mathcal{L} = \mathcal{L}_{recon} + \mathcal{L}_{contrast} \tag{9}$$

SCIGMA is trained using the AdamW optimizer[6] to update the parameters of each modality encoder based on the loss function through backwards passes during training.

With the contrastive loss defined between modality specific and joint embeddings, the number of contrastive loss terms scales linearly rather than in polynomial time with respect to the number of modalities, as we do not need to take pairwise contrastive losses between each pair of modalities. This allows SCIGMA to naturally scale more efficiently to additional modalities. Even so, integration of more than two modalities remains challenging. Tuning a larger model requires finer hyperparameter tuning, the heterogeneity of more than two modalities creates more difficulty in finding a unified joint representation, and there is a non-negligible increase in computational burden. Despite these challenges, SCIGMA still performs admirably in integrating five modalities generated by the recent Spatial-MUX-Seq[5] technology; thus, we demonstrate the capabilities of SCIGMA's extension to more than two modalities, further improving confidence in the framework's scalability.

**1.4 Uncertainty modeling**. SCIGMA incorporates uncertainty estimation by dynamically learning the temperature parameter $\tau$, which is used to measure representation uncertainty for each spatial location. To achieve this, the joint representation obtained from the output of the cross-attention fusion layer is passed through a multilayer perceptron (MLP) that outputs a scalar value $\upsilon$ for each sample in the training batch. The final temperature value is computed as: $\tau = \exp(\upsilon)$. The exponentiation of $\upsilon$ ensures that $\tau$ remains strictly positive, preventing numerical instabilities such as division by zero or negative loss terms. Moreover, dynamic temperature scaling accentuates the distinction between certain and uncertain samples. When $\upsilon$ is small (indicating a confident sample), $\tau$ takes on a larger value, leading to a sharper similarity score and a more discriminative contrastive loss. Conversely, for uncertain samples, where $\upsilon$ is large, $\tau$ increases exponentially, resulting in a flatter similarity score and a smoother contrastive loss. This mechanism prevents the model from being overly influenced by high-uncertainty regions, leading to more stable optimization.

By dynamically adjusting similarity score sharpness based on sample confidence, SCIGMA prioritizes confident samples while mitigating the impact of uncertain ones. This prevents high-uncertainty regions from dominating the contrastive learning process, ensuring robust representation learning. The ability to scale effectively while maintaining a biologically meaningful latent space further demonstrates SCIGMA's potential as a powerful and interpretable framework for spatial transcriptomics and multi-modal data integration.

**1.5 Graph sampling**. To ensure computational efficiency while preserving the structural integrity of the combined graph, SCIGMA employs forest fire sampling[7], a technique that allows for efficient subgraph sampling. In this method, the sampling process begins with a randomly selected seed node, after which edges and adjacent nodes are iteratively "burned," meaning that each burned node propagates the sampling process to its neighboring nodes with a predefined probability. This process ensures that the sampled subgraph retains a representative structure of the larger graph. Given a set of combined graphs from the different modalities, SCIGMA cycles through different combined graphs at each training epoch and perform forest fire sampling with a specified batch size. Once a batch of nodes is selected, the remainder of the graph is restricted to only the sampled nodes. The same nodes and the resulting subgraphs are then sampled for the other modalities. By selecting an appropriate batch size (see Supplementary Table 5, Supplementary Note 1), SCIGMA can efficiently handle large-scale datasets containing millions of spatial locations, even on standard 24GB NVIDIA GPUs.

**1.6 Training.** SCIGMA takes as input the preprocessed, dimensionality-reduced feature matrices along with the constructed combined graphs. During training, graph sampling is performed as described above, allowing the model to process mini-batches efficiently. SCIGMA's parameters are optimized using stochastic gradient descent via the AdamW optimizer[6]. The use of gradient-based optimization ensures effective learning across both the contrastive and reconstruction loss components.

**1.7 Hyperparameter search.** Hyperparameters for training runs for each dataset are in Supplementary Table 5. A grid search was performed over the following hyperparameter ranges: number of spatial neighbors = [6, 12, 18], number of feature neighbors = [6, 12, 18], reconstruction loss weight = [1, 1e-1, 1e-2], contrastive loss weight = [1, 1e-1, 1e-2], batch size = [1000, 2000, 4000, 8000], latent space size = [16, 20, 24, 36, 40, 64], epoch = [300, 400, 500, 600, 800], learning rate = [1e-3, 1e-4]. For latent space size and batch size, hyperparameter values that exceeded the dataset's feature matrix size or sample size were removed from the search. In datasets with two modalities, a batch size of 8000 or lower allowed training on a single 24GB GPU without exceeding memory limitations. Minimizing the contrastive loss function completely can lead to representation collapse, where embedding features become indistinguishable. To avoid this, the hyperparameter search results were evaluated by periodically assessing clustering performance on the joint embedding rather than directly optimizing for downstream evaluation tasks. This approach ensures that SCIGMA's hyperparameter selection does not introduce information leakage or overfitting to predefined evaluation metrics while avoiding representation collapse, thus preserving the model's generalizability.

#### 2. Ablation studies

To systematically evaluate the contribution of individual components and design choices within SCIGMA, we conducted a series of ablation studies. These studies aim to quantify the impact of each component on the model's overall performance and integration capabilities. Specifically, we evaluated SCIGMA's modeling design from six distinct perspectives: (1) Contrastive loss: To demonstrate the necessity of contrastive loss for learning well-integrated joint representations, we trained SCIGMA without the contrastive loss component. This helped assess how critical this element is for aligning different modalities within the integrated framework. (2) Spatial Information: To highlight the importance of spatial information, we trained SCIGMA using only feature graph. This variation allowed us to understand the extent to which spatial relations contribute to model accuracy and data integration. (3) Feature graph: To highlight the importance of the feature graph, we trained SCIGMA using only spatial graph, testing whether modality-specific feature similarities further enhance integration (4) Graph attention layers: To demonstrate the benefits of using graph attention layers, we substituted GAT with GCN. This change helped to evaluate the impact of different graph modeling architectures on the model's ability to learn from complex structures. (5) CrossCLR loss: To quantify the importance of the CrossCLR loss, we trained SCIGMA with a regular contrastive loss objective, assessing its role in aligning modality-specific and joint representations (6) Uncertainty parameter: Finally, to underscore the importance of learning the uncertainty parameter for training stability and effective integration of heterogeneous data, we trained SCIGMA using vanilla CrossCLR contrastive loss (without the trained uncertainty parameter). This experiment demonstrates that incorporating an uncertainty parameter improves model stability and enhances multimodal integration. These ablation experiments were conducted on the Spatial ATAC-seq dataset, where the biological structure of the P22 mouse brain is well-defined. We evaluated each model variant from different perspectives, including clustering performance, Moran's I score, Jaccard index, Dirichlet energy, pairwise ARI/NMI, and the RMSE of SCIGMA's modality-specific reconstruction error. Details of calculating these metrics see Supplementary Notes 5.1 – 5.5.

Specifically, the clustering results (Figure S1A) revealed that models trained without contrastive loss or without the spatial graph failed to produce well-separated and biologically meaningful spatial clusters, indicating poor integration of modality-specific information. In the vanilla contrastive loss setting (the regular contrastive loss), although some domain structures were correctly identified, clear inaccuracies were observed in distinguishing the VL, ACB, and LPO regions, highlighting the importance of SCIGMA's CrossCLR design choice. Conversely, models trained without the feature graph or with GCN instead of GAT produced overly smooth clusters, reinforcing the importance of the feature graph for capturing heterogeneous structures (such as the heterogenous tumor data) and exposing the tendency of GCN to over smooth representations. Comparing the full SCIGMA model to the variant without uncertainty learning, we observed small yet meaningful improvements in the cortical layers, further underscoring the importance of uncertainty score. While the absence of spatial information impaired spatial domain detection, excessive reliance on spatial neighbors and naïve graph averaging resulted in oversmoothed domain predictions, reinforcing the necessity of the combined graph and GAT architecture.

From the Moran's I score (Figure S1B), we observed that spatial-only SCIGMA configuration (SCIGMA no feature graph) achieved the highest scores. This result is expected, as both Moran's I and Jaccard similarity place significant weight on spatial continuity. In contrast, the feature-graph-only SCIGMA model (SCIGMA No Spatial Graph) exhibited markedly lower spatial autocorrelation, suggesting that excluding spatial information hampers the model's ability to capture spatial patterns. Models trained without contrastive loss or with vanilla/regular contrastive loss performed comparably to the full model in terms of Moran's I score but showed reduced Jaccard similarity (Figure S1C), suggesting that the CrossCLR loss improves the joint representation's alignment. The performance of the GCN-based and no-uncertainty variants was similar, indicating that the spatial graph and CrossCLR framework play primary roles in learning structured latent representations. To further quantify over-smoothing effects, we calculated the Dirichlet energy[8] metric (Figure S1D). The metric has been used to evaluate over smoothness in GNNs, as it measures the difference in features between neighboring nodes. Higher values indicate less smoothness in the embedding, while lower values indicate more smoothness. From the Dirichlet energy metric, we see that the no feature graph and GCN ablation models have lower Dirichlet energy, while the no spatial graph ablation model has the highest Dirichlet energy. This suggests potential over-smoothing in the joint representations the no feature graph and GCN ablation models, while the no spatial graph ablation model may not be sufficiently incorporating neighbor information.

The pairwise ARI and NMI results (Figure S1E) demonstrated that the full SCIGMA model consistently outperformed all ablation variants, indicating that the combined graph, GAT layers, and uncertainty parameter learning collectively contribute to model stability across different initialization seeds, which is an increasing area of importance for deep learning models in the biomedical field.

The RMSE of the reconstructed feature matrix (Figure S1F) provided additional insights into how different model components influence alignment at uncertain locations. The feature-graph-only SCIGMA model achieved the lowest RMSE, reinforcing the importance of the feature graph in capturing global feature relationships. The full SCIGMA model ranked second, demonstrating that uncertainty parameter learning improves the model's ability to attend to misaligned regions. In contrast, the GCN-based SCIGMA model exhibited the highest RMSE, further highlighting the over smoothing problem and reinforcing the choice of GAT layers in the full SCIGMA model.

Overall, our ablation evaluations indicate that each design choice in SCIGMA is essential for achieving optimal performance. Spatial graph information is critical for capturing biological spatial structures, as evidenced by the Moran's I and Jaccard similarity results. The feature graph and graph attention layers are crucial for learning well-integrated representations while avoiding over smoothing, as supported by the clustering experiments and the Dirichlet energy metric. Finally, uncertainty parameter learning plays a key role in maintaining training stability, as demonstrated by the pairwise ARI and NMI comparisons. These findings validate SCIGMA's architectural choices and illustrate that each component contributes to its effectiveness in spatial multi-omics data integration.

In addition to these ablation studies, SCIGMA has been extensively evaluated across 19 datasets spanning 10 different tissues, 8 distinct technology platforms, and 10 different modalities, with the number of spatial locations ranging from 2,653 to 1.15 million. Across this diverse range of datasets, SCIGMA consistently demonstrates strong performance, further supporting the robustness and generalizability of its design choices.

**3. Datasets**

*Spatial ATAC-RNA--seq P22 Mouse Brain*[1]. We obtained the anndata containing counts data and corresponding spatial data from SpatialGlue's data release https://zenodo.org/records/10362607. This dataset measured 9215 spatial locations with 22914 genes and 121068 peaks. Using scanpy, we filtered spatial locations with less than 200 total gene counts, and filtered genes and peaks that were expressed in under 10 cells, and took the intersection of shared spatial locations across all modalities. This resulted in 9196 remaining spatial locations. We selected the top 3000 highly variable genes and 3000 highly variable peaks using scanpy.pp.highly_variable_genes. For the gene expression, we performed log-normalization with a target sum of 1e6 using scanpy.pp.normalize_total and scanpy.pp.log1p. We then scaled the data using scanpy.pp.scale and performed PCA using scanpy.pp.pca with 50 principal components. For the peak matrix, we performed LSI normalization to 50 components. We used the spatial coordinates to construct the spatial graph with 18 neighbors using sklearn.neighbors.NearestNeighbors. We used the processed features matrices to generate the feature graph using sklearn.neighbors.kneighbors_graph with 6 neighbors. We obtained the combined graph by taking the graph union of the spatial and feature graphs and binarizing the resulting adjacency graph.

*SPOTS Mouse Spleen*[9]*.* We obtained the anndata containing counts data and corresponding spatial data from SpatialGlue's data release https://zenodo.org/records/10362607. This dataset measured 2653 spatial locations with 33285 genes and 21 proteins. Using scanpy, we filtered genes that were expressed in under 10 cells. We selected the top 3000 highly variable genes with scanpy.pp.highly_variable_genes. For the gene expression, we performed log-normalization with a target sum of 1e6 using scanpy.pp.normalize_total and scanpy.pp.log1p. We then scaled the data using scanpy.pp.scale and performed PCA using scanpy.pp.pca with 20 principal components. For the protein expression matrix, we performed CLR normalization. We then scaled the data using scanpy.pp.scale and performed PCA using scanpy.pp.pca with 20 principal components. We used the spatial coordinates to construct the spatial graph with 6 neighbors using sklearn.neighbors.NearestNeighbors. We used the processed features matrices to generate the feature graph using sklearn.neighbors.kneighbors_graph with 6 neighbors. We obtained the combined graph by taking the graph union of the spatial and feature graphs and binarizing the resulting adjacency graph.

*10x Xenium Prime 5K FF Human Ovarian Adenocarcinoma*[10]*.* We obtained the gene counts matrix and morphology image with the corresponding spatial data from 10x Genomics, https://www.10xgenomics.com/datasets/xenium-prime-fresh-frozen-human-ovary. This dataset measured 1157659 spatial locations with 5001 genes with a corresponding morphology tif image. Using scanpy, we filtered spatial locations with less than 50 total gene counts and genes that were expressed in under 10 cells, and took the intersection of shared spatial locations across all modalities. This resulted in 1144582 remaining spatial locations. We selected the top 3000 highly variable genes with scanpy.pp.highly_variable_genes. For the gene expression, we performed log-normalization with a target sum of 1e6 using scanpy.pp.normalize_total and scanpy.pp.log1p. We then scaled the data using scanpy.pp.scale and performed PCA using scanpy.pp.pca with 30 principal components. For morphology image, we first aligned the spatial coordinates by converting the coordinate locations used by the gene expression to pixel dimensions. Each of the channels were normalized to values between 0 and 1. For each spatial location, we extracted a tile from the morphology image of dimension 64 by 64 pixels and passed through a pretrained ResNet18 model[11] from torchvision.models. We then scaled the ResNet embeddings using scanpy.pp.scale and performed PCA using sklearn.decomposition. PCA with 30 principal components. We used the spatial coordinates to construct the spatial graph with 18 neighbors using sklearn.neighbors.NearestNeighbors. We used the processed features matrices to generate the feature graph using sklearn.neighbors.kneighbors_graph with 6 neighbors. We obtained the combined graph by taking the graph union of the spatial and feature graphs and binarizing the resulting adjacency graph.

*10x Xenium Prime 5K FFPE Human Cervical Cancer[10].* We obtained the gene counts matrix and morphology image with the corresponding spatial data from 10x Genomics, https://www.10xgenomics.com/datasets/xenium-prime-ffpe-human-cervical-cancer. This dataset measured 840397 spatial locations with 5101 genes with a corresponding morphology tif image. Using scanpy, we filtered spatial locations with less than 50 total gene counts and genes that were expressed in under 10 cells, and took the intersection of shared spatial locations across all modalities. This resulted in 480346 remaining spatial locations. We selected the top 3000 highly variable genes with scanpy.pp.highly_variable_genes. For the gene expression, we performed log-normalization with a target sum of 1e6 using scanpy.pp.normalize_total and scanpy.pp.log1p. We then scaled the data using scanpy.pp.scale and performed PCA using scanpy.pp.pca with 30 principal components. For morphology image, we first aligned the spatial coordinates by converting the coordinate locations used by the gene expression to pixel dimensions. Each of the channels were normalized to values between 0 and 1. For each spatial location, we extracted a tile from the morphology image of dimension 64 by 64 pixels and passed through a pretrained ResNet18[11] model from torchvision.models. We then scaled the ResNet embeddings using scanpy.pp.scale and performed PCA using sklearn.decomposition. PCA with 30 principal components. We used the spatial coordinates to construct the spatial graph with 18 neighbors using sklearn.neighbors.NearestNeighbors. We used the processed features matrices to generate the feature graph using sklearn.neighbors.kneighbors_graph with 6 neighbors. We obtained the combined graph by taking the graph union of the spatial and feature graphs and binarizing the resulting adjacency graph.

*10x Visium HD FFPE Human Colorectal Cancer[10].* We obtained the gene counts matrix and morphology image with the corresponding spatial data from 10x Genomics, https://www.10xgenomics.com/datasets/visium-hd-cytassist-gene-expression-libraries-of-human-crc. This dataset measured 1052488 spatial locations with 17239 genes with a corresponding morphology tif image. Using scanpy, we filtered spatial locations with less than 50 total gene counts and genes that were expressed in under 10 cells, and took the intersection of shared spatial locations across all modalities. This resulted in 1052488 remaining spatial locations. We selected the top 3000 highly variable genes with scanpy.pp.highly_variable_genes. For the gene expression, we performed log-normalization with a target sum of 1e6 using scanpy.pp.normalize_total and scanpy.pp.log1p. We then scaled the data using scanpy.pp.scale and performed PCA using scanpy.pp.pca with 40 principal components. For morphology image, we first aligned the spatial coordinates by converting the coordinate locations used by the gene expression to pixel dimensions. Each of the channels were normalized to values between 0 and 1. For each spatial location, we extracted a tile from the morphology image of dimension 64 by 64 pixels and passed through a pretrained ResNet18[11] model from torchvision.models. We then scaled the ResNet embeddings using scanpy.pp.scale and performed PCA using sklearn.decomposition. PCA with 40 principal components. We used the spatial coordinates to construct the spatial graph with 18 neighbors using sklearn.neighbors.NearestNeighbors. We used the processed features matrices to generate the feature graph using sklearn.neighbors.kneighbors_graph with 6 neighbors. We obtained the combined graph by taking the graph union of the spatial and feature graphs and binarizing the resulting adjacency graph.

*10x Visium HD FFPE Mouse Small Intestine*[10]*.* We obtained the gene counts matrix and morphology image with the corresponding spatial data from 10x Genomics, https://www.10xgenomics.com/datasets/visium-hd-cytassist-gene-expression-libraries-of-mouse-intestine. This dataset measured 748762 spatial locations with 16024 genes with a corresponding morphology tif image. Using scanpy, we filtered spatial locations with less than 50 total gene counts and genes that were expressed in under 10 cells, and took the intersection of shared spatial locations across all modalities. This resulted in 748272 remaining spatial locations. We selected the top 3000 highly variable genes with scanpy.pp.highly_variable_genes. For the gene expression, we performed log-normalization with a target sum of 1e6 using scanpy.pp.normalize_total and scanpy.pp.log1p. We then scaled the data using scanpy.pp.scale and performed PCA using scanpy.pp.pca with 40 principal components. For morphology image, we first aligned the spatial coordinates by converting the coordinate locations used by the gene expression to pixel dimensions. Each of the channels were normalized to values between 0 and 1. For each spatial location, we extracted a tile from the morphology image of dimension 64 by 64 pixels and passed through a pretrained ResNet18[11] model from torchvision.models. We then scaled the ResNet embeddings using scanpy.pp.scale and performed PCA using sklearn.decomposition. PCA with 40 principal components. We used the spatial coordinates to construct the spatial graph with 18 neighbors using sklearn.neighbors.NearestNeighbors. We used the processed features matrices to generate the feature graph using sklearn.neighbors.kneighbors_graph with 6 neighbors. We obtained the combined graph by taking the graph union of the spatial and feature graphs and binarizing the resulting adjacency graph.

*Spatial Mux Seq 5 Month Mouse Brain*[5]. We obtained the 5-month mouse brain five modality dataset from Gene Expression Omnibus accession no. GSE263333. From the provided RNA feature matrix, barcodes, and spatial coordinates, we spatially resolved the RNA expression data. From the provided protein feature matrix, barcodes, and spatial coordinates, we spatially resolved the protein expression data. From the fragments files provided for ATAC, H3K27ac, and H3K27me3, we used ArchR to generate peak matrices. We first created arrow files using the parameters: minFrags = 0, maxFrags = 1e+05, addGeneScoreMat = True, and ‘the mm10’ genome. We computed the dimensionality reduced space via IterativeLSI with dims = 1:30. We generated peak sets using 'addReproduciblePeakSet' and Macs2. We then called addPeakMatrix to generate a peak matrix for the three epigenomics modalities. This resulted in 10000 spatial locations and 48400 measured genes, 131 measured proteins, 12208 measured ATAC peaks, 16226 measured H3K27ac peaks, and 1468 measured H3K27me3 peaks. Using the RNA matrix, we filtered spatial locations with less than 80 total counts and genes that were expressed in under 10 cells with scanpy and took the intersection of shared spatial locations across all modalities. This resulted in 9879 remaining spatial locations. We selected the top 3000 highly variable genes with scanpy.pp.highly_variable_genes. We performed log-normalization with a target sum of 1e6 using scanpy.pp.normalize_total and scanpy.pp.log1p. We then scaled the data using scanpy.pp.scale and performed PCA using scanpy.pp.pca with 30 principal components. For ATAC/H3K27ac/H3K27me3, we selected the top 3000 highly variable genes with scanpy.pp.highly_variable_genes. We performed LSI normalization to 30 components. For the protein expression matrix, we performed CLR normalization. We then scaled the data using scanpy.pp.scale and performed PCA using scanpy.pp.pca with 30 principal components. We used the spatial coordinates to construct the spatial graph with 18 neighbors using sklearn.neighbors.NearestNeighbors. We used the processed features matrices to generate the feature graph using sklearn.neighbors.kneighbors_graph with 6 neighbors. We obtained the combined graph by taking the graph union of the spatial and feature graphs and binarizing the resulting adjacency graph.

*Spatial CUT&Tag-seq P22 Mouse Brain*[1]. We obtained the anndata containing counts data and corresponding spatial data from SpatialGlue's data release https://zenodo.org/records/10362607. This dataset measured 9370~9752 spatial locations with 18373~20231 genes and 23993~24029 peaks. Using scanpy, we filtered genes that were expressed in under 10 cells. We selected the top 3000 highly variable genes with scanpy.pp.highly_variable_genes. For the gene expression, we performed log-normalization with a target sum of 1e6 using scanpy.pp.normalize_total and scanpy.pp.log1p. We then scaled the data using scanpy.pp.scale and performed PCA using scanpy.pp.pca with 40 principal components. For the peak matrix, we performed LSI normalization with 40 principal components. We used the spatial coordinates to construct the spatial graph with 18 neighbors using sklearn.neighbors.NearestNeighbors. We used the processed features matrices to generate the feature graph using sklearn.neighbors.kneighbors_graph with 6 neighbors. We obtained the combined graph by taking the graph union of the spatial and feature graphs and binarizing the resulting adjacency graph.

*Stereo-CITE-Seq Mouse Thymus*[12]. We obtained the anndata containing counts data and corresponding spatial data from SpatialGlue's data release https://zenodo.org/records/10362607. This dataset measured 4697 spatial locations with 23622 genes and 51 proteins. Using scanpy, we filtered spatial locations with less than 80 total gene counts and filtered genes that were expressed in under 10 cells, and took the intersection of shared spatial locations across all modalities. This resulted in 4468 remaining spatial locations.

We selected the top 3000 highly variable genes with scanpy.pp.highly_variable_genes. For the gene expression, we performed log-normalization with a target sum of 1e6 using scanpy.pp.normalize_total and scanpy.pp.log1p. We then scaled the data using scanpy.pp.scale and performed PCA using scanpy.pp.pca with 50 principal components. For the protein expression matrix, we performed CLR normalization. We then scaled the data using scanpy.pp.scale and performed PCA using scanpy.pp.pca with 50 principal components. We used the spatial coordinates to construct the spatial graph with 6 neighbors using sklearn.neighbors.NearestNeighbors. We used the processed features matrices to generate the feature graph using sklearn.neighbors.kneighbors_graph with 6 neighbors. We obtained the combined graph by taking the graph union of the spatial and feature graphs and binarizing the resulting adjacency graph.

*10x Xenium Breast Cancer*[10]. We obtained the gene counts matrix and morphology image with the corresponding spatial data from 10x Genomics, https://www.10xgenomics.com/products/xenium-in-situ/human-breast-dataset-explorer. This dataset consists of two replicates from the same patient sample, which measured 167780/118752 spatial locations with 313/313 genes with a corresponding morphology tif image. Using scanpy, we filtered spatial locations with less than 100 total gene counts and genes that were expressed in under 10 cells, and took the intersection of shared spatial locations across all modalities. This resulted in 123273/88803 remaining spatial locations. We selected the top 3000 highly variable genes with scanpy.pp.highly_variable_genes. For the gene expression, we performed log-normalization with a target sum of 1e6 using scanpy.pp.normalize_total and scanpy.pp.log1p. We then scaled the data using scanpy.pp.scale and performed PCA using scanpy.pp.pca with 30 principal components. For morphology image, we first aligned the spatial coordinates by converting the coordinate locations used by the gene expression to pixel dimensions. Each of the channels were normalized to values between 0 and 1. For each spatial location, we extracted a tile from the morphology image of dimension 64 by 64 pixels and passed through a pretrained ResNet18[11] model from torchvision.models. We then scaled the ResNet embeddings using scanpy.pp.scale and performed PCA using sklearn.decomposition.PCA with 30 principal components. We used the spatial coordinates to construct the spatial graph with 18 neighbors using sklearn.neighbors.NearestNeighbors. We used the processed features matrices to generate the feature graph using sklearn.neighbors.kneighbors_graph with 6 neighbors. We obtained the combined graph by taking the graph union of the spatial and feature graphs and binarizing the resulting adjacency graph.

*Spatial Metabolome-Transcriptome Mouse Brain*[13]*.* We obtained the anndata containing counts data and corresponding spatial data from Mendeley Data https://data.mendeley.com/datasets/w7nw4km7xd/1. This dataset measured 1791~2442 spatial locations with 32285 genes and 1538 neurotransmitters. Using scanpy, we filtered genes that were expressed in under 10 cells. We selected the top 3000 highly variable genes with scanpy.pp.highly_variable_genes. For the gene expression, we performed log-normalization with a target sum of 1e6 using scanpy.pp.normalize_total and scanpy.pp.log1p. We then scaled the data using scanpy.pp.scale and performed PCA using scanpy.pp.pca with 30 principal components. For the protein expression matrix, we performed CLR normalization. We then scaled the data using scanpy.pp.scale and performed PCA using scanpy.pp.pca with 30 principal components. We used the spatial coordinates to construct the spatial graph with 6 neighbors using sklearn.neighbors.NearestNeighbors. We used the processed features matrices to generate the feature graph using sklearn.neighbors.kneighbors_graph with 6 neighbors. We obtained the combined graph by taking the graph union of the spatial and feature graphs and binarizing the resulting adjacency graph.

**4. Benchmark models: details on implementations/training of the benchmarks**

*Seurat.* We benchmarked Seurat following their Github tutorial with default parameters (https://satijalab.org/seurat/). We then extracted the representation "cell.embeddings" from the joint representations for downstream analysis.

*MOFA+.* We benchmarked MOFA+ following their Github tutorial with default parameters (https://biofam.github.io/MOFA2/). We then extracted the factors from the "mofa" object for downstream analysis.

*MultiVI.* We benchmarked MultiVI[14] following their Github tutorial with default parameters (https://docs.scvi-tools.org/en/stable/user_guide/models/multivi.html). We then extracted the representation 'X_MultiVI' for downstream tasks.

*SpatialGlue.* We benchmarked SpatialGlue[15] using their default dataset preprocessing and hyperparameters, following their Github tutorial (https://spatialglue-tutorials.readthedocs.io/en/latest/). We then extracted the joint representation for downstream tasks.

**5. Downstream Analyses**

**5.1 Clustering**. SCIGMA learns a joint representation from the expression data of different modalities. We then followed spatialGlue's paper to apply the mclust[16] algorithm to the learned joint representation to identify spatial domains. For each dataset, we used the number of clusters determined from the analyses as a starting point (if available) to test different numbers of clusters to select the clustering that best captured biological structures, and treated cluster number as a hyperparameter in grid search for model selection (Supplementary Note 1). We used the R implementation of mclust.

**5.2 Pairwise ARI/NMI.** To assess the stability of SCIGMA's clustering results across different training runs, we computed the Adjusted Rand Index (ARI) and Normalized Mutual Information (NMI) between cluster assignments obtained from multiple random seeds. These metrics quantify the consistency of cluster partitions and their agreement with known biological structures. The Rand Index (RI) measures the similarity between two clustering assignments by considering how many pairs of elements are grouped consistently across both partitions. Given two sets of assigned labels, $\mathrm{N}$ and $\mathrm{M}$, let $\mathrm{k}$ be the number of element pairs that are in the same cluster in both $\mathrm{N}$ and $\mathrm{M}$, and let $\mathrm{l}$ be the number of element pairs that are in different clusters in both $\mathrm{N}$ and $\mathrm{M}$. The Rand Index is then given by:

$$\mathrm{RI} = \frac{\mathrm{k} + \mathrm{l}}{\#\ \text{all possible pairs}} \tag{14}$$

The Adjusted Rand Index (ARI) corrects the Rand Index by accounting for the expected similarity under random labeling. This adjustment ensures that an ARI score close to zero corresponds to random clustering, while a score close to one indicates nearly identical clustering assignments:

$$\mathrm{ARI} = \frac{\mathrm{RI} - \mathrm{E}[\mathrm{RI}]}{\max(\mathrm{RI}) - \mathrm{E}[\mathrm{RI}]} \tag{15}$$

To further evaluate clustering agreement between multiple training runs, we computed the Mutual Information (MI), which measures how much information is shared between two cluster assignments. Given a clustering assignment $\mathrm{N}$, the entropy of $\mathrm{N}$ is defined as:

$$\mathrm{H}(\mathrm{N}) = -\sum_{\mathrm{i}=1}^{|\mathrm{N}|} \mathrm{P}(\mathrm{i})\log\big(\mathrm{P}(\mathrm{i})\big) \tag{16}$$

where $\mathrm{P}(\mathrm{i})$ is the probability that a randomly chosen element belongs to cluster $\mathrm{N_i}$. Similarly, let $\mathrm{P}(\mathrm{j})$ be the corresponding probability for clusters in assignment $\mathrm{M}$. The MI is defined as

$$\mathrm{MI}(\mathrm{N},\mathrm{M}) = \sum_{\mathrm{i}=1}^{|\mathrm{N}|}\sum_{\mathrm{i}=1}^{|\mathrm{M}|} \mathrm{P}(\mathrm{i},\mathrm{j})\log\left(\frac{\mathrm{P}(\mathrm{i},\mathrm{j})}{\mathrm{P}(\mathrm{i})\mathrm{P}'(\mathrm{j})}\right) \tag{17}$$

where $\mathrm{P}(\mathrm{i},\mathrm{j})$ represents the joint probability of an element being assigned to cluster $\mathrm{i}$ in $\mathrm{N}$ and cluster $\mathrm{j}$ in $\mathrm{M}$. The NMI is computed as:

$$\mathrm{NMI}(\mathrm{N},\mathrm{M}) = \frac{\mathrm{MI}(\mathrm{N},\mathrm{m})}{\mathrm{mean}\big(\mathrm{H}(\mathrm{N}),\mathrm{H}(\mathrm{M})\big)} \tag{18}$$

We used the ARI and NMI between clustering results from different seeds of different SCIGMA training runs as a measure of training stability.

**5.3 Moran's I**. Moran's I was computed to evaluate the spatial autocorrelation of SCIGMA's joint representation. Given feature values and their corresponding spatial locations, Moran's I quantifies whether the spatial pattern is clustered, dispersed, or random. The metric is defined as:

$$I = \frac{N}{W}\frac{\sum_i^N \sum_i^N w_{i,j} z_i z_j}{\sum_i^n z_i^2} \tag{19}$$

where $z_i = x_i - \tilde{X}$ represents the deviation of feature $x_i$ from the mean, $w_{i,j}$ is the spatial weight between locations $i$ and $j$, $N$ is the total number of spatial locations, and $W$ is the sum of all spatial weights $w_{i,j}$. A positive Moran's I value indicates spatial clustering, whereas a negative value suggests dispersion. We computed Moran's I using Scanpy's morans_i function, where the SCIGMA embeddings were used as features, and spatial weights were derived from the spatial coordinates.

**5.4 Jaccard similarity.** To assess the alignment between the original modality feature space and the learned joint representation, we computed the Jaccard similarity between their respective nearest-neighbor sets for each spatial location. Given a spatial location $i$, we defined its neighborhood in two ways: (1) Original feature space neighborhood ($N_{original}^{i}$): The 10 nearest spatial neighbors and 10 nearest feature-space neighbors. (2) Joint representation neighborhood ($N_{joint}^{i}$): The 20 nearest neighbors in the joint embedding space. The Jaccard similarity at spatial location $i$ is the defined as

$$J_i = \frac{|N_{original}^{i} \cap N_{joint}^{i}|}{|N_{original}^{i} \cup N_{joint}^{i}|} \tag{20}$$

where a larger Jaccard similarity value indicates greater consistency between the original modality-specific feature space and the learned joint representation. By comparing these neighborhoods, we assess how well SCIGMA preserves local relationships while integrating multi-omics data into a unified embedding space.

**5.5 Dirichlet metric calculation**. To evaluate the smoothness of the joint representations in the ablated SCIGMA models, we computed the Dirichlet energy metric, a commonly used measure of over-smoothing in graph neural networks (GNNs)[8]. Given an embedding $Z$ and a graph $G = (V, E)$, the Dirichlet energy is calculated as:

$$\epsilon(Z) = \frac{1}{|E|} \sum_{i \in V} \sum_{j \in N_i} \left|\left|Z_i - Z_j\right|\right|_2^2$$

where $N_i$ denotes the neighborhood of node $i$, and the metric is normalized by the number of edges $|E|$ to account for variations in graph structure across ablation models. Lower Dirichlet energy indicates greater similarity between neighboring nodes, reflecting a smoother representation, whereas higher Dirichlet energy signifies greater differences between neighbors, indicating less smoothness and greater retention of distinct features.

**5.6 Neighborhood enrichment analysis.** To assess the spatial relationships between clusters, we calculated neighborhood enrichment scores with the Squidpy[17] package. We first obtained clusters as described above using the joint representation from SCIGMA. To find neighborhood enrichment to evaluate spatial autocorrelations of clusters, we used the identified clusters and spatial coordinates as input to Squidpy's neighborhood enrichment score function, nhood_enrichment. The function calculates the following: for a pair of clusters $i$ and $j$, it first counts the number of nodes that belong to those two classes and are proximal to each other ($x_{ij}$). Then, it permutes the cluster labels while keeping the connectivities fixed to estimate mean

$\mu_{ij}$ and standard deviation $\sigma_{ij}$ of random cluster labelings. Then, it calculates a z-score for each cluster pair $i$ and $j$:

$$Z_{ij}^{nhood} = \frac{x_{ij} - \mu_{ij}}{\sigma_{ij}} \tag{21}$$

The z-score shows if a pair of clusters are over or under represented in cluster-to-cluster node interactions in the spatial connectivity graph. Thus, it shows spatial relationships between pairs of clusters.

**5.7 Differential gene and protein expression analysis.** On the 'SCT' assay, we first performed log-normalization and data scaling using the Seurat package. To find differentially expressed genes, we used the 'FindAllMarkers' function with the following recommended parameter settings: logfc.threshold = 0.1, min.pct = 0.1 and 'wilcoxon' test for all datasets except for the SPOTS and VisiumHD datasets. As the VisiumHD datasets were especially sparse and the SPOTS dataset had few samples, we used the following parameter settings for 'FindAllMarkers': logfc.threshold = 0.0, min.pct = 0.0 and 'wilcoxon' test. The differentially expressed genes or proteins were defined as the those with adjusted pvalues < 0.05, and the top 10 differentially expressed markers were determined by ranking their logfoldchange from the largest to the smallest.

**5.8 Differentially expressed peak analysis.** To find differentially expressed peaks, we used the the ArchR package (v1.0.2)[18]. We first created arrow files using the parameters: minFrags = 0, maxFrags = 1e+07, tile_size = 5000, addGeneScoreMat = True, and 'the mm10' genome. We computed the dimensionality reduced space via IterativeLSI with dims = 1:30. We defined the peak set found in the anndatas if the dataset provided them, or generated them using 'addReproduciblePeakSet'. The differentially expressed peaks were then identified for SCIGMA's clusters in the 'PeakMatrix' with the 'getMarkerFeatures' function. Marker genes with differential gene scores were also computed from the 'GeneScoreMatrix' using the same function. Finally, we computed the linkage between genes and peaks using the 'addPeak2GeneLinks' function with the 'Iterative LSI' reductions and 'GeneScoreMatrix' values. We ran the 'addPeak2GeneLinks' function with the following settings: corCutOff = 0.45 and resolution = 1,000. To visualize the correlation between peaks and genes, we used the 'plotPeak2GeneHeatmap' function to plot the peak-to-gene links heatmap. The differentially expressed peaks were defined as the those with adjusted pvalues < 0.05, and the top 10 differentially expressed markers were determined by ranking their logfoldchange from the largest to the smallest.

**5.9 Gene set enrichment analysis**. We performed gene set enrichment analysis[19] on the differentially expressed genes. We used the python library 'gseapy' and the corresponding GSEA function 'prerank' to run the analysis. For the mouse datasets, we used the MSigDB Mouse curated gene sets, and for the human datasets, we used the MSigDB Human curated gene sets[20]. To rank the genes, we performed raw p-value cutoff by setting unadjusted p-values to 1e-200 for any unadjusted p-values lower than 1e-200, and ranked them according to the following formula: sign(average logfoldchange) * abs(Normal_PPF(unadjusted p-value * 0.5)). For the two modality mouse datasets, the following parameters were used for 'prerank': min_size = 5, max_size = 500, permutation_num=1000. For the two modality human datasets, the following parameters were used for 'prerank': min_size = 5, max_size = 300, permutation_num=1000. For the five-modality mouse dataset, the following parameters were used for 'prerank': min_size = 2, max_size = 1000, permutation_num=1000. The resulting GSEA outputs were filtered with the following to identify upregulated processes: for the mouse datasets, NES >=0 and FDR <= 0.25; for the human datasets, NES >= 1 and FDR <= 0.25.

**5.10 PAGA analysis.** We ran PAGA using the 'paga' function from the scanpy library. The analysis began with the construction of a k-nearest neighbor graph using 'scanpy.pp.neighbors', followed by the computation of the trajectory structure using 'scanpy.tl.paga'. The resulting PAGA graph was then used as the initial positioning framework for generating a low-dimensional UMAP embedding via 'scanpy.tl.umap'. This approach allows for the visualization of spatially and biologically relevant trajectories based on the learned joint representation.

**5.11 XGBoost reconstruction analysis.** To reconstruct the modality specific data, such as the gene expression, protein expression, and chromatin accessibility, we performed a reconstruction analysis by regressing SCIGMA's reconstructed PCA embeddings back to the original feature space using an XGBoost[21] regressor. To do so, given the reconstructed PCA feature matrices, the original PCA feature matrices, and the original feature matrices, we trained the regressor using the XGBoost Python library. The training and validation datasets were generated by splitting the original PCA feature matrices and their corresponding raw feature matrices using sklearn.model_selection.train_test_split. To optimize model performance, we conducted a hyperparameter search over the following parameters: max_depth = {3,5,8}, n_estimators = {50,100,150,250}, eta = {0.03, 0.05}, and early_stopping = {3,5}. After training the regressor, we applied it to the reconstructed PCA feature matrices from SCIGMA to generate reconstructions in the original feature space.

**5.10 Uncertainty analysis.** We provide details for the implementation of each of the uncertainty analyses. Specifically, to interpret the uncertainty estimates provided by SCIGMA's model, we conducted four independent evaluations: spatial distribution of uncertainty estimates, expression difference analysis, RMSE analysis, and Jaccard similarity analysis. We first checked the spatial distribution of uncertainty estimates by visualization to examine which region has a high uncertainty. Then we performed the following quantification analysis:

To evaluate expression differences in certain and uncertain regions, we computed the absolute difference between the log-normalized and scaled preprocessed feature matrices for the corresponding modalities. Uncertain regions were defined by ranking the uncertainty estimates from SCIGMA and selecting locations in the top $k$% (where $k$ = 1, 5, 10, 20) as uncertain, with the remaining locations classified as certain. We then compared the distribution of absolute expression differences between uncertain and certain regions using a one-sided Wilcoxon rank-sum test to assess whether uncertainty arises from differences between modalities.

For the RMSE analysis, we computed location-wise RMSE between the original PCA feature matrix and the reconstructed PCA feature matrix. Uncertain regions were identified using the same thresholding approach as in expression difference analysis. We then compared RMSE distributions between uncertain and certain regions using a one-sided Wilcoxon rank-sum test.

For Jaccard similarity analysis, we computed the Jaccard similarity between modality-specific embeddings and joint embeddings. Uncertain regions were identified using the same thresholding approach as in expression difference analysis. We then compared Jaccard similarity distributions between uncertain and certain regions for each modality using a one-sided Wilcoxon rank-sum test.

21 Chen, T. & Guestrin, C. in *Proceedings of the 22nd acm sigkdd international conference on knowledge discovery and data mining.* 785-794.
